# Human skeletal muscle CD90^+^ fibro-adipogenic progenitors are associated with muscle degeneration in type 2 diabetic patients

**DOI:** 10.1101/2020.08.25.243907

**Authors:** Jean Farup, Jesper Just, Frank de Paoli, Lin Lin, Jonas Brorson Jensen, Tine Billeskov, Ines Sanchez Roman, Cagla Cömert, Andreas Buch Møller, Luca Madaro, Elena Groppa, Rikard Göran Fred, Ulla Kampmann, Steen B. Pedersen, Peter Bross, Tinna Stevnsner, Nikolaj Eldrup, Tune H. Pers, Fabio M. V. Rossi, Pier Lorenzo Puri, Niels Jessen

## Abstract

Aging and type 2 diabetes mellitus (T2DM) are associated with impaired skeletal muscle function and degeneration of the skeletal muscle microenvironment. However, the origin and mechanisms underlying the degeneration are not well described in human skeletal muscle. Here we show that skeletal muscles of T2DM patients exhibit pathological degenerative remodeling of the extracellular matrix that was associated with a selective increase of a subpopulation of fibro-adipogenic progenitors (FAPs) marked by expression of *THY1* (CD90) - the FAP^CD90+^. We identified Platelet-derived growth factor (PDGF) signaling as key regulator of human FAP biology, as it promotes proliferation and collagen production at the expense of adipogenesis, an effect accompanied with a metabolic shift towards glycolytic lactate fermentation. FAPs^CD90+^ showed a PDGF-mimetic phenotype, with high proliferative activity and clonogenicity, increased production of extracellular matrix production and enhanced glycolysis. Importantly, the pathogenic phenotype of T2DM FAP^CD90+^ was reduced by treatment with the anti-diabetic drug Metformin. These data identify PDGF-driven conversion of a sub-population of FAPs as a key event in the pathogenic accumulation of extracellular matrix in T2DM muscles.

## INTRODUCTION

Skeletal muscle is a major organ comprising ~40% of human body mass, accountable for ~30% of basic metabolic rate^1^ and solely responsible for generating mechanical forces to enable breathing and locomotion. In addition, skeletal muscle is a major storage organ for glucose (glycogen) and the primary target for peripheral insulin stimulated glucose uptake^2^. With the importance of skeletal muscle for overall body health, it is of no surprise that degeneration of skeletal muscle and loss of muscle mass is associated with a multitude of different diseases and ultimately shortened life span^3^. Degeneration of skeletal muscle has often been investigated in models of severe genetic diseases such as Duchenne’s muscular dystrophy (DMD) or severe trauma induced paralysis e.g. in spinal cord injury. Under such circumstances, the parenchymal muscle cells are progressively replaced by non-contractile tissue including extracellular matrix (ECM) proteins and adipocytes, resulting in loss of contractile and metabolic function. Aging and age-related diseases are strongly associated with degeneration of human skeletal muscle, in particular evidenced as muscle atrophy. In contrast to the aggressive degenerative progression of muscle environment in DMD or following trauma, muscle degeneration associated to aging and age-related diseases, such as type 2 diabetes mellitus (T2DM), develop gradually and is accompanied by muscle atrophy. Ectopic deposition of non-parenchymal tissue, such as fibrotic scars and fat in a number of different organs and tissues, including liver, kidney, fat and skeletal muscle is a common pathogenic feature of many age-associated diseases, ultimately leading to multi-organ failure and death^4^. In T2DM, which can be viewed as an accelerated aging process ^5,6^, an excessive deposition of ectopic adipocytes has been observed in the viscera, the pancreas and skeletal muscle^7–10^. Pre-clinical rodent models of insulin resistance and T2DM show accumulation of adipocytes and ECM proteins in skeletal muscle that results in an impairment of metabolic and contractile muscle functions^11–13^. Importantly, similar features have been reported in insulin resistant subjects or patients affected by diabetic polyneuropathy ^8,10^. Understanding the mechanism driving such pathological muscle remodeling is fundamental to provide interventions toward preventing muscle atrophy and fibrosis as well as metabolic alterations in T2DM patients.

In order to understand and potentially reduce or prevent the degeneration of skeletal muscle, it is of key importance to determine the cellular origin of fibrosis and adipocytes. In mice, skeletal muscle resident cells with fibrogenic and adipogenic potential (i.e. fibro-adipogenic progenitors; FAPs) were identified a decade ago by two independent reports as stem cell antigen 1^+^, CD34^+^, Platelet Derived Growth Factor Receptor α (PDGFRα)^+^ cells^14,15^. Notably, several reports have shown that FAPs are cellular targets of therapeutic interventions toward reducing or preventing fibrosis and adipocyte accumulation during in mouse models of regeneration, disease or aging ^16–21^. In particular, the tyrosine kinase receptor and highly specific FAP marker, PDGFRα, has been shown to play a key role in FAP activation and induction of tissue fibrosis in several organs including adipose tissue and skeletal muscle ^18,22,23^.

In models of T2DM, aberrant FAP activity has been shown to underlie fatty degeneration of the diaphragm muscle during a high fat diet intervention, which ultimately impaired mechanical muscle function ^11^.

Unfortunately, the translation of these studies into human skeletal muscle has been challenged by the lack of *in vivo* human FAP markers and the general scarcity of human skeletal muscle tissue biopsies. However, studies utilizing cultured cells from muscle homogenates suggest that similar cells are present in human muscle^24–26^, indicating that targeting these cells may represent a novel way of preventing tissue degeneration in T2DM.

Here we used FACS-mediated isolation and transcriptomic profiling at population and single-cell level (scRNA-seq) of FAPs from human T2DM patient-derived biopsies to identify a subpopulation of FAPs, defined by their expression of *THY1* (CD90), as putative cellular source of the fibro-fatty infiltration of muscles from T2DM patients and potential target of therapeutic interventions with anti-diabetic agents.

## RESULTS

### T2DM is associated with degenerative remodeling of the extracellular matrix

To investigate if T2DM is associated with fibro-fatty degeneration of skeletal muscle, we collected muscle biopsies from age and overweight-matched subjects (OBS), T2DM patients (T2D) and insulin treated T2DM patients (itT2D) with severe insulin resistance (treated with 196±26 IU insulin/day)^27,28^. This allowed us to investigate the potential degeneration of the skeletal muscle environment as a consequence T2DM and the severity of insulin resistance. These subjects/patients have been extensively described previously in relation to insulin sensitivity (hyperinsulimic euglycemic clamp and glucose tracer), c-peptide, HbA^1c^ and fasting glucose ^28,29^. To perform an unbiased evaluation of major transcriptional alterations, RNA was isolated from crude muscle tissue, subjected to RNA-sequencing ^27^ and followed by differential gene expression (DE) and pathway enrichment analysis (Fig 1 A). In addition to a dysregulated muscle morphology, contraction and inflammation, our analysis revealed that DE genes were substantially enriched in biological processes and pathways related to ECM turnover and remodeling in itT2D patients (Fig 1 B, C, Fig S1 A, B). This was particularly evident as the severity of insulin resistance increased (OBS < T2D < itT2D), which was validated by qPCR of common genes associated with fibrosis (Fig 1 D, E, F, G). Downregulated genes in itT2D patients were enriched in pathways related to insulin signaling, glucose response and circadian rhythm, consistent with the insulin resistance profile of these patients (Fig S1 C). We extended our analysis to sections of skeletal muscle biopsies obtained from T2DM and non-diabetic patients undergoing coronary artery bypass surgery, by performing immunohistochemical staining’s that showed an increased deposition of Collagen-1 in T2DM patients, as compared to non-diabetics (Fig 1 H, I). Interestingly, we also observed Perilipin-1^+^ adipocytes in the interstitial space in the T2DM patients, which was not evident in control biopsies (Fig 1 J). Collectively, our findings support the hypothesis that T2DM is associated with a degeneration of skeletal muscle evidenced here as increased ECM content and potential adipocyte accumulation.

**Fig 1.**
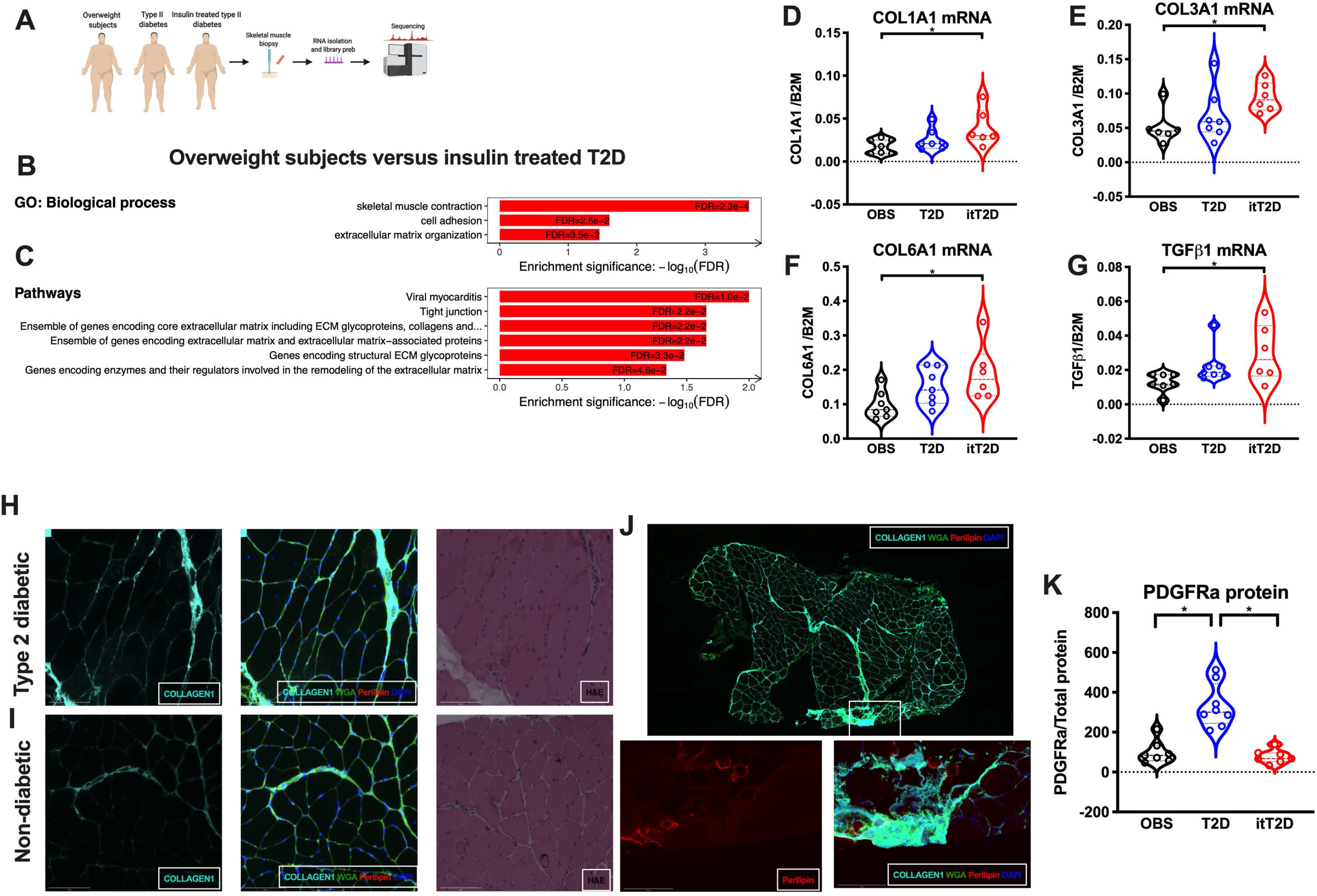
Type 2 diabetes is associated with degenerative remodeling of the extra-cellular matrix. **A**, Schematic presentation of the sample work-flow collected from overweight subjects (OBS, n=7), type II diabetic patients (T2D, n=7) and insulin treated type 2 diabetes (itT2D, n=6); **B**, Biological processes increased in itT2D compared to OBS; **C**, Enriched pathways in itT2D compared to OBS; **D**, *COL1A1* **E**, *COL3A1*; **F**, *COL6A1*; **G**, *TGFB1* mRNA expression in OBS, T2D and itT2D; **H+I**, Immunohistochemical (IHC) staining of Collagen-1 (Cyan), Wheat Germ Agglutinin (WGA, green), Perilipin 1 (Red) and DAPI (Blue) in on muscle sections from patients with type 2 diabetes (**H**) and without type 2 diabetes (**I**); **J**, IHC of muscle sections from type 2 diabetes displaying adipocytes in the interstitial space; **K**, PDGFRα protein expression in skeletal muscle from OBS, T2D, itT2D;. Significant difference denoted by *p<0.05.

### Human skeletal muscle FAPs can be prospectively isolated as CD34^+^CD56^−^CD45^−^CD31^−^

In mice, skeletal muscle fibrosis and fat accumulation is thought to primarily originate from FAPs, which have been shown to specifically express PDGFRα^14,15,30^. To examine if FAPs could be the cellular source of fibrosis and adipocytes in T2DM, we quantified protein content of PDGFRα in whole muscle homogenate. We found a specific increase in the T2D patients (Fig 1 K, S1 D), which declined to control (OBS) levels in itT2D. When stromal cells differentiate into mature myofibroblasts or adipocytes they downregulate PDGFRα expression, which likely explains its lower expression in the itT2D group^31^.

To further understand the cellular mechanism underlying the degeneration of the skeletal muscles in T2DM patients, we set out to isolate FAPs from human skeletal muscle by FACS. We initially attempted to isolate human FAPs freshly from skeletal muscle, using PDGFRα as a specific positive marker and negative selection for CD56 (muscle stem cells; MuSCs), CD31 (endothelial cells) and CD45 (hematopoietic cells). However, we obtained no clear signal or cell population from any of the PDGFRα antibodies tested (data not shown). We therefore tested alternative mesenchymal stem cell and FAP markers previously used to isolate mouse FAPs, including ALCAM (CD166), CD34 and THY1 (CD90) ^15,32^. While no signal was obtained from CD166 (not shown), a population of CD34^+^CD56^−^CD45^−^CD31^−^ cells (Fig 2 A i, S2 A i-vii) could be isolated, which could be further subdivided into a CD34^+^CD90^−^ and a CD34^+^CD90^+^ population (Fig 2 A ii, S2 A i-vii). For comparison, we also identified MuSCs based on expression of CD56 as well as CD82 (Fig 2 A iii, S2 A i-vii). Endothelial cells and hematopoietic cells were initially selected on CD31/CD45 expression and could then be discriminated based on CD34 expression by endothelial cells (Fig S2 iv; Fig S2 B i-ii)^33^. For sorting, we included Propidium Iodide and fluorescence minus one controls were used to set the gates (Fig S2 C i-iv).

**Fig 2.**
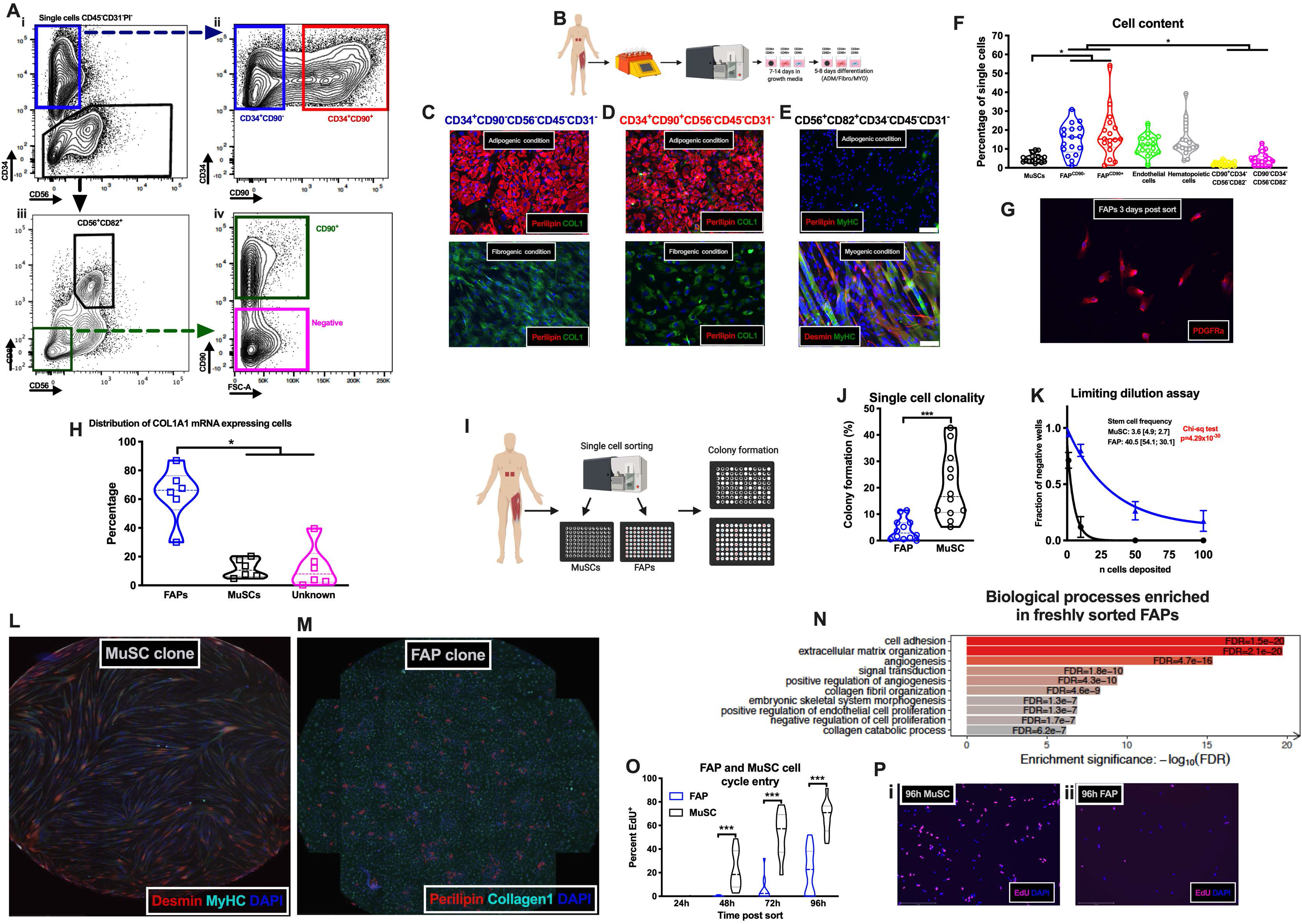
Identification and characterization of human fibro-adipogenic progenitors. **A-i**, Contour flow-plots from the sorting strategy identifying cell negative for CD45, CD31 and Propidium Iodide (PI) and either CD34^+^CD56^−^ or CD34^−^; **A-ii**, displaying a CD34^+^CD90^−^ and a CD34^+^CD90^+^ population; **A-iii**, displaying a CD56^+^CD82^+^ and a CD56^−^CD82^−^ population; **A-iv**, displaying a CD90^+^ and a CD90^−^ population; **B**, FACS work flow for cell characterization; **C-E**, adipogenic, fibrogenic and myogenic differentiation of CD34^+^CD90^−^CD56^−^ (**C**), CD34^+^CD90^+^CD56^−^ (**D**) and CD56^+^CD82^+^CD34^−^ (**E**) populations verified by immunocytochemistry for Perilipin-1 (Red) and Collagen-1 (green) or Desmin (Red) and Myosin Heavy Chain (MyHC, green); **F**, Content of mononuclear cells in human skeletal muscle (n=17) in percent of the total number of single cells; **G**, Expression of PDGFRa in freshly sorted FAPs; **H**, Distribution of COL1A1 mRNA expressing CD90^+^CD56^−^ (Fibro-adipogenic progenitors; FAPs), CD56^+^ (Muscle stem cells; MuSCs) and CD90^−^CD56^−^ (Un-identified) cells in human skeletal muscle (n=6) after activation for 3-9 days in vitro; **I**, Work-flow for single-cell clonal experiment; **J**, Colony formation (percentage wells containing colonies) of single sorted FAPs and MuSCs (n=12); **K**, Limiting dilution assay of sorted FAPs and MuSCs (n=3) with fraction of negative wells as a function of deposited cells. Solid line represents non-linear fit; **L**, MyHC and Desmin positive myotubes from clonal cultures from single cell sorted MuSCs (stitched images from entire well); **M**, Adipogenic (Perilipin-1^+^) and fibrogenic (Collagen-1^+^) cells in clonal cultures from single cell sorted FAPs (stitched images from entire well); **N**, GO: Biological processes enriched in freshly sorted FAPs; **O**, Percentage of MuSCs and FAPs incorporating EdU (S-phase entry) post isolation (n=10-16); **P**, Images of EdU incorporation 96h post sort of MuSCs (**i**) and FAPs (**ii**). Significant difference denoted by *p<0.05, **p<0.01 and ***p<0.001.

The above-mentioned FACS-isolated populations and were tested for their adipogenic, fibrogenic and myogenic capacity in culture. Only the two CD34^+^ cell populations (CD90^−^ and CD90^+^) gave rise to both adipocytes and collagen/α-smooth muscle actin expressing cells when cultured under appropriate conditions (Fig 2 C-E; Fig S3 A i-ii; B, C i-ii). In contrast, the MuSCs only rarely gave rise to adipocytes (Fig 2 E), whereas these robustly differentiated into myosin heavy chain (MyHC)/Desmin expressing myotubes (Fig 2 E). Moreover, we found no adipogenic differentiation and very little collagen expression in endothelial or hematopoietic cells (Fig S3 D i-ii). The two minor populations (Fig 2 A iv) grew poorly and we were therefore unable to induce adipogenesis in these.

For FAPs we also noted osteogenic capacity (Fig S3 E i-iii, indicated by calcium deposit formation) after 10 days in osteogenic induction medium, as previously shown in rodent FAPs ^14,34^. This confirms the mesenchymal nature of FAPs, although heterotopic ossification is only rarely observed in humans^35,36^. We therefore chose to adhere to the FAP nomenclature. We confirmed the myogenic nature and purity of the MuSCs by Pax7 staining 24h post isolation followed by differentiation into multinuclear myotubes (Fig S3 F i-iv). Finally, we found that the CD90^+^CD56^−^CD34^−^CD45^−^CD31^−^ cells expressed NG2^+^and CD90 (Fig S3 G i-ii) and thus we speculate if these are smooth muscle cells or pericytes, as described by Giordani, et al. ^37^.

Using flow-data, we quantified the cell populations from which it appeared that FAPs (FAP^CD90−^ and FAP^CD90+^ combined) constitute a major proportion of the mononuclear (~40%) cells in human skeletal muscle (Fig 2 F). Staining of FACS isolated FAPs three days post isolation confirmed the expression of PDGFRα (Fig 2 G) as well as TCF7L2, TE7, and CD90 (Fig S3 H i-ii, I, J). To understand if the FAPs are the major collagen producing cells in human skeletal muscle, we digested non-diabetic human muscle biopsies and plated the entire content, allowing all adherent cells to attach and activate. After a week we detached the cells and probed for expression of *COL1A1* mRNA in FAPs, MuSCs and unidentified cells^38^. When selecting cells expressing *COL1A1* (Fig S4 A i-ii, B, C), it was apparent that FAPs was the major collagen producing population with only a minority of MuSCs and unidentified cells expressing *COL1A1* (Fig 2 H). Similar results were obtained when identifying FAPs with a PDGFRα antibody (Fig S4 B, C) ^24^. Furthermore, when selecting the entire population of FAPs, MuSCs or unidentified cells, it was clear that FAPs were the primary *COL1A1* producing cell type (Fig S4 E). Collectively, our results suggest that human skeletal muscle FAPs are likely the most represented cell population in human skeletal muscle and a putative source of fibro-fatty degeneration.

### Human FAPs are clonally and transcriptionally a distinct cell population

To understand if human FAPs have clonal (progenitor) potential and if these give rise to both adipocytes and fibroblasts, we performed a clonal assay, as previously described in mice^15^. FAPs and MuSCs were FACS isolated and immediately re-sorted in 96-wells with one cell per well (Fig 2 I). Notably, the MuSCs had approximately four times greater clonal potential (~17% in MuSCs versus ~4% in FAPs) than the FAP population (CD90^−^ and CD90^+^ combined, Fig 2 J). To substantiate these findings, we performed a limiting dilution assay, which predicted a markedly greater stem cell frequency (~10 times) of MuSCs compared to FAPs (Fig 2 K). In contrast, Joe, et al. ^15^ reported similar levels of clonality between FAPs and MuSCs, which, besides species-related differences, may be explained by the age of our donors compared to the age of the mice utilized by Joe, et al. ^15^. Aging has recently been shown to alter the phenotype of FAPs in mice, including the ability to enter the cell cycle ^39^. When differentiating the single sorted MuSC >92% of these clones gave rise to Desmin^+^/MyHC^+^ myoblasts/myotubes (Fig 2 L) similar to mouse MuSCs^15^. For the FAP colonies more than 72% contained Perilipin-1^+^ adipocytes and Collagen-1^+^ cells (Fig 2 M), while the remaining were non-adipogenic Collagen-1^+^ fibroblasts.

Next, in order to understand the nature of FAPs *in vivo*, we flow-sorted FAPs, MuSCs and the negative population and immediately processed these for transcriptomic analysis (RNA-seq). Principal component analysis, based on the expression of all detected genes, of the cell populations revealed that FAPs cluster separately from the other two sorted populations (Fig S4 F, G, H, I, J, K) and are enriched in FAP markers (Fig S4 G, H, I, J, K). Gene enrichment analysis of FAPs (CD90^−^ and CD90^+^ combined) revealed that FAPs *in vivo* display a gene expression pattern associated to ECM production, turnover and signaling, and also suggesting potential contribution to angiogenic events (Fig 2 N, S5A). In contrast, the MuSCs were enriched for straited muscle development and cell cycle control (Fig S5 A, B).

The negative population was devoid of MuSC markers, however, there was a specific expression of other myogenic genes (e.g. MyHC, Titin). Enrichment analysis confirmed this and revealed that these were myonuclei (Fig S5 C, D, E, F). Comparing FAPs to myonuclei confirmed their unique phenotype in relation to ECM regulation and angiogenesis (Fig S5 C, F). Collectively, these data support the existence of human FAPs and their potential contribution to fibro-fatty degeneration of human skeletal muscles.

MuSCs are known to be quiescent *in vivo* under homeostatic conditions, however, it has not been described if this is also the case for FAPs. Given their continuous role in ECM maintenance one might speculate that FAPs possess higher mitotic turnover, as compared to MuSCs. We also reasoned that FAPs implicated in fibro-fatty degeneration of T2DM muscles could have higher proliferative activity. To address this issue, we flow-sorted MuSCs and FAPs from human skeletal muscle biopsies and cultured them in wash-buffer plus EdU as described previously^40^. At 24h very limited EdU incorporation was detected in MuSCs as well as FAPs (0.0-0.8%), suggesting that the cells were quiescent at the time of extraction (Fig 2 O, P i-ii).

Moreover, while MuSC started to enter the S-phase between 24h and 48h, FAPs remained inactive until 48-72h and then slowly progressed through G1-S phase (Fig 2 O). This would suggest that human FAPs are mainly mitotically inactive *in vivo* (as MuSCs) and upon activation they display a slow kinetic of progression into S phase, as compared to MuSCs.

### Single-cell RNA-sequencing confirms the specificity and nature of the identified FAPs

We further investigated the transcriptional identity of human FAPs by performing single-cell transcriptome analysis (scRNA-seq) on crude human muscle tissue (mm. Rectus Abdominis and Gastrocnemius), using the 10x Genomics Chromium platform as previously utilized on mouse skeletal muscle ^37,41^ (Fig 3 A). After quality control (Fig S6 A-F) we performed an unsupervised clustering and initially identified 17 cell populations. Utilizing markers from mouse ^37,41^ as well as human skeletal muscle ^24,26,42^ we annotated nine major cell populations, with some containing one or several subpopulations (Fig 3 B). As expected, we found a population highly enriched in *PDGFRA* and *CD34* as well as partly expressing *THY1*/CD90 (Fig 3 C, D, E) and robust expression of *COL1A1* (Fig 3 F). This population had high expression of collagens, laminins and other matrix related proteins consistent with our FACS identified FAP population (Fig S7 A, B, C, S8). By using the FAP population specific genes as input for gene set enrichment analysis (GO: Biological processes, Fig S9 A) this was in agreement with the pathways observed in population-based RNA-seq data (Fig 2 N), including PDGF signaling. Interestingly, the PDGF pathway has been implicated in the regulation of the FAP quiescence ^18^. Under homeostatic conditions, mouse FAPs produce an intronic variant of PDGFRα, resulting in a decoy isoform of PDGFRα, which maintains FAPs in a non-cycling state ^18^. It is likely that the PDGF pathway also plays a key role in human FAPs.

**Fig 3.**
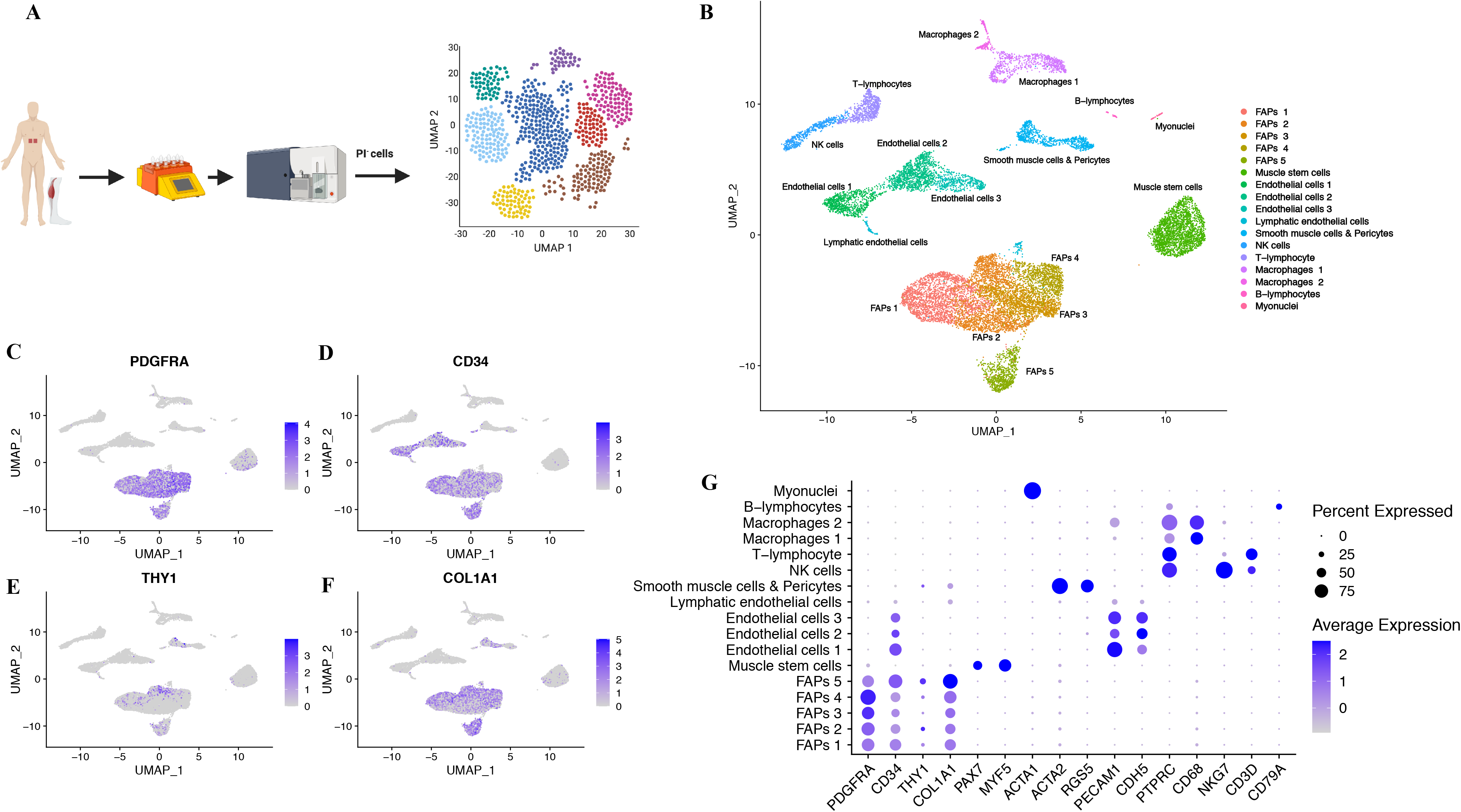
Single cell RNA sequencing confirms unique FAP population. **A**, Schematic presentation of work-flow single-cell RNA seq (scRNA-seq) experiment; **B**, UMAP cluster plot of all captured cells obtained from skeletal muscle single cell suspension (n=4) revealed 17 populations with some constituting subpopulations of a lager population; Feature plots of *PDGFRA* (**C**), *CD34* (**D**), *THY1* (CD90, **E**), *COL1A1* (**F**) expression; **G**, Dot-plot displaying differentially expressed genes in the identified 17 population (as in **B**). Coloring indicates mean expression and dot-size indicates percentage of cells in which the gene is expressed.

Our initial clustering resulted in five FAP subpopulations, which could indicate that the FAPs contain subsets with the potential to carry out distinct functions or at least sub-fractions at different stages. We noted one other population high in collagens and ECM proteins, as well as alpha smooth muscle actin (*ACTA2*), but negative for *CD34* and *PDGFRA* (Fig 3 G, Fig S8, S9 D). A similar population was recently described by Giordani, et al. ^37^ in which they were designated as smooth muscle-mesenchymal cells. In agreement with Giordani, et al. ^37^, the population is distinct from FAPs and given the much larger presence of FAPs (Fig 2 F, 3 B), we find it likely that FAPs constitute a more potent contributor to skeletal muscle fibrosis and adipocyte formation.

### PDGF controls FAP fibro- and adipogenic fate

Prompted by the finding that *PDGFRA* mRNA is specifically enriched in freshly sorted FAPs and in our scRNA-seq data, whereas protein expression was only detectable after 3-6 days in culture, we next asked what the role of PDGF signaling was in our identified human FAPs.

From our in-situ RNA hybridization flow-data, we noticed that PDGFRα expression was strongly associated with *COL1A1* expression in FAPs (Fig 4 A, B), suggesting that the ability to signal through the PDGF pathway is key for collagen expression. To test if PDGF stimulation increased collagen expression in freshly isolated FAPs, we isolated FAPs, exposed them to PDGF-AA (Fig 4 C) and found that Collagen-1 expression per cell was increased in response to PDGF-AA (Fig 4 D, E).

**Fig 4.**
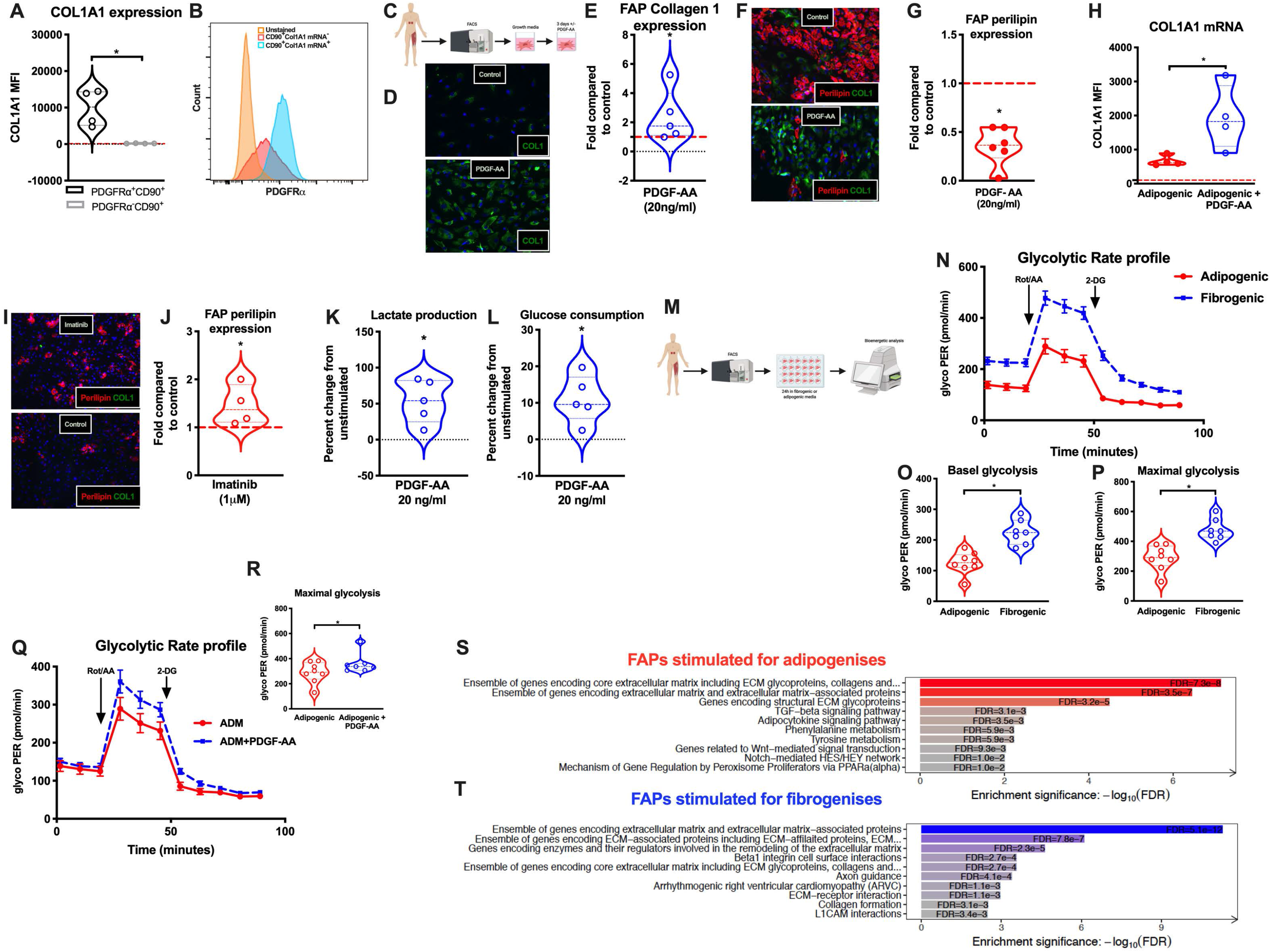
PDGF controls FAP differentiation and is associated metabolic alterations. **A**, COL1A1 mRNA expression (median fluorescence intensity, MFI) in PDGFRa^+^CD90^+^ and PDGFRa^−^CD90^+^ FAPs (n=4); **B**, PDGFRa expression in COL1A1 mRNA^+^ and COL1A1 mRNA^low/−^ FAPs; **C**, Schematic presentation of work-flow of PDGF-AA experiment; **D**, Collagen-1 protein (green) expression (positive area per cell) in FAPs following 24h of control (PBS) or PDGF-AA (20ng/ml) stimulation; **E**, quantification of Collagen-1 protein expression in FAPs following 24h of control (PBS) or PDGF-AA (20ng/ml) stimulation (n=5); **F**, Adipogenesis (Perilipin-1, red, positive area per cell) of FAPs with addition of control (PBS) or PDGF-AA (20ng/ml); **G**, Quantification of FAP adipogenesis (Perilipin-1 expression) with addition of control (PBS) or PDGF-AA (20ng/ml) (n=6); **H**, *COL1A1* mRNA expression during FAP adipogenesis (3 days) with addition of control (PBS) or PDGF-AA (20ng/ml) (n=4); **I**, FAP adipogenesis (Perilipin-1, red, positive area per cell) with inhibition of PDGF through Imatinib (1uM); **J**, Quantification of FAP adipogenesis with addition of control (PBS) or Imatinib (1uM) (n=4); **K**, FAP lactate production (change from unstimulated) with addition of PDGF-AA (20ng/ml) for 3 days (n=5); **L**, FAP glucose consumption (change from unstimulated) with addition of PDGF-AA (20ng/ml) for 3 days (n=5); **M**, Schematic presentation of work-flow of FAP bioenergetic profile experiments; **N**, Glycolytic profile (Glycolytic proton efflux rate; GlycoPER, pmol/min) of FAPs stimulated for 24h with adipogenic or fibrogenic media with sequential injections of Rotenone/Antimycin and 2-deoxy-glucose (n=7, technical replicates); **O**, Quantification of basal FAP glycolysis (glycolytic proton efflux rate; glycoPER, pmol/min per 6×10^4^ cells) following 24h with adipogenic or fibrogenic (containing 20ng/ml PDGF-AA) media; **P**, Quantification of maximal FAP glycolysis (glycolytic proton efflux rate; glycoPER, pmol/min per 6×10^4^ cells) following 24h with adipogenic or fibrogenic (containing 20ng/ml PDGF-AA) media; **Q**, Glycolytic profile (glycolytic proton efflux rate; glycoPER, pmol/min per 6×10^4^ cells) of FAPs stimulated for 24h with adipogenic media containing either control (PBS + Adipogenic) or 20ng/ml PDGF-AA (Adipogenic+PDGF-AA); **R**, Quantification of maximal FAP glycolysis (glycolytic proton efflux rate; glycoPER, pmol/min per 6×10^4^ cells) following 24h with adipogenic media containing either control (PBS+Adipogenic) or 20ng/ml PDGF-AA (Adipogenic+PDGF-AA); **S**, Pathways enriched in FAP stimulated for 6 days towards adipogenesei; **T**, Pathways enriched in FAP stimulated for 6 days towards fibrogenesis. Significant difference denoted by *p<0.05.

In addition to collagen production, earlier studies have reported that PDGF mediated signaling increases FAP proliferation ^18,24^. We confirmed the mitogenic effect of PDGF-AA in our human FAPs suggesting that this growth factor is also an important regulator of FAP activation and proliferation in human FAPs (Fig S10 A). In mice, constitutively active PDGF signaling is associated with widespread fibroses of organs and tissues, including adipose tissue and skeletal muscles ^22,23,43,44^. Thus, we asked whether PDGF-AA could affect the adipogenic differentiation of human FAPs. To address this, we cultured FAPs in adipogenic medium, while exposing them to PDGF-AA. Interestingly, PDGF-AA markedly reduced the ability to form Perilipin-1^+^ adipocytes from human FAPs (Fig 4 F, G) while maintaining/inducing a more fibrogenic nature as evidenced by increased collagen expression (Fig 4 F, H). Conversely, when inhibiting the PDGF pathway, using a tyrosine-kinase inhibitor (Imatinib), we could further increase FAP adipogenesis (Fig 4 I, J).

Collectively, our data reveal that PDGF signaling is a key determinant of human FAP cell fate, by promoting the expansion of FAP with fibrogenic potential at the expense of the adipogenic fate. A recent report on FAP regulation in aged mice found that FAP adipogenic potential is reduced with aging whereas fibrogenic conversion is increased ^39^, similar to what we report from PDGF-AA. Combined with other reports showing that inhibiting PDGF signaling can prevent skeletal muscle fibrosis ^16,18^, we propose that this pathway is key in FAP mediated skeletal muscle degeneration in humans. Considering these findings, it is interesting to speculate as to what the cellular source of PDGF-AA or PDGF-CC is *in vivo*. From our population-based RNA-seq data we found that the mature muscle fibers are particular high in *PDGFA* (Fig S10 B) and less in *PDGFC* (Fig S10 C). Moreover, examining our scRNA-seq data set indicate that also smooth muscle cells/ pericytes may be a source of this mitogen (Fig S10 D) in human skeletal muscle. Thus, our data suggest that PDGF-AA is key in controlling FAP behavior and paracrine signaling in the local niche can thereby influence FAP fate, although the needs to be confirmed directly.

### Increased reliance on glycolysis and lactate production is associated with FAP fibrogenesis

In addition to its role in fibrosis PDGF signaling has been associated with increased cellular reliance on glycolysis and lactate production for energy and substrate generation^45^. Thus, we investigated the potential relationship between metabolism and PDGF signaling in cell fate determination of human FAPs. Exposure to PDGF increased lactate production (Fig 4 K) and glucose consumption (Fig 4 L), in human FAPs, as compared to untreated cells. This prompted us to do a more detailed metabolic profiling using real-time bioenergetics analysis. Before bioenergetic analysis the FAPs were induced for 24h towards adipogenesis or fibrogenesis (PDGF-AA stimulation, Fig 4 M). We found no difference in basal or maximal oxygen consumption (Fig S10 E). In contrast, we observed a robust activation of both basal and maximal glycolysis (Fig 4 N, O, P), suggesting a functional link between activation of glycolysis (lactate fermentation) and the fibrogenic phenotype of human FAPs. When adding PDGF to FAPs induced for adipogenesis for 24h this also increased maximal glycolysis (Fig 4 Q, R) suggesting that this metabolic switch may be involved also in the concurrent reduction of adipogenesis in FAP stimulated with PDGF. This enhanced glycolysis is likely instrumental for the enhanced production of collagen since the glycolytic pathway is essential for amino acid synthesis of glycine as well as glycosylation, hydroxylation etc ^4^.

Since no previous studies have isolated human skeletal muscle FAPs without prior culture, limited knowledge exists on the transcriptome of human FAPs undergoing differentiation. To provide greater detail of this process we performed RNA-seq on FAPs cultured for six days in adipogenic or fibrogenic (PDGF stimulation) differentiation media. Indeed, altered transcripts related to the PPARα, a major lipid metabolism regulator ^46,47^, were enriched in the adipogenic FAPs (Fig 4 S; S10 F), along with common genes enriched during adipogenic differentiation (Fig S10 G, H, I, J). By contrast fibrogenic FAPs exhibited strong enrichment for gene implicated in ECM production, turnover and signaling (Fig 4 T, S10 F). Interestingly, in the fibrogenic FAPs, we noted a strong enrichment in Periostin (*POSTN*, Fig S10 F), an extracellular matrix protein acting as a ligand and previously associated with fibrosis and wound healing ^48^. More recently, Periostin has been shown to increase in activated FAPs during regeneration in mouse skeletal muscle along with lysyl oxidase enzymes (*LOX*) ^49^. Notably, the lysyl oxidase homolog 4 (*LOXL4*) was the most differentially expressed gene in our fibrogenic FAPs (Fig S10 F). This finding strongly supports the ability of PDGF signaling to stimulate a fibroblast or myofibroblast conversion of FAPs. We therefore propose that PDGF stimulates the fibrogenic nature of human FAPs and conversely reduces the adipogenic differentiation capacity by inducing a metabolic switch to increased reliance on glycolysis.

We next speculated if different subsets of FAPs exist *in vivo* with varying ability to transform into adipocyte or fibroblasts based on PDGF signaling. Recent studies in mouse and human adipose tissue have revealed stromal cell subpopulations *in vivo*, with some being more pro-adipogenic and others displaying a more regulatory or progenitor-like phenotype ^50,51^. Moreover, Malecova, et al. ^52^ recently reported a fibrogenic FAP subpopulation in mouse skeletal muscle (marked by VCAM1 expression), which increased during muscle degeneration in mdx mice. To address this question, we took advantage of our scRNA-seq data. We re-clustered the FAP population (Fig 3 B, FAPs 1-5), which resulted in seven subpopulations (Fig S11 A), from which we generated a heatmap of top-twenty differentially expressed genes (Fig S12). We immediately noted the high expression of *THY1* (CD90) mainly in the cluster 4 (Fig S11 B, C, S12). This was of interest, since we had already noted that CD90 marks a subset of FAPs (Fig 2 A ii). We therefore decided to investigate if the expression of CD90 defined a specific human FAP population with a distinct phenotype.

### THY1 marks a specific subpopulation of skeletal muscle FAPs *in vivo*

As a first step we examined if there was a correlation between the transcriptome of FAP^CD90−^ and FAP^CD90+^ and the DE genes in the muscle biopsies from T2DM patients, to indicate clinical relevancy. To do this, we compared DE genes in itT2D with DE genes upregulated in FAP^CD90+^ and FAP^CD90−^ (Fig S13 A). Strikingly, this revealed that genes upregulated in FAP^CD90−^ tended to be downregulated in T2DM, whereas genes upregulated in T2DM tended to be upregulated in FAP^CD90+^ (Fig S13 A). Based on this, we propose that in particular the FAP^CD90+^ have a central role in the development of the degenerative phenotype observed in T2DM muscle.

To investigate subpopulation specific traits, we initially inspected our flow data for potential differences in morphology and noted that the FAP^CD90+^ were larger in cell size than the FAP^CD90−^ (Fig 5 A). We then asked if the larger cell size could be related to the higher mitotic activity of FAP^CD90+^. Following isolation both FAP^CD90+^ and FAP^CD90−^ were completely EdU negative for the first 24h (Fig 5 B), suggesting that both were quiescent *in vivo*. However, at 48h, 72h and in particular 96h, the FAP^CD90+^ displayed an increased number of EdU^+^ cells compared to FAP^CD90−^, suggesting that the FAP^CD90+^enter the cell cycle more rapidly than FAP^CD90−^ (Fig 5 B). As such, although inactive *in vivo*, the FAP^CD90+^ have an increased propensity to enter the cell cycle, which is associated with a larger cell size. This finding suggests that FAP^CD90+^ could be a subset of progenitor cells within FAPs that is poised for entry into cell cycle and further fibrogenic activation, as proposed for cardiac FAPs^53^. To examine this further, we decided to perform the limiting dilution and single-cell clonal assay. In agreement with our EdU data, the FAP^CD90+^ had a greater level of clonality/progenitor formation than the FAP^CD90−^ (Fig 5 C, S13 B) and the single-hit Poisson model predicted a robust difference in stem cell frequency between the two FAP populations. In contrast, the clones formed by the FAP^CD90−^ had a greater proportion of clones becoming adipogenic, while in the FAP^CD90+^ the content of adipogenic and non-adipogenic clones was similar (Fig 5 D, E i-ii, S13 C). Collectively, our data support a model in which the FAP^CD90+^ have a progenitor phenotype, whereas the FAP^CD90−^ are prone towards adipocyte differentiation or have other stromal support functions.

**Fig 5.**
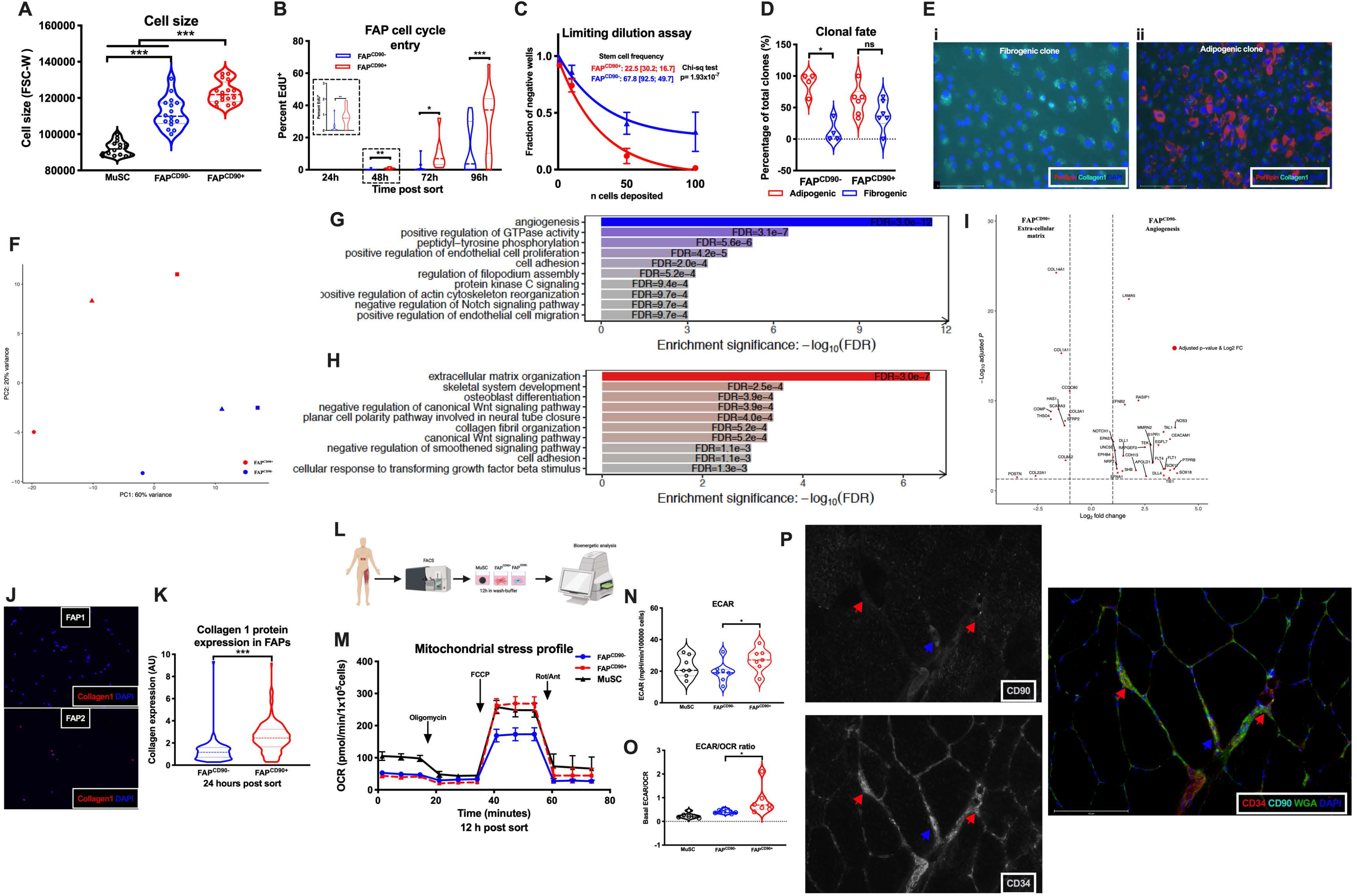
Two FAP populations are present in vivo defined by the expression of *THY1* (CD90) **A**, Cell size (FSC-W) from FACS on MuSCs, FAP^CD90−^ and FAP^CD90+^ (n=16); **B**, EdU incorporation of FAP^CD90−^ and FAP^CD90+^ at 24, 48, 72 and 96 hours (h) post FACS (n=9-13); **C**, Limiting dilution assay of freshly sorted FAP^CD90−^ and FAP^CD90+^ (n=3) with fraction of negative wells as a function of deposited cells. Solid line represents non-linear fit; **D**, Percentage of adipogenic (red) or fibrogenic (blue) clones from single sorted FAP^CD90−^ and FAP^CD90+^; **E**, Example images stained with Collagen-1 (green) and Perilipin-1 (red) containing either mainly fibrogenic (i) or adipogenic (ii) clones; **F**, Principal component analysis (PCA) plot based on differentially expressed genes in freshly sorted FAP^CD90−^ and FAP^CD90+^ (n=3); **G**, GO: Biological processes enriched in FAP^CD90−^ (n=3); **H**, GO: Biological processes enriched in FAP^CD90+^(n=3); **I**, Volcano plot (Log2 change versus −Log10 p-value) of genes underlying angiogenic (FAP^CD90−^) or extra-cellular matrix (ECM, FAP^CD90+^) processes enriched in FAP^CD90−^ and FAP^CD90+^(n=3); **J**, Images displaying Collagen-1 (Red) protein expression in FAP^CD90−^ and FAP^CD90+^ cells 24 hours post FACS; **K**, Quantification of Collagen-1 single cell protein expression 24h post sort (n=5); **L**, Schematic work-flow of bioenergetic profiling experiment; **M**, Mitochondrial profile (Oxygen consumption rate, pmol/min) of freshly sorted Muscle stem cells (MuSCs), FAP^CD90−^ and FAP^CD90+^ with sequential injections of Oligomycin, Carbonyl cyanide-*4*-(trifluoromethoxy) phenylhydrazone (FCCP) and Rotenone/Antimycin (n=7); **N**, Basal normalized extracellular acidification rate/glycolysis (ECAR, mpH/min) of MuSCs, FAP^CD90−^ and FAP^CD90+^ (n=7); **O**, Ratio between basal EACR (extra-cellular acidification rate) and OCR of freshly sorted MuSCs, FAP^CD90−^ and FAP^CD90+^; **P**, Immunohistochemical staining of skeletal muscle cross-section of CD34 (Red), CD90 (Cyan), wheat germ agglutinin (WGA, green) and DAPI (blue). Blue arrow indicates a CD34^+^ CD90^+^ cells, while red arrows indicates CD34^+^CD90^−^ cells. Significant difference denoted by *p<0.05.

### FAP^CD90+^ are poised for extracellular matrix production associated with an increased glycolytic flux

Given the phenotypic difference between the two FAP populations, we decided to directly compare the population-based transcriptome of freshly sorted FAP^CD90−^ and FAP^CD90+^ cells. Principal component analysis confirmed that the two groups cluster separately (Fig 5 F), supporting that CD90 expression defines a distinct population in accordance with the scRNA-seq data. Gene set enrichment analysis revealed a marked enrichment in processes related to angiogenesis and regulation of endothelial cell proliferation/migration in the FAP^CD90−^ (Fig 5 G, I). In contrast the FAP^CD90+^ genes were enriched in processes related to ECM development, collagen formation and response to TGFβ (Fig 5 H, I). We confirmed this using our generated scRNA-seq data by dividing the FAPs into CD90^+^ and CD90^−^ (Fig S14 A-B). To validate our transcriptomic findings, we flow-sorted FAP^CD90−^ and FAP^CD90+^ and stained for Collagen-1 24h post sort (Fig 5 J, K). As expected, the FAP^CD90+^ showed a higher Collagen-1 protein expression compared to FAP^CD90−^ (Fig 5 J, K).

Given the PDGF-phenotype of FAP^CD90+^ and the metabolic changes associated with PDGF-AA treatment, we sought to further investigate the FAP^CD90+^ metabolism *ex vivo*. To ensure that the FAPs were as close as possible to *in vivo*, we performed the bioenergetic analysis 10-12h post isolation, a time at which none of the FAP populations would be engaged in the cell cycle (Fig 5 L). At baseline the FAP populations were similar in oxygen consumption (Fig 5 M; Fig S14 C). In contrast, glycolysis was greater in FAP^CD90+^ compared to FAP^CD90−^ (Fig 5 N) supporting the hypothesis that the ECM producing FAP^CD90+^ phenotype is associated with an increased reliance on glycolysis. This difference was further exacerbated when calculating a glycolysis/oxygen consumption ratio (Fig 5 O). In addition, the maximal oxygen consumption was also higher in FAP^CD90+^ compared to FAP^CD90−^ (Fig 5 M; S14 D), suggesting a higher capacity for energy generation in FAP^CD90+^.

Staining for CD90 and CD34 in skeletal muscle tissue sections confirmed the presence of both CD90^+^CD34^+^ and CD90^−^CD34^+^ cells (Fig 5 P) *in vivo* in the muscle tissue, excluding that these are circulating cells, although we could unfortunately not combine CD90 and CD34 with CD31 to exclude endothelial cells from the stain. In summary, at least two distinct population of FAPs exist in human skeletal muscle with the FAP^CD90+^ being more primed for producing progeny and delivering ECM than FAP^CD90−^ (Fig S14 E).

### FAP^CD90+^ accumulate in skeletal muscle of T2DM patients

The data presented so far suggest an association between pathogenic fibrotic degeneration of skeletal muscle from T2DM patients and enrichment in DE genes typically driven by PDGF in fibrogenic FAP^CD90+^. Thus, we postulated that FAP^CD90+^ exhibiting a PDGF-mimetic phenotype could accumulate in muscles of T2DM patients and contribute to their pathological alterations.

We collected biopsies from patients (with or without T2DM) undergoing either abdominal aneurysm or coronary bypass surgery. When comparing the total FAP content, we noted that patients with T2DM tended to have a higher content of total FAPs (Fig 6 A, B). However, the difference between non-diabetics and T2DMs became even more pronounced, when we divided the FAPs into FAP^CD90+^ and FAP^CD90−^ and noted a selective increase in FAP^CD90+^ (Fig 6 C, D, E). In fact, the only population in which we identified significant changes was the FAPs and in particular the FAP^CD90+^ (Fig S15 A, B, C, D). These findings support a specific upregulation of FAP^CD90+^ in T2DM. PDGF has previously been shown to be increased in T2DM ^22,54^ and our data suggest that this is a strong modulator of human FAPs, and specifically the FAP^CD90+^.

**Fig 6.**
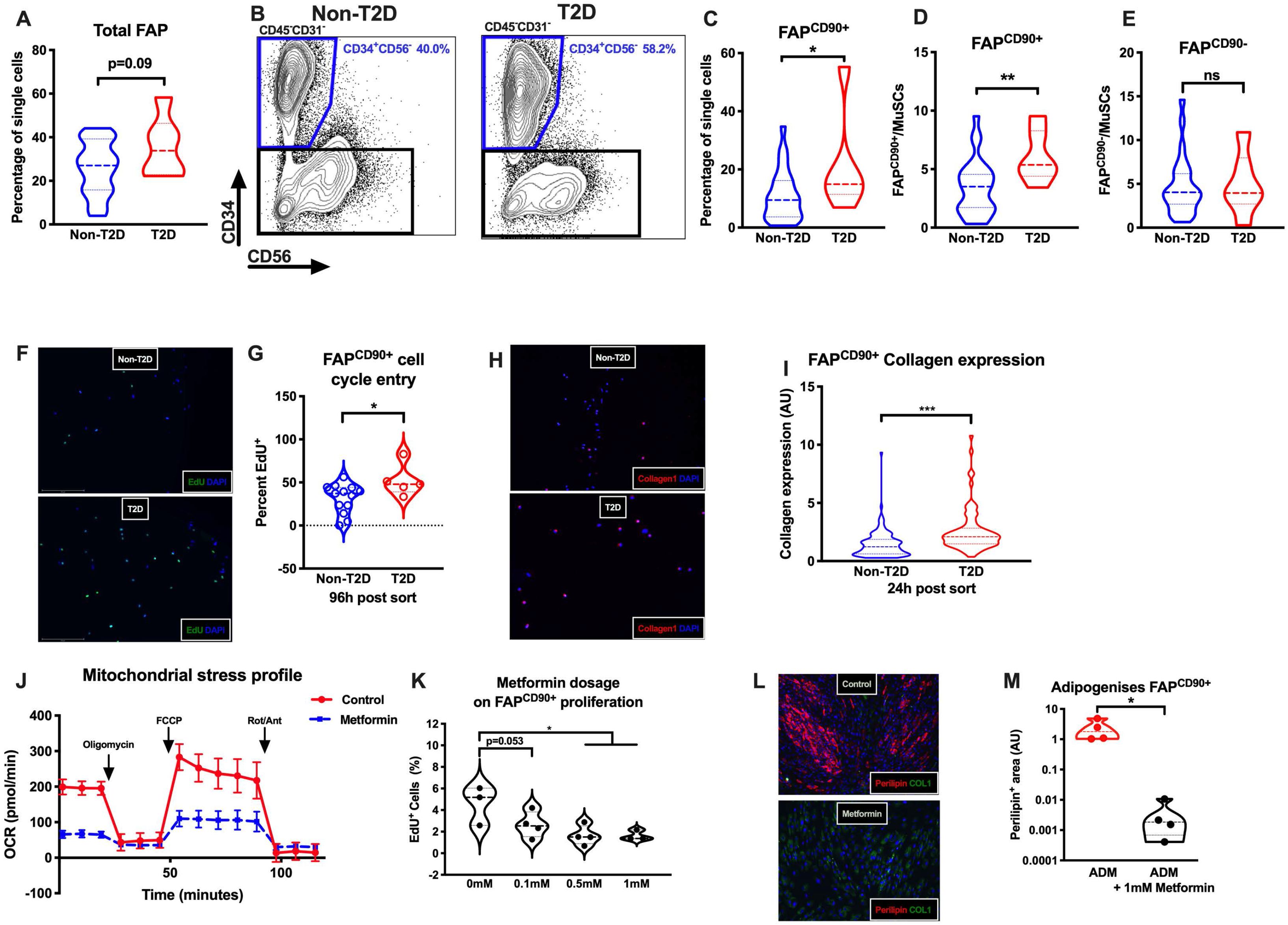
FAP^CD90+^ accumulate in type 2 diabetic skeletal muscle and are responsive to metformin treatment. **A**, Total FAP content (percentage of single cells, %) in patients without type 2 diabetes (Non-T2D, n=27) or with type 2 diabetes (T2D, n=6); **B**, Contour flow-plots showing the total FAP population (CD34^+^CD56^−^CD45^−^CD31^−^) in a Non-T2D patient and a T2D patient; **C**, FAP^CD90+^ content (percentage of single cells, %) in Non-T2D and T2D patients; **D**, FAP^CD90+^ content normalized to muscle stem cell (MuSC) content (FAP^CD90+^/MuSC, %) in Non-T2D and T2D patients; **E**, FAP^CD90−^ content normalized to muscle stem cell content (FAP^CD90−^/MuSC, %) in Non-T2D and T2D patients; **F**, Images displaying EdU+ FAPs 96h post sort from a Non-T2D or T2D patient; **G**, Cell cycle entry (EdU incorporation) in FAP^CD90+^ Non-T2D or T2D patients 96 hours post FACS (n=4-12); **H**, Images displaying Collagen-1 (red) protein expression in FAP^CD90+^ 24 hour post FACS in Non-T2D or T2D patients; **I**, Collagen1 protein expression in single cell FAP^CD90+^ from Non-T2D and T2D patients 24h post sort (n=2-3); **J**, Mitochondrial profile (Oxygen consumption rate, OCR, pmol/min) on FAP^CD90+^ exposed to control (PBS) or 1mM Metformin for 24h prior to bioenergetic analysis. Analysis was performed with a basal period followed by sequential injections of Oligomycin, Carbonyl cyanide-*4*-(trifluoromethoxy) phenylhydrazone (FCCP) and Rotenone/Antimycin (n=5-6, technical replicates); **L**, FAP^CD90+^ cell-cycle entry (EdU incorporation) following 24h of control (PBS), 0.1mM, 0.5mM or 1 mM Metformin; **M**, Images displaying FAP^CD90+^ adipogenesis (Perilipin-1, red, Collagen-1, green) in with addition of control (PBS) or Metformin (1mM) for 14 days; **N**, Quantification of FAP adipogenesis (Perilipin-1 positive area) with addition of control (PBS) or Metformin (1 mM). Significant difference denoted by *p<0.05, ** p<0.01, ***p<0.001.

Two other candidates for controlling FAP fate, given their central role in T2DM, is glucose and insulin. Interestingly, we found that insulin does increase FAP proliferation in general (Fig S15 E) as well as the first entry into the cell cycle post FACS isolation (Fig S15 F). In contrast, we found no effect of high glucose on FAP cell-cycle entry, suggesting that glucose per se does not trigger or accelerate FAP cell cycle entry (Fig S15 F). Importantly, we found that flow-sorted FAP^CD90+^ from patients with T2DM showed higher propensity to enter the cell cycle (Fig 6 F, G) and displayed an increased Collagen-1 expression 24h post sort (Fig 6 H, I), as compared to FAPs from non-diabetic patients. This result suggests that T2DM muscles provide a permissive environment for selective expansion of FAP^CD90+^, leading to their accumulation and subsequent tissue degeneration.

### FAP^CD90+^ can be targeted with metformin to decrease proliferation and inhibit adipogenesis

As a final question we asked if we could pharmacologically target FAP^CD90+^ to prevent excessive FAP accumulation in the muscles of T2DM patients. Several receptor tyrosine kinase inhibitors have been shown to effectively reduce FAP proliferation in mice including nilotinib and imatinib ^16,19,43^. The latter is interesting since imatinib is transported across the cell membrane using cation transporters (SLC22A1 and SLC22A3) that are also essential for transport of the first-in-line T2DM drug; metformin^55^. Metformin is, among other mechanisms, thought to increase cellular AMPK levels by inhibiting complex 1 in the respiratory chain and thereby limiting ATP production from oxidative phosphorylation ^56^. To examine if FAP^CD90+^ metabolism was responsive to metformin treatment we incubated FAP^CD90+^ with metformin (1mM) for 24 hours prior to measuring oxygen consumption and glycolytic rate (no metformin in the assay media). As predicted metformin markedly reduced oxygen consumption in FAP^CD90+^ (Fig 6 J). Conversely, metformin increased glycolysis in FAP^CD90+^ (Fig S15 G). Considering these findings, we predicted that metformin would also efficiently lower FAP^CD90+^ proliferation. Indeed, this was exactly what we observed when exposing FAP^CD90+^ to a titration of metformin dosages (Fig 6 K). Importantly, even at (pharmacologically relevant) dosages around 0.1 mM metformin we observed an effect on proliferation. We therefore speculate that metformin exerts this effect *in vivo* and reduce the expansion of FAP^CD90+^ population.

Moreover, metformin completely blocked the ability of FAP^CD90+^ to form adipocytes (Fig 6 L, M), while increasing Collagen-1 protein expression (Fig S15 H), in agreement with the marked shift towards glycolysis. Although metformin may enhance reliance on glycolysis and Collagen-1 production, the major effect on FAP proliferation is likely to overall reduce the development of fibrosis. However, even though metformin can potentially prevent excessive FAP proliferation, over time, FAP^CD90+^ still appears to accumulate. Thus, combinational treatments for targeting FAPs may be needed to overcome the degeneration of skeletal muscle. However, our data support that metformin may be a potential treatment in conditions associated with multi-organ wide fibrosis and degeneration (incl skeletal muscle), to target highly proliferative FAPs (FAP^CD90+^).

## DISCUSSION

In the present study, we show that human skeletal muscle contains several populations of transcriptionally and functionally distinct FAPs and that a specific subpopulation of the FAPs is a key mediator of the muscle degeneration observed among patients with T2DM. This work highlights that the T2DM condition is characterized by a substantial degeneration of the muscle niche. The identification of the cellular drivers of this degeneration provides the opportunity to target these cells to reduce tissue degeneration and ultimately preserve muscle function.

As a proof-of-concept we included two groups of T2DM patients, to display how the muscle remodeling is altered in the context of less (T2D) and more severe (itT2D) states of insulin resistance and T2DM. We initially found the ECM remodeling to be particular evident in patients treated with high doses of insulin and with a long disease duration (itT2D group). This could indicate that itT2D patients may represent a sub-group of patients with severe insulin resistance, which are more prone to diabetic complications. Severe insulin resistance is a defining feature of a sub-group of T2DM patients identified earlier in an non-biased approach^57^. This group of patients carry an increased risk for complications such as kidney fibrosis and ultimately kidney failure^57^. Interestingly, renal dysfunction and kidney failure in T2DM patients is associated with elevated whole-body collagen production (Collagen 6 formation^58^), which is also highly expressed in skeletal muscle and FAPs. The itT2D patients have many similarities to the insulin resistant patient cluster identified earlier^57^ and may carry a similar risk profile for diabetic complications such as degeneration of skeletal muscle and kidney failure.

Despite the greater remodeling in itT2D patients, the protein expression of PDGFRα in muscle homogenates, indicative of FAP content, was only increased in T2D patients with a shorter disease duration and receiving oral antidiabetic agents. The lower FAP content in the itT2D patients likely reflects differentiation of FAPs into myofibroblasts and adipocytes, a process in which the FAPs loose expression of PDGFRα^31^. This is supported by the extensive fibrogenic remodeling observed in the itT2D patients. As such, the muscle remodeling as a function of disease duration is consistent with the progression from initial increased FAP content in the T2D group, to FAP differentiation in the itT2D group. Earlier reports have shown muscle remodeling to be present already in obese subjects at whole tissue level^10,12,59^. Moreover, there is literature supporting a reduction in muscle quality and accumulation of non-contractile tissue in muscle from obese subjects as well as T2DM patients ^8,60–65^. Thus, it is possible that the muscle remodeling is initiated at very early stages of the disease and that the degenerative profile in the itT2D patients is related to the longer disease duration^28^. None-the-less, we view this remodeling as a complication to T2DM, rather than an underlying etiology to the initial development of insulin resistance and later T2DM that has been demonstrated in risk groups by Warram, et al. ^66^.

In addition to identifying CD34^+^ FAPs we show that that the mitogen; PDGF-AA increases FAP proliferation and collagen production, while also effectively reducing FAP adipogenesis. Importantly, the *in vivo* identified FAP^CD90+^ sub-population carried a phenotype with similar traits; i.e. enrichment for ECM production, high ability to proliferate and increased reliance on glycolysis. Moreover, the clones derived from FAP^CD90+^ were less adipogenic, consistent with the metabolic phenotype. ^45^. This effect is likely not restricted to PDGF-AA but true for other stimulators of ECM protein formation such as TGFβ1^4^ thereby linking ECM protein synthesis to enhanced reliance on glycolysis. Increasing glycolytic flux and lactate production not only delivers ATP, but is also key for essential aspects of collagen formation such as glycine synthesis and collagen hydroxylation ^4,67,68^. In contrast, cell proliferation was not dependent on glycolysis in agreement with previous studies^69^. Thus, we observed that human MuSCs are less glycolytic than the FAP^CD90+^, despite having a substantially faster cell-cycle entry post isolation. Instead, maximal mitochondrial respiration was increased in MuSCs and FAP^CD90+^ compared to FAP^CD90−^. We therefore speculate if the ability to increase ATP production through oxidative phosphorylation is key for cell-cycle entry and progression^70^, similar to stem cells transitioning from G^0^ into an alert stage^70,71^.

The novel finding that human FAPs consist of at least two transcriptionally, metabolically and functionally distinct subsets is particular intriguing since we found these subsets to be altered in the context of T2DM. To our knowledge, this is the first time that functionally distinct FAP subpopulations have been described in human skeletal muscle and furthermore shown to be altered in the context of disease. Subpopulations of muscle FAPs have been described in mice in which VCAM1^+^ FAPs increased in acute and chronic muscle injury ^52^. While the functional role of these subpopulation in mice are yet to be clearly defined, the VCAM1^+^ FAPs were associated with a pro-proliferative and pro-fibrotic phenotype ^52^. We did not find enrichment for VCAM1 protein expression in neither of the FAP populations described here (data not shown), suggesting that the phenotypes observed in our samples are different from those seen in muscle injury per se.

PDGFRα^+^CD90^+^ fibroblasts/FAPs from mouse ears were recently shown to accumulate with age and display a gene expression signature consistent with them being primed to enter the cell cycle^72^, similar to the FAP^CD90+^ described here. Thus, the PDGF-mimic phenotype of FAP^CD90+^ in skeletal muscle may be present in other FAP containing tissues.

In conclusion, skeletal muscle in T2DM patients with a long disease duration and severe insulin resistance is characterized by extensive remodeling of the extracellular niche. This degeneration is driven by a population of human FAPs that are extensively present in human muscle biopsies. *In vivo*, human skeletal muscle contains a subpopulation of FAPs which can be identified by expression of CD90. This population is highly primed for entering the cell cycle, producing clonal progeny, ECM production and increased utilization of glycolysis, likely driven by PDGF. The FAP^CD90+^ accumulate in muscles from T2DMs and constitute a likely driver of the fibrotic remodeling of the muscle niche.

## AVAILABILITY OF DATA AND MATERIAL

All data underlying the findings in this manuscript are provided as part of the article. Raw data on population-based and single-cell gene expression have been deposited in the GEOXXXX. The raw data that are not already presented in the figures are available from the corresponding author upon reasonable request.

## AUTHORS’ CONTRIBUTIONS

Conception and design of research: JF, FDP, PLP, NJ. Performed experiments: JF, LL, JBJ, TB, ISR, CC, ABM, LM, EG, RGF, UK, SP, NE. Analyzed data: JF, JJ, LL, ISR, SP. Interpreted results of experiments: JF, FDP, PB, TS, THP, FMR, PLP, NJ. Prepared figures: JF, JJ. Drafted manuscript: JF, NJ. Approved final version of manuscript: All authors.

## ACKNOWLEDGEMENTS

Flow cytometry / cell sorting was performed at the FACS Core Facility, Aarhus University, Denmark. The patients and surgeons at Department of Thoracic Surgery, Aarhus University Hospital, Denmark, are thanked for their willingness to donate and assist with obtaining skeletal muscle tissue, respectively. Helle Zibrantsen is thanked for assistance with western blotting and cell culture.

## METHODS & MATERIALS

### Ethical approval

All participants and patients were given oral and written information, and gave written consent to participate in accordance with the declaration of Helsinki. The studies were approved by the Local Ethical Committee of Central Denmark Region

### Patients and protocol

Study one included seven age-matched (six males and one female) healthy overweight subjects (OBS), seven (three male and four female) type 2 diabetic patients (T2D) and six (four males and two females) insulin treated type 2 diabetic patients (itT2D)). Patients had their oral antidiabetic treatments (metformin) withdrawn two day before the study and their usual insulin treatment were replaced with a continuous infusion of short acting insulin (Actrapid, Novo Nordisk, Denmark) and glucose one day before the study. The rates of insulin and glucose infusions were adjusted to reach a plasma glucose level of 8 mM. More detailed patient information is described previously ^28^.

Study two included patients admitted to aortic aneurism surgery (m. rectus abdominis) or conarary artery by-pass surgery (CABG, m. gastrocnemius. Inclusion criteria were age >50 years and <75 years and BMI >20. Patients with Type 1 diabetes, Polymyalgia, Lipodystrophy, active chemo or radiation therapy, Thiazalidinedione treatment, current haemodialysis treatment or glucocorticoid treatment with six months prior to inclusion were excluded from participation. Furthermore, patients were classified into non-diabetics and T2DMs.

### Muscle Biopsies

For study one the same skilled physician obtained all skeletal muscle biopsies from m. vastus lateralis under local anesthesia with a Bergström needle after an overnight fast. The biopsies were collected early in the morning in a resting and non-stimulated situation. Muscle biopsies were immediately frozen in liquid-nitrogen and thereafter stored at −80°C until further analyses.

For study two all skeletal muscle biopsies were obtained from m. rectus abdominis or m. gastrocnemius under full anesthesia by a skilled surgeon. The biopsies for tissue dissociation were immediately submerged into 4°C wash-buffer [Hams F10 incl. glutamine and bicarbonate (Cat nb N6908 Sigma, Sigma-Aldrich, Denmark) 10% Horse serum (cat nb 26050088, Gibco, ThermoFisher Scientific, MA, USA), 1% Penstrep (cat nb 15140122, Gibco)]. For histology the samples were dissected free of visible fat and connective tissue. A well-aligned portion of the biopsy was immediately mounted in Tissue-Tek (Qiagen, Valencia, CA, USA), frozen in isopentane pre-cooled with liquid nitrogen and stored at −80°C until further analysis.

### Tissue dissociation

Muscle was transported from the operation suite to the laboratory within 15 min in ice-cold wash-buffer. Upon arrival the muscle biopsy was initially dissected free of visible tendon/connective tissue and fat. The biopsy was then divided into pieces of up to 0.5-0.8 grams and briefly mechanically minced with sterile scissors. The muscle slurry was then transferred to C-tubes (cat nb. 130-093-237, Miltenyi Biotec, Lund, Sweden) containing 8ml wash buffer including 700 U/ml Collagenase II (lot 46D16552, Worthington, Lakewood, NJ, USA) and 3.27 U/ml Dispase II (cat nb. 04 942 078 001, Roche Diagnostics, Basel, Switzerland). Mechanical and enzymatic muscle digestion was then performed at 37°C on the gentleMACS with heaters (cat nb. 130-096-427, Miltenyi Biotec) for 60 min using a skeletal muscle digestion program (37C_mr_SMDK1).

When digestion was complete 8 ml wash buffer was added to the single cell solution and this was filtered through a 70μm cell strainer and washed twice to collect any remaining cells. The suspension was centrifuged at 500g for 5 min and the supernatant removed. The cell pellet was resuspended in freezing buffer (StemMACS, cat nb. 130-109-558, Miltenyi Biotec) and stored 1-3 weeks at −80.

### Flow cytometry and FACS

Approximately 1.5 hour before FACS the frozen cell suspension was thawed until a small amount of ice was left and resuspended in 10ml wash buffer. The solution was centrifuged at 500g for 5 min and the supernatant removed to clear the freezing buffer. The cells were then resuspended in wash buffer and incubated in MACS human FcR blocking solution (20μl/sample, cat nb. 130-059-901, Miltenyi Biotec) and primary antibodies against CD45-FITC (12μl/sample, Cat nb. 130-114-567, Clone 5B1, Miltenyi Biotec), CD31-FITC (4μl/sample, Cat nb. 130-110-668, Clone REA730, Miltenyi Biotec), CD31-PerCP-Vio700 (4μl/sample, Cat nb. 130-110-673, clone REA730, Miltenyi Biotec), CD90-PE (3.2μl/sample, Clone 5E10, eBioscience, Thermofisher), CD56-BV421 (1:100, Cat nb. 562752, BD Bioscience, San Jose, CA, USA), CD82-PE-vio770 (Cat nb. 130-101-302 10μl/sample, Miltenyi Biotec) and CD34-APC (20μl/sample, clone 581, BD Bioscience) in darkness at 4°C for 30 min. Propidium iodide (PI, 10μl/sample, cat 556463, BD Bioscience) immediately before sorting to exclude non-viable cells. The suspension was washed in 10 ml wash-buffer and centrifuged at 500g. Finally, the sample resuspended in wash-buffer and filtered through a 30μm filter to remove any remaining debris/aggregates. Non-stained cells and single-color controls were prepared in combination with the primary (full color) samples. To ensure bright single-color controls for compensation, compensation beads (cat nb. 01-2222-41, eBioscience, ThermoFisher) was utilized. Cell sorting was performed using a FACS-AriaIII cell sorter (BD Bioscience) with 405nm, 488nm, 561nm and 633nm lasers. A 100μm nozzle at 20 psi was utilized to lower pressure/stress on the cells as well as prevent clogging. Gating strategies were optimized through multiple earlier experiments, which included various full color samples (Fig S2 A i-vi), unstained sample, single color samples and fluorescence minus one (FMO) controls for CD34, CD90, CD82 and CD56 (Fig S2 C i-iv). Cells were sorted into 4 degC cooled collection tubes containing wash-buffer. Data was collected in FACSdiva software and later analyzed in FlowJo (FlowJo 10.6.1, BD)

The following populations were FACS isolated; CD56^+^CD82^+^CD34^−^ Lin^−^(CD45 and CD31) PI^−^ (MuSCs), CD34^+^CD90^−^CD56^−^Lin^−^PI^−^ (FAPs^CD90−^), CD34^+^CD90^+^CD56^−^Lin^−^PI^−^ (FAPs^CD90+^), Lin^+^CD34^+^PI^−^ (Endothelial cells), Lin^+^CD34^−^PI^−^ (Hematopoietic cells), CD90^+^CD34^−^CD56^−^Lin^−^PI^−^ (smooth muscle cells/pericytes) and CD90^−^CD34^−^CD56^−^Lin^−^PI^−^ (unknown). The purity FAP and MuSC populations were checked following the sort by re-running the samples which yielded >96% pure populations (Fig S2 D-F) and later by immunocytochemistry (ICC) when cells were plated (Fig 2 G, S4 H-J).

### Single cell sorting and limiting dilution assay

For both single cell and limiting dilution assay the cell populations (MuSCs and FAPs) were first sorted in bulk and immediately resorted depositing 1, 10, 50 or 100 cells in individual wells of a 96-well half-area plate. The number of wells was 96 for single cell sorting, 36 for 10 cells, 36 for 50 cells and 24 for 100 cells. Plates were coated 1:1 with collagen (Cat nb C8919, Sigma) and laminin (Cat nb 23017-015, Gibco, Thermofisher). Media was wash-buffer + 20% FBS (Cat. nb. 16000044, Thermofisher) + 5ng freshly added bFGF (human rbFGF, Cat nb. F0291, Sigma). After two to three weeks the wells were scored for colonies (> 8 cells). Wells with colonies were allowed to grown to confluency before initiating differentiation for myogenic (DMEM (cat nb Cat. no. 11965092, Thermofisher, 4.5 g/L glucose), 5% FBS, 1% Penstrep) or adipogenic (Adipogenic differentiation media; ADM, Cat. no. 130-091-677, Miltenyi Biotec) linage. Wells were scored for being myogenic (Myosin heavy chain/Desmin expressing cells and myotubes), Adipogenic (containing >3 Perilipin-1 positive adipocytes) or fibrogenic (containing Collagen-1 positive cells and < Perilipin-1 positive adipocytes). Limiting dilution analysis calculations were based on the single hit Poisson model (see statistics section).

### Cell culture

All tissue-culture plates or chamberslides were treated with extra-cellular matrix (ECM gel, Cat nb. E1270, Sigma) and chamberslides were in additions pre-treated with Poly-D-Lysine (Cat nb A-003-E, Millipore, Sigma) to increase adherence of the ECM. Cells from FACS were initially plated at a density of approximately 1×10^4^/cm^2^. All cells were plated in washbuffer and after 24 h this was switched to a growth media (GM, Bio-AMF 2, Cat. nb. 01-194-1A, Biological Industries, CT, USA). Media was changed every 2-3 days. If cells were passaged, they were maintained <60% confluency. When reaching >95% confluency, the media was switched to a differentiation media, depending on the cell type. For myogenic differentiation we used DMEM (4.5g/L glucose), 5% FBS and 1% penstrep. For adipogenic differentiation we used either a complete adipogenic differentiation media (ADM, Miltenyi Biotec) or DMEM (4.5g/L glucose) + 20% FBS + 1μg/ml Insulin (Cat nb. 91077C, Sigma) + 0.25 μM Dexamethasone (Cat nb. D4902, Sigma) + 0.5 mM 3-isobutyl-1-methylxanthine (IBMX, Cat nb I7018, Sigma) + 5 μM Rosaglitazone (Cat nb. R2408, Sigma) and 1% Penstrep. For fibrogenic stimulation we utilized DMEM (4.5g/L glucose), 10% FBS, 1% Penstrep and 1 ng/ml TGFβ (Cat nb. T7039, Merck, Sigma) or 20 ng/ml PDGF-AA (human PDGF-AA, Cat nb. 130-108-983, Miltenyi Biotec). Inhibition of PDGF-signaling was performed by addition of 1μM imatinib (Cat nb. SML1027, Sigma) during adipogenic differentiation. To examine the effect of metformin on FAP proliferation and differentiation, FAP were exposed to metformin (Cat nb. D150959, Sigma) in either growth conditions or during adipogenic differentiation. Osteogenic differentiation was induced utilizing a mesenchymal osteogenic differentiation media (Cat nb. 130-091-678, Miltenyi Biotec).

### Immunohisto- and cytochemistry and imaging

For whole skeletal muscle tissue cryosections (8 μm) were cut at −18°C and placed on a slide and stored at −80°C. Before staining sections were allowed to reach room temperature. Sections were fixed in Histofix (Histolab, Gothenborg, Sweden) followed by 1 hour in blocking buffer (10% goat serum, 0.2% Triton X in PBS). Sections were stained for Collagen 1 (mouse, 1:1000; Cat. no. C2456, Sigma), Perilipin-1 (rabbit, 1:200, Cat. no. 9349, Cell Signaling Technologies), CD34 (rabbit, 1:100, Cat nb ab81289, Abcam, Cambridge, UK), CD90 (mouse, Cat nb 14-0909-80, eBioscience, Thermofisher) and incubated at 4°C overnight in blocking buffer. This was followed by incubation with the secondary antibody Alexa-fluor 647 goat-anti-mouse and Alexa-fluor 568 goat-anti-rabbit (1:500; Cat no A-21235 and Cat no A-11011, Invitrogen, Thermo Fisher) combined with Wheat-germ-agglutinin (WGA; Cat nb. W1126, Thermofisher) conjungated to Alexa-fluor 488 for 1.5 hours at room temparature. Finally, sections were washed 3×5 min, with one wash containing DAPI (1:50000, D3571, Invitrogen, Thermofisher), mounted with mouting media and stored in darkness at 4°C. For cells in plates or chamberslides, these were fixed for 10 min in 4% paraformaldehyde, washed twice in PBS and stored in PBS until staining. Cells were then incubated for 30 min in blocking buffer (10% goat serum, 0.2% Triton X in PBS). Afterwards cells were incubated with primary antibody against PDGFRa (goat, 1:200, AF-307-NA, R&D Systems, MN, USA), CD90 (mouse, 1:500, Cat nb 14-0909-80, eBioscience, Thermofisher), TE-7 (mouse, 1:200, Cat nb CBBL271, Merck, Sigma), Pax7 (mouse, 1:50, Pax7, Developmental Studies Hybridoma Bank, IA, USA), Desmin (rabbit, 1:200, Cat nb. D93F5, Cell Signalling Technologies), Myosin Heavy Chain (MyHC, mouse, 1:5, MF20, DHSB), Collagen 1 (1:500, Cat nb Cat. no. C2456, Sigma), alpha-smooth muscle actin (1:150, Cat nb. A5228, Sigma) or NG2 (rabbit, 1:200, AB5320, Millipore-Merck, Sigma) at 4°C overnight in blocking buffer. This was followed by incubation with the secondary antibody Alexa-fluor 647 goat-anti-mouse, Alexa-fluor 488 goat-anti-mouse and Alexa-fluor 568 goat-anti-rabbit (1:500; Cat nb A-21235, cat nb A11001 and cat nb A-11011, Invitrogen, Thermofisher) for 1.5 hours at room temperature. Finally, cells were washed 3×5 min, with one wash containing DAPI, and either maintained in PBS or mounted in mounted media. Minus primary controls were included for all stains during optimization to ensure specificity.

Alzirian red (Cat nb. A5533, Sigma) solution (2g/100ml dH2O, pH 4.2) staining was performed for two min and subsequently washed extensive with dH2O to stain for calcium depots (osteogenic differentiation) for 5 min and thoroughly was afterwards before imaging. Cells were stored in dH2O.

Images were acquired using a Leica DM2000 fluorescent microscope and a Leica Hi-resolution Color DFC camera (Leica, Stockholm, Sweden) or EVOS M7000 automated imaging system (ThermoFisher).

### Primeflow (RNA-flow cytometry)

For primeflow, muscle tissue was digested as previously described and all cells were plated in GM in a 25 cm^2^ tissue culture flask precoated with gelatine (0.2%). All cells were allowed to adhere for 3 days before the media was changed, to allow for more slowly or poorly adhering cells to attach. After 6-8 days the cells reached 80-90% confluency and were trypsinized and the primeflow protocol was initiated. Initially the cells were stained with antibodies against CD90-PE (3.2ul/sample, Clone 5E10, eBioscience), CD56-BV421 (1:100, Cat No.562752, BD Bioscience or PDGFRa-BV421 (1:100, Cat nb 562799, BD Bioscience) for 40 min in wash-buffer. The cell suspension was then washed, centrifuged at 500g for 5 min, the supernatant was removed and cells were resuspended in residual volume. Here after we followed the instructions provided by the manufacturer (PrimeFlow™ RNA assay kit, Cat no 88-18005-210, eBioscience/Thermo Fisher). In brief the samples were incubated with the first fixation buffer (PrimeFlow fixation buffer 1) for 30min in dark at 4°C. Samples were centrifuged for 5 min at 500g, supernatant removed and resuspended in residual volume. Primeflow RNA permabilization buffer including RNAase inhibitors were added, samples gently mixed and then centrifuged for 5 min at 500g, supernatant removed and samples resuspended in residual volume and then repeated once more. Samples were the incubated with a second fixation buffer (PrimeFlow fixation buffer 2) for 1h at room temp at in dark. The samples were then washed twice (PrimeFlow RNA wash buffer). Prewarmed target probe (1:20, COL1A1, NM_000088.3, Type 4 probe; RPL13A, NM_001270491.1, Type 1 probe) was then added to the samples and then placed in a hybridization oven at 40°C for 2h. Samples were again washed twice (PrimeFlow RNA wash buffer) with the final wash including RNAase inhibitors. Samples were left in final wash-buffer overnight in dark at 4°C.

On day 2 samples were initially incubated with a pre-amplification mix (1:1 PrimeFlow RNA Pre-AMP mix) for 1.5h in hybridization oven at 40°C. Samples were washed three times in washbuffer (PrimeFlow RNA wash buffer) and then incubated for 1.5h in the hybridization oven at 40°C (1:1, PrimeFlow RNA AMP mix). Samples were washed twice (PrimeFlow RNA wash buffer) and finally incubated with label probes (1:100, PrimeFlow RNA label probes) for 1h in hybridization oven at 40°C. Samples were the washed twice (PrimeFlow RNA wash buffer) and then analyzed using a BD LSR Fortessa (equipped with 405, 488, 561 and 640 nm lacers and 18 detectors). We only included one fluorophore per laser to minimize then need for compensation. Compensations were performed using compensation beads specific to the PrimeFlow kit. We performed negative controls by omitting the target probes but preforming the rest of the protocol as specified (Fig S3 J). Positive controls for “housekeeping” genes were included (Fig S3 J). Samples were initially gated on forward- and side scatter to exclude debris and dead cells, as well as doublets using forward scatter height versus area.

### 5-ethynyl-2’-deoxyuridine (EdU) incorporation

To detect proliferation/cell cycle entry we utilized the EdU/click-it detection platform to visualize cells that contained newly synthesized DNA (Cat nb. C10337 or C10340, Invitrogen, Thermofisher). For experiments performed immediately post FACS, cells were plated in 96-well half-area tissue culture mircroplates (Corning, Sigma) coated with ECM (Sigma). Cells were plated at approximately 1×10^4^/cm^2^ in wash-buffer + 10μM EdU and the fixed for 10 min in PFA at various timepoints post FACS. After two washes in PBS cells were stored in PBS in dark at 4°C.

For experiments in which we tested the effect of PDGF-AA (Miltenyi Biotec) or Insulin (Sigma) on FAP proliferation we serum starved (0.5% Horse serum) FAPs that were in the growth phase for 18-20h to synchronize the cells. After this we changed the media with media (F10 or DMEM and 0.5% horse serum) added control or PDGF-AA/Insulin to the cells + 10μM EdU. After 24h we fixed the cells for 10 min in PFA, washed them twice with PBS and stored them in PBS in dark at 4°C. To study the effect of glucose (1 g/L versus 4.5g/L) and Insulin (PBS versus 1ng/ml Insulin) on FAP activation following FACS we cultured the FAPs for 96h in DMEM + 0.5% Horse serum + 10uM EdU.

To detect EdU incorporation we followed the manufacturer instructions and counter-stained the cells with DAPI. Images were acquired using Images were acquired using an EVOS M7000 automated imaging system (ThermoFisher) and performed automatically to ensure similar treatment of all wells.

### Real time qPCR on whole skeletal muscle

Skeletal muscle (20 mg) was homogenized in TriZol reagent (Gibco BRL, Life Technologies, Roskilde, Denmark). RNA was quantified by measuring absorbance at 260 nm and 280 nm and the integrity of the RNA was checked by visual inspection of the two ribosomal RNAs on an ethidium bromide stained agarose gel. Reverse transcription was performed using random hexamer primers as described by the manufacturer (GeneAmp RNA PCR Kit from Perkin Elmer Cetus, Norwalk, CT). PCR-mastermix containing the specific primers and Taq DNA polymerase (HotStar Taq, Quiagen Inc. USA) was added. The following primers were designed using the primer analysis software Oligo version 6.64: B2M: Sense AATGTCGGATGGATGAAACC, Antisense TCTCTCTTTCTGGCCTGGAG; TGFB1: Sense GGACACCAACTATTGCTTCAGCTC, Antisense AAGTTGGCATGGTAGCCCTTGG; COL1A1: Sense TGCGATGACGTGATCTGTGACG, Antisense TTTCTTGGTCGGTGGGTGACTCTG; COL3A1: Sense ATTGCTGGGATCACTGGAGCAC, Antisense CCTGGTTTCCCACTTTCACCCTTG; COL6A1: Sense ATCAGCCAGACCATCGACACCATC, Antisense TTCGAAGGAGCAGCACACTTGC

Real time quantitation of target gene to B2M mRNA was performed with a SYBR-Green real-time PCR assay using an ICycler from BioRad. The threshold cycle (Ct) was calculated, and the relative gene-expression was calculated essentially as described in the User Bulletin #2, 1997 from Perkin Elmer (Perkin Elmer Cetus, Norwalk, CT).

### Western blotting

Frozen crude muscle tissue were homogenized in ice-cold lysis buffer (50 mM HEPES, 137 mM NaCl, 10 mM Na4P2O7, 10 mM NaF, 1 mM MgCl2, 2 mM EDTA, 1% NP-40, 10% glycerol (vol/vol), 1 mM CaCl2, 2 mM Na3VO4, 100 mM AEBSF [4-(2-aminoethyl) benzenesulfonyl fluoride], hydrochloride, pH 7.4) using a Precellys homogenizer (Bertin Technologies, France). Insoluble materials were removed by centrifugation at 14,000g for 20 minutes at 4 °C. Protein concentration of the supernatant was determined using a Bradford assay (BioRad, CA, USA). Samples were adjusted to equal concentrations with milli-Q water and denatured by mixing with 4x Laemmli’s buffer and heating at 95 °C for 5 minutes.

For cells, each well was washed twice in ice-cold PBS and the cells were scraped off in ice-cold PBS with added protease inhibitors (1:100 Halt, cat nb. 78429, Thermofisher; 5 mM NAM; 2 mM NaOV). The suspension was transferred to a tube and centrifuged for 10 min at 2000g at 4 °C. The supernatant was removed and the pellet placed in a −80°C freezer for 4 hours to lyse cells after which ice-cold lysis buffer was added.

Equal amounts of protein were separated by SDS-PAGE using the BioRad Criterion system, and proteins were electroblotted onto PVDF membranes (BioRad). Control for equal loading was performed using the Stain-Free technology that allows visualization of total protein amount loaded to each lane and has been shown to be superior to beta-actin and GAPDH in human skeletal muscle ^73,74^. Membranes were blocked for 2 hours in a 2% bovine serum albumin solution (Sigma-Aldrich, MO, USA) and incubated overnight with primary antibodies PDGFRa (1:1000, AF-307-NA, R&D Systems, USA), Collagen 1 (Cat nb. T40777R, Meridian Life Science Inc, TN, USA)) and Collagen 3 (Cat nb GWB-7D650E, Genway Biotec Inc, CA, USA). After incubation in primary antibodies the membranes were incubated 1 hour with HRP-conjugated secondary antibodies. Proteins were visualized by chemiluminescence (Pierce Supersignal West Dura, Thermo Scientific, IL, USA) and quantified with ChemiDocTM MP imaging system (BioRad). Protein Plus Precision All Blue standards were used as marker of molecular weight (BioRad).

### Glucose and lactate analysis

Pre-conditioned media from FAP cell culture stimulated 3 days with PDGF-AA (20 ng/ml, human PDGF-AA, Cat nb. 130-108-983, Miltenyi Biotec) or control was collected and stored at −20°C. For analysis samples were quickly defrosted and glucose and lactate concentrations were determined in triplicates on an YSI Select (YSI Life Sciences, Yellow Springs, OH, USA).

### Bioenergetic analysis

Realtime analysis of oxygen consumption rate (OCR) and extracellular acidification rate (ECAR), as a proxy of cellular glycolysis, was performed using the Seahorse technology.

For freshly FACS isolated cells, we plated 1×10^5^ cells in ECM (Sigma) coated Seahorse XF96 cell culture microplate (part nb 101085-004, Agilent) in wash-buffer (to minimize change in cell bioenergetics related to high concentration of growth-factors etc). The number of cells was optimized before running the experiment. MuSCs and FAPs were allowed to adhere for 12h before running the bioenergetic analysis. One hour before the analysis the cells were washed twice in phenol red and bicarbonate free DMEM Seahorse Media (SM) (part nb 103575-100, Agilent) containing 25 mM glucose (Cat nb. G8769, Sigma), 1 mM sodium-pyruvate (Cat nb. 11360-039, Gibco) and 2 mM L-glutamine (Cat nb. G7513, Sigma), (pH 7.4) and allowed to equilibrate in a non-Co2 incubator for an hour. During this time brightfield images of all wells were captured to ensure a sufficient monolayer of cells. For freshly sorted cells, we performed the mitochondrial stress test using the Seahorse Mito-stress test kit (part nb 103015-100, Agilent) with a final concentration of 4mM Oligomycin, 5mM FCCP and 2.5 mM Antimycin A/Rotenone diluted into the SM media. Bioenergetic analysis was performed on the Seahorse XFe96 Analyzer (Agilent). Following the bioenergetic analysis Hoechst 33342 (Cat nb. H3570, Invitrogen) was added to all wells and images obtained of all wells to count the number of cells in each well. The number of cells per well was directly incorporated into the Wave software (Seahorse Wave Controller 2.4 software, Agilent) to normalize OCR and ECAR values. When possible, we included up to three replicates for each cell type from each donor, however, for some donors/cell types it was only possible to include one or two replicates.

To assess bioenergetics during stimulation towards fibro- or adipogeneses in cultured FAPs we plated them in a XF24 cell culture plate at a density of 6×10^4^ per well resulting in a confluent monolayer in GM (several cell titrations were initially performed). Twenty-four hours before the bioenergetic profiling the media was switched to adipogenic (ADM, Miltenyi Biotec), ADM+20ng/ml PDGF-AA or fibrogenic (DMEM+5%FBS+20ng/ml PDGF-AA) differentiation media. One hour before the analysis the cells were washed twice in SM phenol red free and bicarbonate free DMEM (part nb 103335-100, Agilent), 25 mM glucose, 1 mM sodium-bicarbonate and 2 mM L-glutamine, (pH 7.4) and allowed to equilibrate in a non-Co2 incubator. For the glycolytic rate assay HEPES (part nb 103337-100, Agilent) was added to a final concentration of 5 mM. Before bioenergetic analysis the wells were examined to ensure a confluent monolayer of cells in all wells, otherwise these were excluded from the final analysis. To assess mitochondrial function we utilized the strategy described above, by sequentially injecting Oligomycin (4mM), FCCP (5mM) and Antimycin/Rotenone (2.5 mM) diluted into the SM media. To asses the glycolytic rate and profile this we sequentially injected Antimycin/Rotenone (2.5 mM) and 2-deoxy glucose (2-DG, 100mM). For this analysis all chemicals were purchased from sigma. Initial data analysis was performed in Wave software (Agilent).

### RNA sequencing on tissue and FACS isolated cell populations

For whole tissue in study one the protocol has been described previously ^27^. In brief, total RNA was purified from frozen biopsies using the QiaSymphony robot in combination with the QiaSymphony RNA Mini kit (Qiagen, CA, USA) according to the Manufacturers protocol including DNase treatment. We were not able to isolate muscle RNA from one of the itT2D patients and one of the OBS subjects, leaving six patients/ subjects in each of the groups for RNA-sequencing. RNA concentration was determined using a spectrophotometer with absorbance at 260 nM (NanoDrop ND-1000) and RNA integrity was assessed using a 2100 Bioanalyzer (Agilent Technologies, Santa Clara, CA, USA). Whole transcriptome, strand-specific RNA-Seq libraries facilitating multiplexed paired-end sequencing were prepared from 500 ng total-RNA using the Ribo-Zero Magnetic Gold technology (Epicentre, an Illumina company) for depletion of rRNA followed by library preparation using the ScriptSeq v2 technology (Epicentre). The RNA-Seq libraries were combined into 2 nM pooled stocks, denatured and diluted to 10 pM with pre-chilled hybridization buffer and loaded into TruSeq PE v3 flowcells on an Illumina cBot followed by indexed paired-end sequencing (101 + 7 + 101 bp) on an Illumina HiSeq 2000 using TruSeq SBS Kit v3 chemistry (Illumina). Paired de-multiplexed fastq files were generated using CASAVA software (Illumina) and processed using tools from CLC Bio (QIAGEN).

For sorted cells, these were FACS isolated into tubes containing wash-buffer as previously described. After sorting the cells were transferred to RNAase free Eppendorf tubes and centrifuged at 800g after which the supernatant was removed and the cell pellet frozen in liquid nitrogen and stored at −80°C. Following RNA extraction RNA was quality controlled using the Agilent Bioanalyzer 2100. Sequencing libraries were prepared using Takara SMARTer Stranded Total RNA-Seq Kit v2 pico kit. Libraries were quality controlled using the Agilent Bioanalyzer 2100 and quantified using qPCR. Equimolar amounts of libraries were pooled and sequenced on an Illumina HiSeq 4000 lane as Paired end 100bp.

For FAPs stimulated towards adipogenic (ADM + 1% penstrep) and fibrogenic (DMEM+20ng/ml PDGF-AA, 10% FBS, 1% penstrep) differentiation for six days. The cells were then washed in PBS, trypsinized, transferred to RNAase free Eppendorf tubes, centrifuged at 500g and supernatant was removed. The cell pellet was frozen in liquid N2 and stored at −80°C. The library preparation was done using TruSeq® Stranded mRNA Sample preparation kit (Illumina inc). After RNA extraction 100 ng of total RNA was mRNA enriched using the oligodT bead system. The isolated mRNA was subsequently fragmented using enzymatic fragmentation. Then first strand synthesis and second strand synthesis were performed and the double stranded cDNA was purified (AMPure XP, Beckman Coulter). The cDNA was end repaired, 3’ adenylated and Illumina sequencing adaptors ligated onto the fragments ends, and the library was purified (AMPure XP). The mRNA stranded libraries were pre-amplified with PCR (15 cycles) and purified (AMPure XP). The libraries size distribution was validated and quality inspected on a Bioanalyzer 2100 or BioAnalyzer 4200 tapeStation (Agilent Technologies). High quality libraries are pooled based in equimolar concentrations based on the Bioanalyzer Smear Analysis tool (Agilent Technologies). The library pool(s) were quantified using qPCR and optimal concentration of the library pool used to generate the clusters on the surface of a flowcell before sequencing on a NextSeq500 instrument (2×51 cycles, single read) according to the manufacturer instructions (Illumina Inc.).

### Single-cell RNA seq (10x Genomics)

Digestion of four whole human skeletal muscle was performed as described previously and cell suspension was frozen. The cryopreserved samples were thawed in a water bath at 37°C before dilution in 10 ml wash-buffer followed by centrifugation at 500g for 5 min. The samples were resuspended in wash-buffer, stained with antibodies (see FACS section for additional information). Following antibody incubation, the cells were washed in wah-buffer, centrifuged at 500g and resuspended in wash-buffer. The cells were then filtered through a 30 μm cell strainer before sorting. PI was added immediately before sorting. Cells were sorted into DMEM, 10% Horse serum and 1% Penstrep. The PI-negative cells were loaded on the Chromium system (10x Genomics) using manufactures protocol (Chromium Single Cell 3’ Reagent Kits (v3 Chemistry)) for a targeted recovery of 5000 cells. Libraries were sequenced on an Illumina Hiseq4000 and mapped to the human genome (build GRCh38) using Cell Ranger software (10x Genomics, version 3.1.0). Following normalization and quality control, we captured >18000 cells that we included in the analysis with an average of 129000 reads and 1397 identified genes per cell from four biological replicates (Fig S6 A-C, E, F).

### Bioinformatics

#### Bulk RNAseq analysis

For all paired-end reads, trim galore was used for adapter removal and to trim low-quality ends (Babraham Bioinformatics). A Phred33 cut-off score of 20 was applied before adaptor trimming, and all sequences below 20 bp were subsequently removed. The trimmed reads were aligned to the human genome (hg19) by HISAT2 using default settings^75^, and then sorted by gene name using samtools^76^. The overlap between read alignments and genomic annotations were quantified by htseq-count^77^. The counts were then normalised and tested for differential expression (DE) using the R Bioconductor package DESeq2^78^. When the different cell types had been isolated from the same individual, a model design was implemented to correct for inter-person variability. Differentially expressed genes were considered significant when the FDR adjusted p-value were < 0.05 and had a log2fold change > ± 1. For principal component analysis, all gene counts were log2 transformed and was then used as input for principal component analysis by the plotPCA function of DESeq2.

Shared differentially expressed genes (FDR adjusted p-value < 0.05, DESeq2) between the comparisons FAP^CD90+^ vs FAP^CD90−^ and itT2D vs OBS were presented in a Venn plot. Furthermore, the log2fold change of the overlapping genes were presented as a bar plot.

#### Gene ontology and pathway enrichment analysis

The differentially expressed genes were used as input for ontology and pathway enrichment analysis by the R Bioconductor package XGR^79^. Functional pathway analysis on selected pathway gene sets (MsigdbC2CPall) and Gene Ontology (Biological Process) analysis was assessed with a minimum overlap of 3 target genes. The significance was determined using a cut-off of FDR adjusted p-value < 0.05 and visualised according to the adjusted p-value.

Shared genes between the gene ontology term “Extracellular Matrix” (GO:0031012) and genes up-regulated in FAP^CD90+^ compared to FAP^CD90−^ were merged with shared genes between the gene ontology term “Angiogenesis” (GO: 0001525) and genes up-regulated in FAP^CD90−^ compared to FAP^CD90+^. The merged data was presented as a volcano plot created in the R Bioconductor package EnhancedVolcano (Blighe K (2019). *EnhancedVolcano: Publication-ready volcano plots with enhanced colouring and labeling*. R package version 1.2.0)

#### Single cell sequencing analysis

The scRNAseq count data was analysed in R using the Seurat package and workflow^80^. Shortly, cells with unique feature counts above 200 and below 5000, and in addition, had a mitochondrial gene content < 30% were selected. The gene counts for each cell were normalized to the cells’ total gene count, scaled by 10000, and log-transformed. Next, variable features were identified by the FindVariableFeatures function using the “vst” selection method and returning 2000 variable genes per dataset. The gene counts were then scaled and dimensional reduction by principal component analysis taking the variable features as input was performed. The dimensionality of the dataset was identified using the JackStraw procedure and the ElbowPlot function and Seurats graph-based clustering approach, FindNeighbors and FindClusters, was subsequently used to cluster the cells based on the previously determined dimensionality of the dataset. The resolution was set to 0.5 in the FindClusters function. To predict and remove doublets, the pre-processed Seurat object, as prepared above, were used as input for the R package DoubletFinder ^81^. The optimal pK (PC neighbourhood size) and pN (number of generated artificial doublets) values were estimated using a principal component number of 20 and assuming a 7.5% doublet formation rate. The doubletFinder function was then evoked and the predicted doublets were removed from the Seurat object (Fig S6 D). The clusters were then visualized in a Uniform Manifold Approximation and Projection (UMAP) plot. Differential expression analysis by the function FindAllMarkers was used to identify specific cluster biomarkers. As an inclusion criterion, a gene had to be detected in minimum 25% of the cells in either of the compared groups and have at least ±0.25 log2fold difference between the compared groups of cells. The top-10 differentially expressed genes from each cluster were then presented in a heatmap using the DoHeatmap function.

To investigate FAPs in further detail, all cells in the individual FAPs clusters were subsequently extracted, re-processed and analyzed as described above. Furthermore, FAP^CD90+^ and FAP^CD90−^ cells were extracted and the FindAllMarkers function was used to identify differentially expressed genes. Subsequently, the DE genes were used as input for gene set enrichment analysis as described above.

### Statistics

All n’s are true biological replicates (separate biological donors) unless otherwise stated. Statistical tests, including paired/unpaired Student’s t-test (two-sided), one-way ANOVA and non-linear fit analyses were performed using Prism 8 (Version 8.4.1, GraphPad Software). Analysis of limiting dilution data were performed using a web application made available by the Walter and Eliza Hall Institute of Medical Research, Melbourne, Australia (http://bioinf.wehi.edu.au/software/limdil/46)^15^. This software tests departures from the single-hit Poisson model using a generalized linear model. All data, including Supplementary figures, are presented as Violin plots with median and quartiles with superimposed individual values or mean ± SEM.

## REFERENCES

1. Zurlo, F., Larson, K., Bogardus, C. & Ravussin, E. Skeletal muscle metabolism is a major determinant of resting energy expenditure. J Clin Invest 86, 1423–1427 (1990).

2. Baron, A.D., Brechtel, G., Wallace, P. & Edelman, S.V. Rates and tissue sites of non-insulin- and insulin-mediated glucose uptake in humans. Am J Physiol 255, E769–774 (1988).

3. Srikanthan, P. & Karlamangla, A.S. Muscle mass index as a predictor of longevity in older adults. Am J Med 127, 547–553 (2014).

4. Zhao, X., Kwan, J.Y.Y., Yip, K., Liu, P.P. & Liu, F.F. Targeting metabolic dysregulation for fibrosis therapy. Nature reviews. Drug discovery (2019).

5. Morley, J.E. Diabetes and aging: epidemiologic overview. Clin Geriatr Med 24, 395–405, v (2008).

6. Lopez-Otin, C., Blasco, M.A., Partridge, L., Serrano, M. & Kroemer, G. The hallmarks of aging. Cell 153, 1194–1217 (2013).

7. Conte, M., Martucci, M., Sandri, M., Franceschi, C. & Salvioli, S. The Dual Role of the Pervasive “Fattish” Tissue Remodeling With Age. Frontiers in Endocrinology 10(2019).

8. Moore, C.W., Allen, M.D., Kimpinski, K., Doherty, T.J. & Rice, C.L. Reduced skeletal muscle quantity and quality in patients with diabetic polyneuropathy assessed by magnetic resonance imaging. Muscle Nerve 53, 726–732 (2016).

9. Addison, O., Marcus, R.L., Lastayo, P.C. & Ryan, A.S. Intermuscular Fat: A Review of the Consequences and Causes. International journal of endocrinology 2014, 309570 (2014).

10. Goodpaster, B.H., Thaete, F.L. & Kelley, D.E. Thigh adipose tissue distribution is associated with insulin resistance in obesity and in type 2 diabetes mellitus. Am J Clin Nutr 71, 885–892 (2000).

11. Buras, E.D., et al. Fibro-Adipogenic Remodeling of the Diaphragm in Obesity-Associated Respiratory Dysfunction. Diabetes 68, 45–56 (2019).

12. Kang, L., et al. Diet-induced muscle insulin resistance is associated with extracellular matrix remodeling and interaction with integrin alpha2beta1 in mice. Diabetes 60, 416–426 (2011).

13. Inoue, M., et al. Thrombospondin 1 mediates high-fat diet-induced muscle fibrosis and insulin resistance in male mice. Endocrinology 154, 4548–4559 (2013).

14. Uezumi, A., Fukada, S., Yamamoto, N., Takeda, S. & Tsuchida, K. Mesenchymal progenitors distinct from satellite cells contribute to ectopic fat cell formation in skeletal muscle. Nat Cell Biol 12, 143–152 (2010).

15. Joe, A.W., et al. Muscle injury activates resident fibro/adipogenic progenitors that facilitate myogenesis. Nat Cell Biol 12, 153–163 (2010).

16. Lemos, D.R., et al. Nilotinib reduces muscle fibrosis in chronic muscle injury by promoting TNF-mediated apoptosis of fibro/adipogenic progenitors. Nat Med 21, 786–794 (2015).

17. Mozzetta, C., et al. Fibroadipogenic progenitors mediate the ability of HDAC inhibitors to promote regeneration in dystrophic muscles of young, but not old Mdx mice. EMBO Mol Med 5, 626–639 (2013).

18. Mueller, A.A., van Velthoven, C.T., Fukumoto, K.D., Cheung, T.H. & Rando, T.A. Intronic polyadenylation of PDGFRalpha in resident stem cells attenuates muscle fibrosis. Nature 540, 276–279 (2016).

19. Fiore, D., et al. Pharmacological blockage of fibro/adipogenic progenitor expansion and suppression of regenerative fibrogenesis is associated with impaired skeletal muscle regeneration. Stem cell research 17, 161–169 (2016).

20. Dong, Y., Silva, K.A., Dong, Y. & Zhang, L. Glucocorticoids increase adipocytes in muscle by affecting IL-4 regulated FAP activity. FASEB J (2014).

21. Heredia, J.E., et al. Type 2 innate signals stimulate fibro/adipogenic progenitors to facilitate muscle regeneration. Cell 153, 376–388 (2013).

22. Marcelin, G., et al. A PDGFRα-Mediated Switch toward CD9high Adipocyte Progenitors Controls Obesity-Induced Adipose Tissue Fibrosis. Cell Metabolism (2017).

23. Iwayama, T., et al. PDGFRalpha signaling drives adipose tissue fibrosis by targeting progenitor cell plasticity. Genes Dev 29, 1106–1119 (2015).

24. Uezumi, A., et al. Identification and characterization of PDGFRalpha+ mesenchymal progenitors in human skeletal muscle. Cell death & disease 5, e1186 (2014).

25. Arrighi, N., et al. Characterization of adipocytes derived from fibro/adipogenic progenitors resident in human skeletal muscle. Cell death & disease 6, e1733 (2015).

26. Agley, C.C., Rowlerson, A.M., Velloso, C.P., Lazarus, N.R. & Harridge, S.D. Human skeletal muscle fibroblasts, but not myogenic cells, readily undergo adipogenic differentiation. J Cell Sci 126, 5610–5625 (2013).

27. Moller, A.B., et al. Altered gene expression and repressed markers of autophagy in skeletal muscle of insulin resistant patients with type 2 diabetes. Scientific reports 7, 43775 (2017).

28. Kampmann, U., et al. GLUT4 and UBC9 protein expression is reduced in muscle from type 2 diabetic patients with severe insulin resistance. PLoS One 6, e27854 (2011).

29. Kampmann, U., et al. Insulin dose-response studies in severely insulin-resistant type 2 diabetes--evidence for effectiveness of very high insulin doses. Diabetes Obes Metab 13, 511–516 (2011).

30. Wosczyna, M.N., et al. Mesenchymal Stromal Cells Are Required for Regeneration and Homeostatic Maintenance of Skeletal Muscle. Cell reports 27, 2029–2035 e2025 (2019).

31. Contreras, O., et al. Cross-talk between TGF-beta and PDGFRalpha signaling pathways regulates the fate of stromal fibro-adipogenic progenitors. J Cell Sci 132(2019).

32. Madaro, L., et al. Denervation-activated STAT3-IL-6 signalling in fibro-adipogenic progenitors promotes myofibres atrophy and fibrosis. Nat Cell Biol 20, 917–927 (2018).

33. Crisan, M., et al. A perivascular origin for mesenchymal stem cells in multiple human organs. Cell Stem Cell 3, 301–313 (2008).

34. Wosczyna, M.N., Biswas, A.A., Cogswell, C.A. & Goldhamer, D.J. Multipotent progenitors resident in the skeletal muscle interstitium exhibit robust BMP-dependent osteogenic activity and mediate heterotopic ossification. J Bone Miner Res 27, 1004–1017 (2012).

35. Shore, E.M. & Kaplan, F.S. Insights from a rare genetic disorder of extra-skeletal bone formation, fibrodysplasia ossificans progressiva (FOP). Bone 43, 427–433 (2008).

36. Cushner, F.D. & Morwessel, R.M. Myositis ossificans traumatica. Orthop Rev 21, 1319–1326 (1992).

37. Giordani, L., et al. High-Dimensional Single-Cell Cartography Reveals Novel Skeletal Muscle-Resident Cell Populations. Mol Cell (2019).

38. Porichis, F., et al. High-throughput detection of miRNAs and gene-specific mRNA at the single-cell level by flow cytometry. Nature communications 5, 5641 (2014).

39. Lukjanenko, L., et al. Aging Disrupts Muscle Stem Cell Function by Impairing Matricellular WISP1 Secretion from Fibro-Adipogenic Progenitors. Cell Stem Cell (2019).

40. de Morree, A., et al. Alternative polyadenylation of Pax3 controls muscle stem cell fate and muscle function. Science 366, 734–738 (2019).

41. Tabula Muris, C., et al. Single-cell transcriptomics of 20 mouse organs creates a Tabula Muris. Nature 562, 367–372 (2018).

42. Mackey, A.L., Magnan, M., Chazaud, B. & Kjaer, M. Human skeletal muscle fibroblasts stimulate in vitro myogenesis and in vivo muscle regeneration. J Physiol (2017).

43. Ieronimakis, N., et al. PDGFRalpha signalling promotes fibrogenic responses in collagen-producing cells in Duchenne muscular dystrophy. J Pathol 240, 410–424 (2016).

44. Olson, L.E. & Soriano, P. Increased PDGFRα Activation Disrupts Connective Tissue Development and Drives Systemic Fibrosis. Developmental Cell 16, 303–313 (2009).

45. Xiao, Y., et al. PDGF Promotes the Warburg Effect in Pulmonary Arterial Smooth Muscle Cells via Activation of the PI3K/AKT/mTOR/HIF-1alpha Signaling Pathway. Cellular physiology and biochemistry : international journal of experimental cellular physiology, biochemistry, and pharmacology 42, 1603–1613 (2017).

46. Derosa, G., Sahebkar, A. & Maffioli, P. The role of various peroxisome proliferator-activated receptors and their ligands in clinical practice. J Cell Physiol 233, 153–161 (2018).

47. Pawlak, M., Lefebvre, P. & Staels, B. Molecular mechanism of PPARalpha action and its impact on lipid metabolism, inflammation and fibrosis in non-alcoholic fatty liver disease. J Hepatol 62, 720–733 (2015).

48. Walker, J.T., McLeod, K., Kim, S., Conway, S.J. & Hamilton, D.W. Periostin as a multifunctional modulator of the wound healing response. Cell Tissue Res 365, 453–465 (2016).

49. Scott, R.W., Arostegui, M., Schweitzer, R., Rossi, F.M.V. & Underhill, T.M. Hic1 Defines Quiescent Mesenchymal Progenitor Subpopulations with Distinct Functions and Fates in Skeletal Muscle Regeneration. Cell Stem Cell 25, 797–813 e799 (2019).

50. Schwalie, P.C., et al. A stromal cell population that inhibits adipogenesis in mammalian fat depots. Nature (2018).

51. Merrick, D., et al. Identification of a mesenchymal progenitor cell hierarchy in adipose tissue. Science 364(2019).

52. Malecova, B., et al. Dynamics of cellular states of fibro-adipogenic progenitors during myogenesis and muscular dystrophy. Nature communications 9(2018).

53. Soliman, H., et al. Pathogenic Potential of Hic1-Expressing Cardiac Stromal Progenitors. Cell Stem Cell 26, 205–220 e208 (2020).

54. Abderrahmani, A., et al. Increased Hepatic PDGF-AA Signaling Mediates Liver Insulin Resistance in Obesity-Associated Type 2 Diabetes. Diabetes 67, 1310–1321 (2018).

55. Sundelin, E., et al. Genetic Polymorphisms in Organic Cation Transporter 1 Attenuates Hepatic Metformin Exposure in Humans. Clin Pharmacol Ther 102, 841–848 (2017).

56. Zhou, G., et al. Role of AMP-activated protein kinase in mechanism of metformin action. J Clin Invest 108, 1167–1174 (2001).

57. Ahlqvist, E., et al. Novel subgroups of adult-onset diabetes and their association with outcomes: a data-driven cluster analysis of six variables. The Lancet Diabetes & Endocrinology 6, 361–369 (2018).

58. Rasmussen, D.G.K., et al. Higher Collagen VI Formation Is Associated With All-Cause Mortality in Patients With Type 2 Diabetes and Microalbuminuria. Diabetes Care 41, 1493–1500 (2018).

59. Miljkovic, I., et al. Greater adipose tissue infiltration in skeletal muscle among older men of African ancestry. J Clin Endocrinol Metab 94, 2735–2742 (2009).

60. Volpato, S., et al. Role of muscle mass and muscle quality in the association between diabetes and gait speed. Diabetes Care 35, 1672–1679 (2012).

61. Almurdhi, M.M., et al. Reduced Lower-Limb Muscle Strength and Volume in Patients With Type 2 Diabetes in Relation to Neuropathy, Intramuscular Fat, and Vitamin D Levels. Diabetes Care 39, 441–447 (2016).

62. Li, J.J., et al. Muscle grip strength predicts incident type 2 diabetes: Population-based cohort study. Metabolism 65, 883–892 (2016).

63. Yeung, C.H.C., Au Yeung, S.L., Fong, S.S.M. & Schooling, C.M. Lean mass, grip strength and risk of type 2 diabetes: a bi-directional Mendelian randomisation study. Diabetologia 62, 789–799 (2019).

64. Nomura, T., Kawae, T., Kataoka, H. & Ikeda, Y. Aging, physical activity, and diabetic complications related to loss of muscle strength in patients with type 2 diabetes. Phys Ther Res 21, 33–38 (2018).

65. Murai, J., et al. Low muscle quality in Japanese type 2 diabetic patients with visceral fat accumulation. Cardiovasc Diabetol 17, 112 (2018).

66. Warram, J.H., Martin, B.C., Krolewski, A.S., Soeldner, J.S. & Kahn, C.R. Slow glucose removal rate and hyperinsulinemia precede the development of type II diabetes in the offspring of diabetic parents. Ann Intern Med 113, 909–915 (1990).

67. de Paz-Lugo, P., Lupianez, J.A. & Melendez-Hevia, E. High glycine concentration increases collagen synthesis by articular chondrocytes in vitro: acute glycine deficiency could be an important cause of osteoarthritis. Amino Acids 50, 1357–1365 (2018).

68. Im, M.J., Freshwater, M.F. & Hoopes, J.E. Enzyme activities in granulation tissue: Energy for collagen synthesis. J Surg Res 20, 121–125 (1976).

69. Ran, C., Liu, H., Hitoshi, Y. & Israel, M.A. Proliferation-independent control of tumor glycolysis by PDGFR-mediated AKT activation. Cancer Res 73, 1831–1843 (2013).

70. Pala, F., et al. Distinct metabolic states govern skeletal muscle stem cell fates during prenatal and postnatal myogenesis. J Cell Sci 131(2018).

71. Rodgers, J.T., et al. mTORC1 controls the adaptive transition of quiescent stem cells from G0 to GAlert. Nature (2014).

72. Mahmoudi, S., et al. Heterogeneity in old fibroblasts is linked to variability in reprogramming and wound healing. Nature 574, 553–558 (2019).

73. Gurtler, A., et al. Stain-Free technology as a normalization tool in Western blot analysis. Anal Biochem 433, 105–111 (2013).

74. Vigelso, A., et al. GAPDH and beta-actin protein decreases with aging, making Stain-Free technology a superior loading control in Western blotting of human skeletal muscle. J Appl Physiol (1985) 118, 386–394 (2015).

75. Kim, D., Langmead, B. & Salzberg, S.L. HISAT: a fast spliced aligner with low memory requirements. Nat Methods 12, 357–360 (2015).

76. Li, H., et al. The Sequence Alignment/Map format and SAMtools. Bioinformatics 25, 2078–2079 (2009).

77. Anders, S., Pyl, P.T. & Huber, W. HTSeq--a Python framework to work with high-throughput sequencing data. Bioinformatics 31, 166–169 (2015).

78. Love, M.I., Huber, W. & Anders, S. Moderated estimation of fold change and dispersion for RNA-seq data with DESeq2. Genome Biol 15, 550 (2014).

79. Fang, H., Knezevic, B., Burnham, K.L. & Knight, J.C. XGR software for enhanced interpretation of genomic summary data, illustrated by application to immunological traits. Genome Med 8, 129 (2016).

80. Stuart, T., et al. Comprehensive Integration of Single-Cell Data. Cell 177, 1888–1902 e1821 (2019).

81. McGinnis, C.S., Murrow, L.M. & Gartner, Z.J. DoubletFinder: Doublet Detection in Single-Cell RNA Sequencing Data Using Artificial Nearest Neighbors. Cell Syst 8, 329–337 e324 (2019).

