## Supplementary data for "Human skeletal muscle CD90^+^ fibro-adipogenic progenitors are associated with muscle degeneration in type 2 diabetic patients"

**Figure S1**

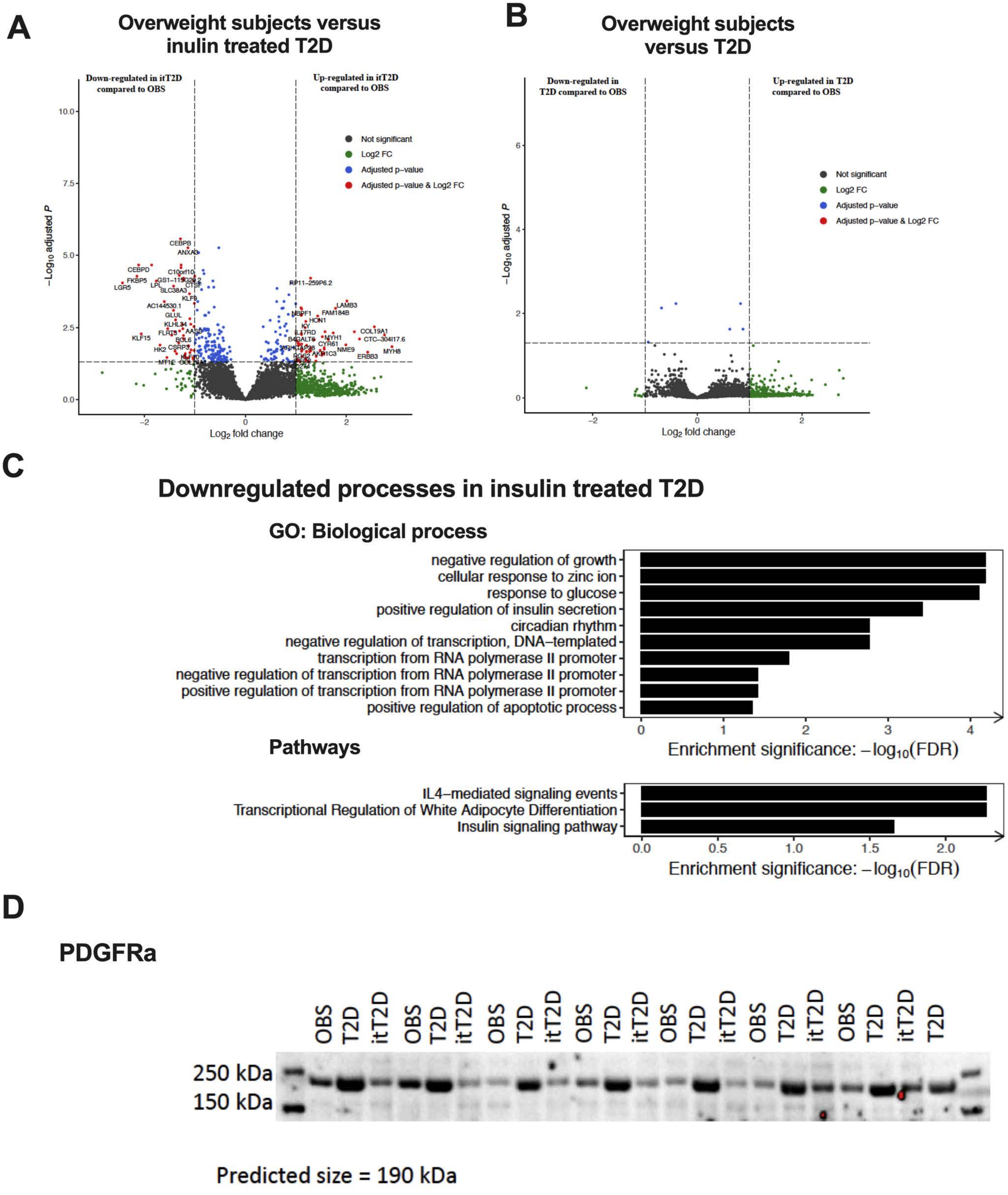
A
Overweight subjects versus inulin treated T2D
Down-regulated in itT2D compared to OBS
Up-regulated in itT2D compared to OBS
Not significant
Log2 FC
Adjusted p–value
Adjusted p–value & Log2 FC
$-\log_{10}$ adjusted $P$
$\log_2$ fold change
CEBPB
ANXA3
CEBPD
FKBP5
LGR5
LPL
SLC38A3
KLF9
GLUL
KLF15
LAMB3
FAM184B
COL19A1
MYH8
ERBB3
NME9
B
Overweight subjects versus T2D
Down-regulated in T2D compared to OBS
Up-regulated in T2D compared to OBS
Not significant
Log2 FC
Adjusted p–value
Adjusted p–value & Log2 FC
$-\log_{10}$ adjusted $P$
$\log_2$ fold change
C
Downregulated processes in insulin treated T2D
GO: Biological process
negative regulation of growth
cellular response to zinc ion
response to glucose
positive regulation of insulin secretion
circadian rhythm
negative regulation of transcription, DNA–templated
transcription from RNA polymerase II promoter
negative regulation of transcription from RNA polymerase II promoter
positive regulation of transcription from RNA polymerase II promoter
positive regulation of apoptotic process
Enrichment significance: $-\log_{10}$(FDR)
Pathways
IL4–mediated signaling events
Transcriptional Regulation of White Adipocyte Differentiation
Insulin signaling pathway
Enrichment significance: $-\log_{10}$(FDR)
D
PDGFRa
OBS T2D itT2D OBS T2D itT2D OBS T2D itT2D OBS T2D itT2D OBS T2D itT2D OBS T2D itT2D OBS T2D itT2D T2D
250 kDa
150 kDa
Predicted size = 190 kDa

**Figure S1**

**A,** Volcano plot displaying genes differentially expressed in skeletal muscle from insulin treated type 2 diabetes (itT2D, n=6) compared to overweight subjects (OBS, n=7); **B**, Volcano plot displaying genes differentially expressed in skeletal muscle from type 2 diabetes (T2D, n=7) compared to OBS; **C**, Down-regulated GO: Biological processes and pathways in itT2D compared to OBS; **D**, Western blot of PDGFRα protein in all subjects/patients.

### Figure S2

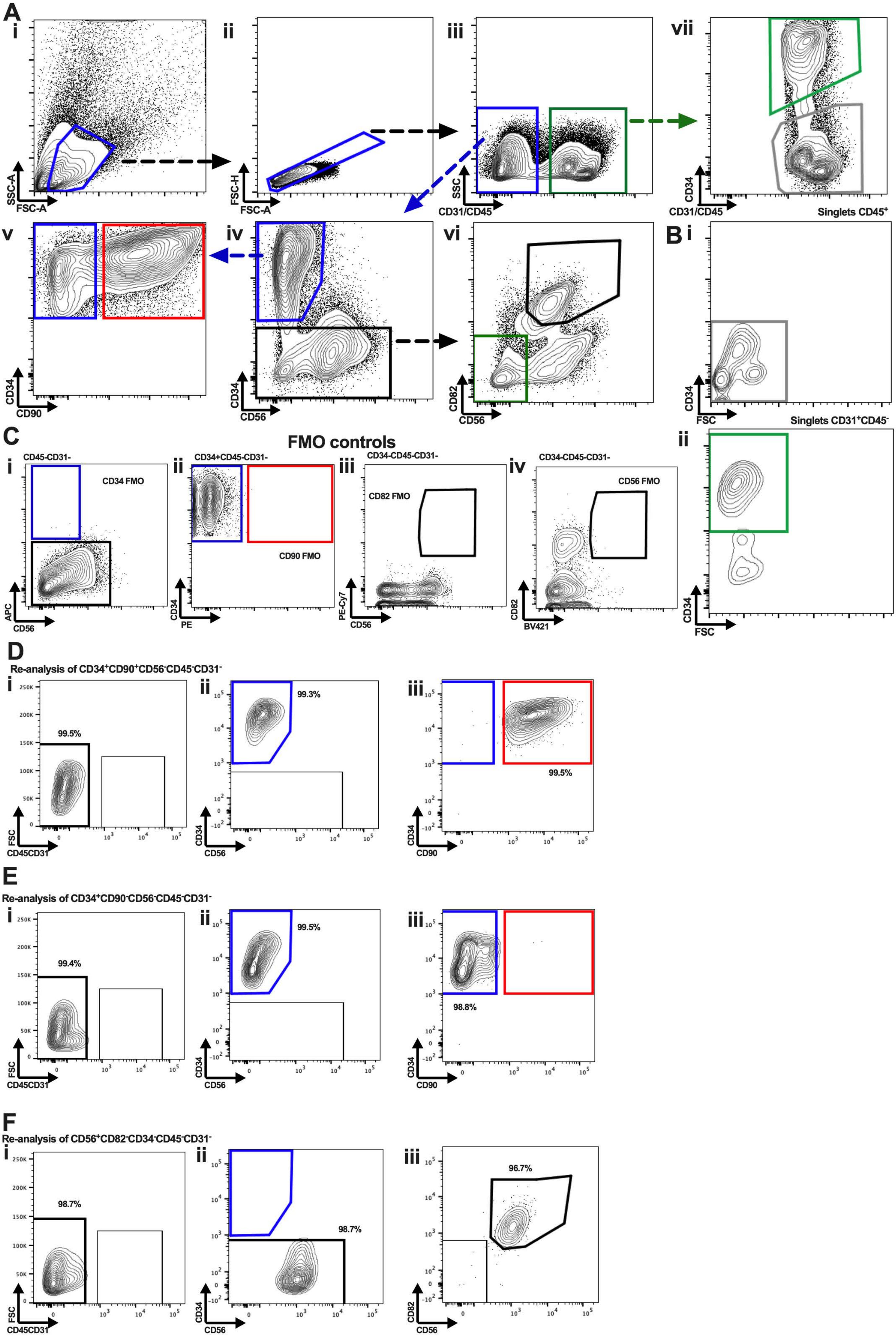
A
i
ii
iii
vii
SSC-A
FSC-A
FSC-H
FSC-A
SSC
CD31/CD45
CD34
CD31/CD45
Singlets CD45+
v
iv
vi
B i
CD34
CD90
CD34
CD56
CD82
CD56
CD34
FSC
Singlets CD31+CD45-
C
FMO controls
ii
CD45-CD31-
CD34 FMO
CD34+CD45-CD31-
CD90 FMO
CD34-CD45-CD31-
CD82 FMO
CD34-CD45-CD31-
CD56 FMO
APC
CD56
CD34
PE
PE-Cy7
CD56
CD82
BV421
CD34
FSC
D
Re-analysis of CD34+CD90+CD56-CD45-CD31-
99.5%
99.3%
99.5%
FSC
CD45CD31
CD34
CD56
CD34
CD90
E
Re-analysis of CD34+CD90-CD56-CD45-CD31-
99.4%
99.5%
98.8%
FSC
CD45CD31
CD34
CD56
CD34
CD90
F
Re-analysis of CD56+CD82-CD34-CD45-CD31-
98.7%
98.7%
96.7%
FSC
CD45CD31
CD34
CD56
CD82
CD56

**Figure S2**

**A,** Gating strategy identifying (**i**) Cells, (**ii**) Single cells, (**iii**) $CD45CD31^-$ and $CD45CD31^+$ cells, (**iv**) $CD34^+CD56^-$ and $CD34^-$ cells, (**v**) $CD34^+CD90^-$ and $CD34^+CD90^+$ Fibro-adipogenic progenitors (FAPs), (**vi**) $CD56^+CD82^+$ Muscle stem cells (MuSCs) and $CD56^-CD82^-$ cells, (**vii**) $CD34^+$ endothelial cells and $CD34^-$ hematopoietic cells; **B**, Gating strategy displaying (**i**) $CD34^+$ cells in the $CD45^+$ cell population and (**ii**) $CD34^+$ cells in the $CD31^+$ cell population; **C**, Fluorescence minus one (FMO) controls for key cell populations including (**i**) CD34 FMO, (**ii**) CD90 FMO, (**iii**) CD82 FMO and (**iv**) CD56 FMO; **D i-iii**, Re-analysis of FACS sorted $CD34^+CD90^+CD56^-CD45^-CD31^-$ cells; **E i-iii,** Re-analysis of FACS sorted $CD34^+CD90^-CD56^-CD45^-CD31^-$ cells; **F i-iii,** Re-analysis of FACS sorted $CD56^+CD82^+CD34^-CD45^-CD31^-$ cells.

### Figure S3

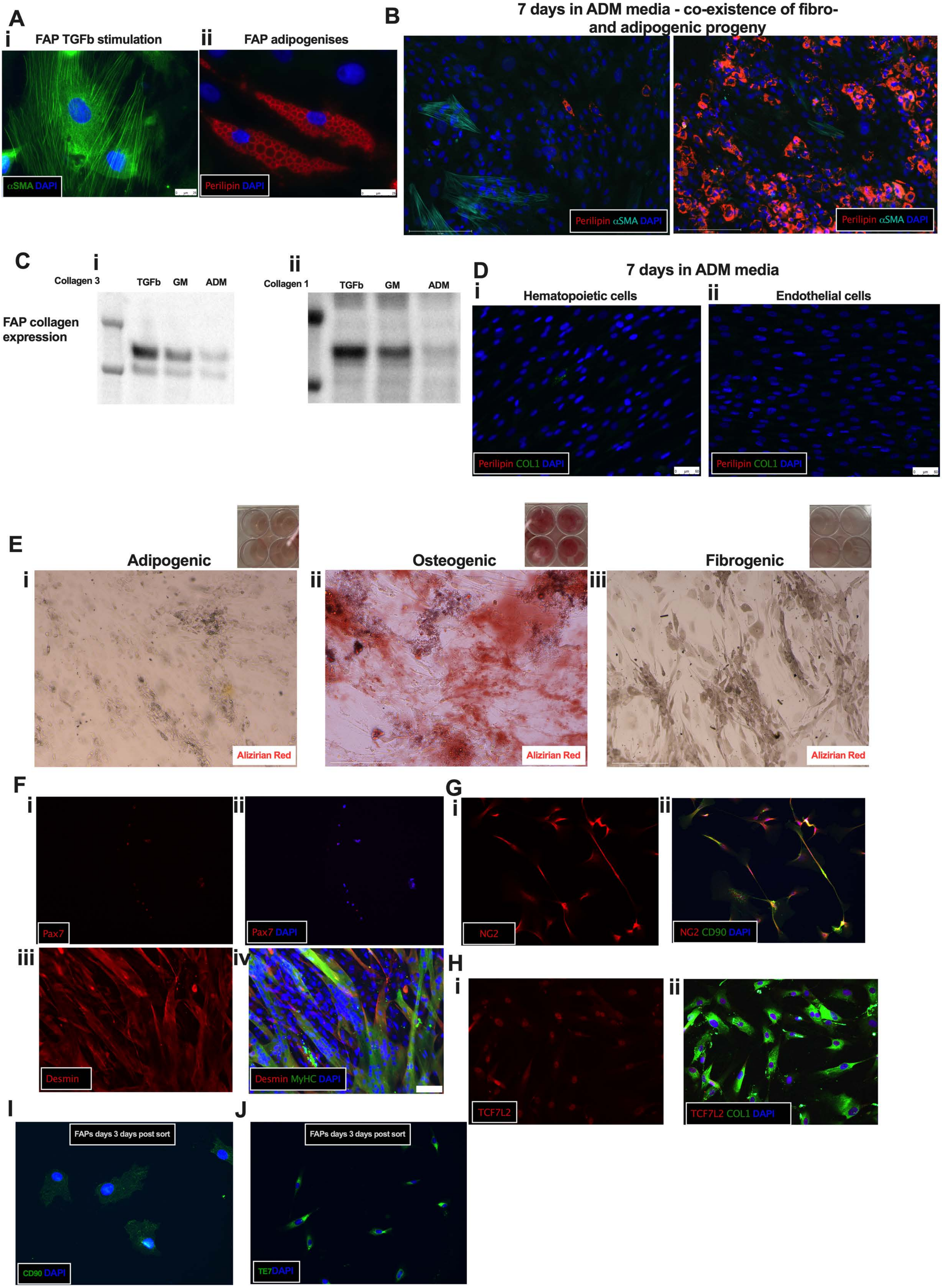
A
i FAP TGFb stimulation
ii FAP adipogenises
αSMA DAPI
Perilipin DAPI
B
7 days in ADM media - co-existence of fibro- and adipogenic progeny
Perilipin αSMA DAPI
Perilipin αSMA DAPI
C
i Collagen 3
TGFb GM ADM
ii Collagen 1
TGFb GM ADM
FAP collagen expression
D
7 days in ADM media
i Hematopoietic cells
ii Endothelial cells
Perilipin COL1 DAPI
Perilipin COL1 DAPI
E
i Adipogenic
ii Osteogenic
iii Fibrogenic
Alizirian Red
Alizirian Red
Alizirian Red
F
i Pax7
ii Pax7 DAPI
iii Desmin
iv Desmin MyHC DAPI
G
i NG2
ii NG2 CD90 DAPI
H
i TCF7L2
ii TCF7L2 COL1 DAPI
I
FAPs days 3 days post sort
CD90 DAPI
J
FAPs days 3 days post sort
TE7DAPI

**Figure S3**

**A**, FACS isolated FAPs ($CD34^+CD56^-CD31^-CD45^-$) stimulated with (**i**) TGFβ (1ng/ml) for 3 days towards fibrogenesis and stained with alpha smooth muscle actin (aSMA) or (**ii**) adipogenic induction media towards adipogenesis and stained with Perilipin-1; **B**, Co-existence of Perilipin-$1^+$ (red) adipocytes and $aSMA^+$ (Cyan) fibroblasts in FAP culture; **C**, Western blotting of Collagen-3 (**i**) and Collagen1 (**ii**) protein expression in FAPs ($CD34^+CD56^-CD31^-CD45^-$) stimulated with TGFβ (1ng/ml), growth media or adipogenic induction media; **D**, (**i**) hematopoietic or (**ii**) endothelial cells isolated from skeletal muscle and stimulated for seven days towards adipogenesis and stained with Perilipin-1 and Collagen-1; **E**, FAPs ($CD34^+CD90^+CD56^-CD31^-CD45^-$) stimulated for 10 days with (**i**) adipogenic, (**ii**) osteogenic or fibrogenic (**iii**) media and stained with Alizarin Red to mark calcium depots; **F**, FACS isolated MuSCs stained with (**i-ii**) Pax7 (red) 24 hours post isolation and (**iii-iv**) Myosin Heavy Chain (MyHC, green) and Desmin (red) after three days of differentiation; **G i-ii**, FACS isolated $CD90^+CD34^-CD56^-CD45^-CD31^-$ cells from skeletal muscle, expanded in growth media and stained for NG2 and CD90; **H i-ii**, FACS isolated FAPs ($CD34^+CD56^-CD31^-CD45^-$) stained after three days in culture with TCF7L2 (red) and Collagen-1 (green); **I**, CD90 (green) staining of sorted $CD34^+CD90^+$ FAPs 3 days post sort; **J**, TE7 staining (green) staining of sorted $CD34^+CD90^+$ FAPs 3 days post sort.

**Figure S4**

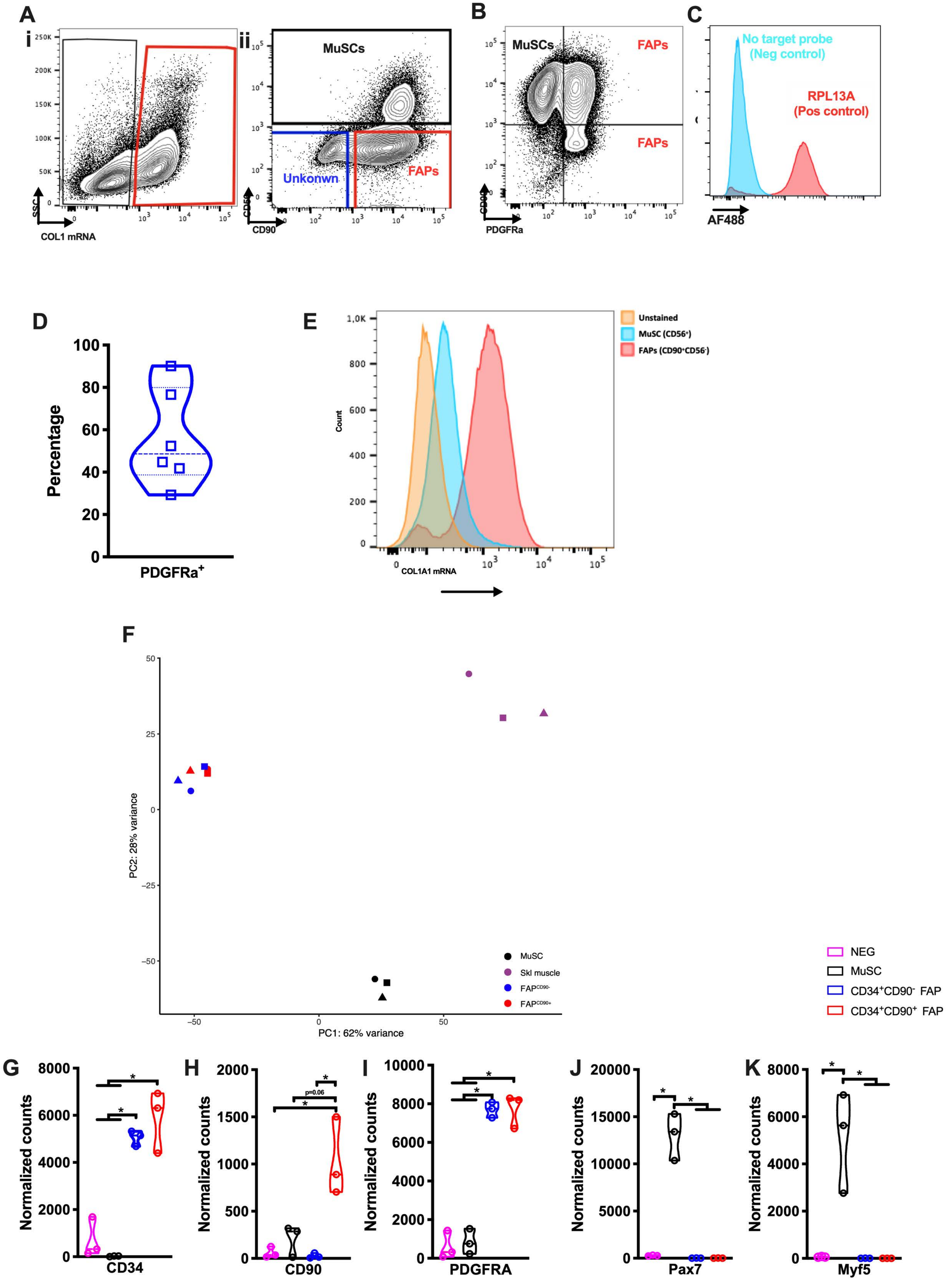

A
i
ii
MuSCs
Unkonwn
FAPs
COL1 mRNA
CD90
B
MuSCs
FAPs
FAPs
PDGFRa
C
No target probe
(Neg control)
RPL13A
(Pos control)
AF488
D
Percentage
PDGFRa+
E
Unstained
MuSC (CD56+)
FAPs (CD90+CD56-)
Count
COL1A1 mRNA
F
PC2: 28% variance
PC1: 62% variance
MuSC
Skl muscle
FAPCD90-
FAPCD90+
NEG
MuSC
CD34+CD90- FAP
CD34+CD90+ FAP
G
H
I
J
K
Normalized counts
p=0.06
CD34
CD90
PDGFRA
Pax7
Myf5

**Figure S4**

**A**, Gating strategy from in situ hybridization flow assay with identification of (**i**) COL1A1 mRNA expressing cells and from this the fraction of (**ii**) MuSC ($CD56^{+}$), FAPs ($CD90^{+}CD56^{-}$) and unknown ($CD90^{-}CD56^{-}$) cells; **B**, Gating strategy of COL1A1 mRNA expressing cells that are FAPs ($PDGFR\alpha^{+}$) or MuSCs ($CD90^{+}PDGFR\alpha^{-}$); **C**, Control probes for in situ RNA flow experiment including positive control (RPL13A) and negative control (omission of target probes); **D**, Percentage of *COL1A1* mRNA expressing staining positive of PDGFRa in human skeletal muscle (n=6) after activation for 3-9 days in vitro; **E**, *COL1A1* mRNA expression (median fluorescence intensity, MFI) in $CD90^{+}CD56^{-}$ (Fibro-adipogenic progenitors; FAPs), $CD56^{+}$ (Muscle stem cells; MuSCs) and $CD90^{-}CD56^{-}$ (Un-identified) cells in human skeletal muscle (n=6) after activation for 3-9 days in vitro; **F**, Principal component analysis-plot from RNA-seq (n=3) on freshly sorted FAPs ($CD90^{-}$ and $CD90^{-}$), MuSCs and Skeletal muscle (Neg); Expression of; **G**, *CD34*; **H**, *THY1*/CD90; **I**, *PDGFRA*; **J**, *PAX7*; **K**, *MYF5* in FAPs, MuSCs and Skeletal muscle cells.

Figure S5

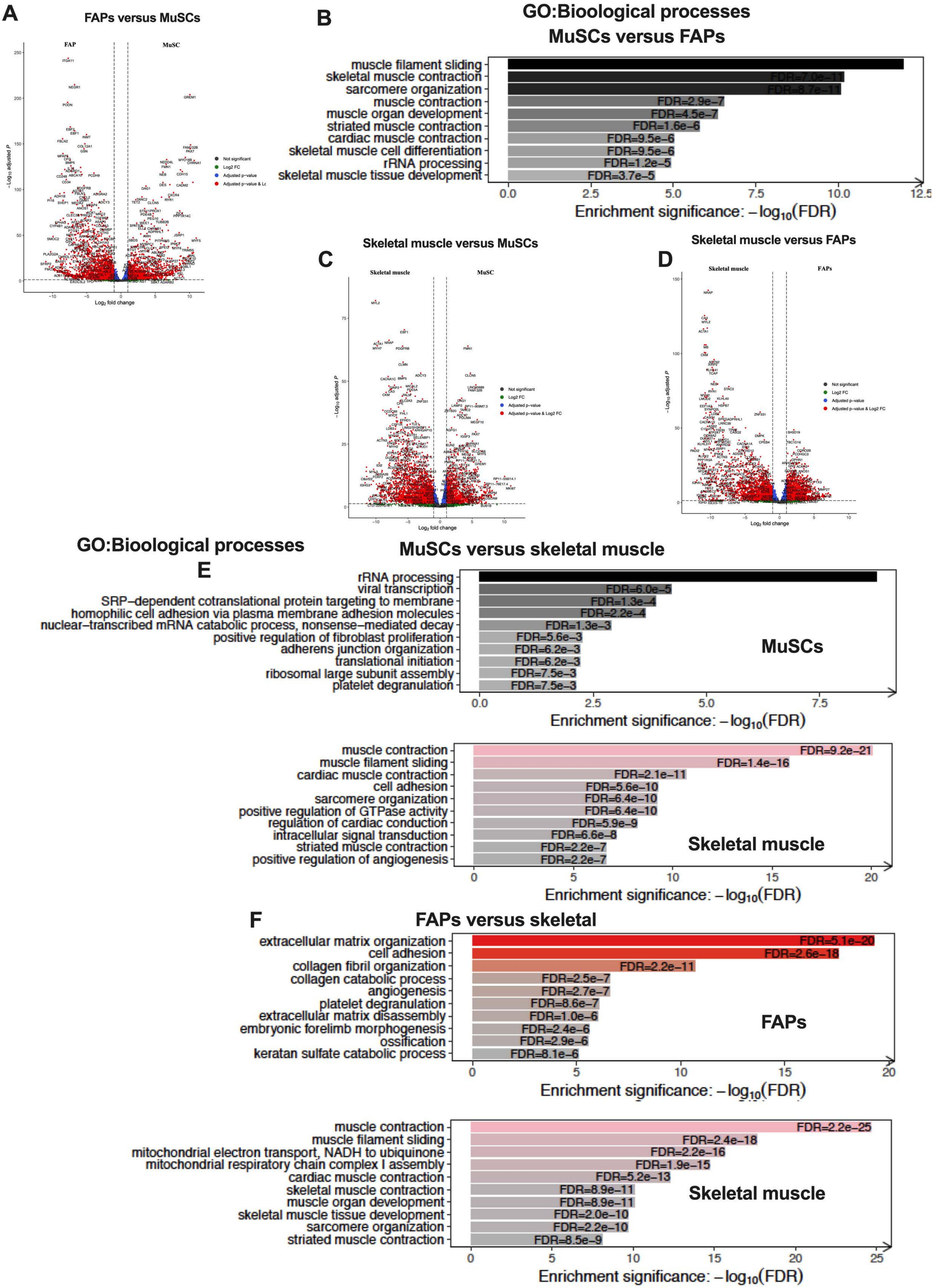
A
FAPs versus MuSCs
FAP
MuSC
−Log10 adjusted P
Log2 fold change
Not significant
Log2 FC
Adjusted p-value
B
GO:Bioological processes
MuSCs versus FAPs
muscle filament sliding
skeletal muscle contraction
sarcomere organization
muscle contraction
muscle organ development
striated muscle contraction
cardiac muscle contraction
skeletal muscle cell differentiation
rRNA processing
skeletal muscle tissue development
FDR=7.0e−11
FDR=8.7e−11
FDR=2.9e−7
FDR=4.5e−7
FDR=1.6e−6
FDR=9.5e−6
FDR=9.5e−6
FDR=1.2e−5
FDR=3.7e−5
Enrichment significance: −log10(FDR)
C
Skeletal muscle versus MuSCs
Skeletal muscle
MuSC
Not significant
Log2 FC
Adjusted p-value
Adjusted p-value & Log2 FC
D
Skeletal muscle versus FAPs
Skeletal muscle
FAPs
GO:Bioological processes
MuSCs versus skeletal muscle
E
rRNA processing
viral transcription
SRP−dependent cotranslational protein targeting to membrane
homophilic cell adhesion via plasma membrane adhesion molecules
nuclear−transcribed mRNA catabolic process, nonsense−mediated decay
positive regulation of fibroblast proliferation
adherens junction organization
translational initiation
ribosomal large subunit assembly
platelet degranulation
FDR=6.0e−5
FDR=1.3e−4
FDR=2.2e−4
FDR=1.3e−3
FDR=5.6e−3
FDR=6.2e−3
FDR=6.2e−3
FDR=7.5e−3
FDR=7.5e−3
MuSCs
Enrichment significance: −log10(FDR)
muscle contraction
muscle filament sliding
cardiac muscle contraction
cell adhesion
sarcomere organization
positive regulation of GTPase activity
regulation of cardiac conduction
intracellular signal transduction
striated muscle contraction
positive regulation of angiogenesis
FDR=9.2e−21
FDR=1.4e−16
FDR=2.1e−11
FDR=5.6e−10
FDR=6.4e−10
FDR=6.4e−10
FDR=5.9e−9
FDR=6.6e−8
FDR=2.2e−7
FDR=2.2e−7
Skeletal muscle
Enrichment significance: −log10(FDR)
F
FAPs versus skeletal
extracellular matrix organization
cell adhesion
collagen fibril organization
collagen catabolic process
angiogenesis
platelet degranulation
extracellular matrix disassembly
embryonic forelimb morphogenesis
ossification
keratan sulfate catabolic process
FDR=5.1e−20
FDR=2.6e−18
FDR=2.2e−11
FDR=2.5e−7
FDR=2.7e−7
FDR=8.6e−7
FDR=1.0e−6
FDR=2.4e−6
FDR=2.9e−6
FDR=8.1e−6
FAPs
Enrichment significance: −log10(FDR)
muscle contraction
muscle filament sliding
mitochondrial electron transport, NADH to ubiquinone
mitochondrial respiratory chain complex I assembly
cardiac muscle contraction
skeletal muscle contraction
muscle organ development
skeletal muscle tissue development
sarcomere organization
striated muscle contraction
FDR=2.2e−25
FDR=2.4e−18
FDR=2.2e−16
FDR=1.9e−15
FDR=5.2e−13
FDR=8.9e−11
FDR=8.9e−11
FDR=2.0e−10
FDR=2.2e−10
FDR=8.5e−9
Skeletal muscle
Enrichment significance: −log10(FDR)

**Figure S5**

**A,** Volcano plot displaying genes differentially expressed in FACS isolated FAPs versus MuSCs; **B**, Volcano plot displaying genes differentially expressed in FACS isolated MuSCs versus mature skeletal muscle (fragments); **C**, Volcano plot displaying genes differentially expressed in FACS isolated FAPs versus mature skeletal muscle (fragments); **D**, GO: Biological processes enriched in FACS isolated muscle stem cells (MuSCs) compared to fibro-adipogenic progenitors (FAPs); **E**, GO: Biological processes enriched in FACS isolated MuSCs versus mature skeletal muscle (fragments); **F,** GO: Biological processes enriched in FACS isolated FAPs versus mature skeletal muscle (fragments).

#### Figure S6

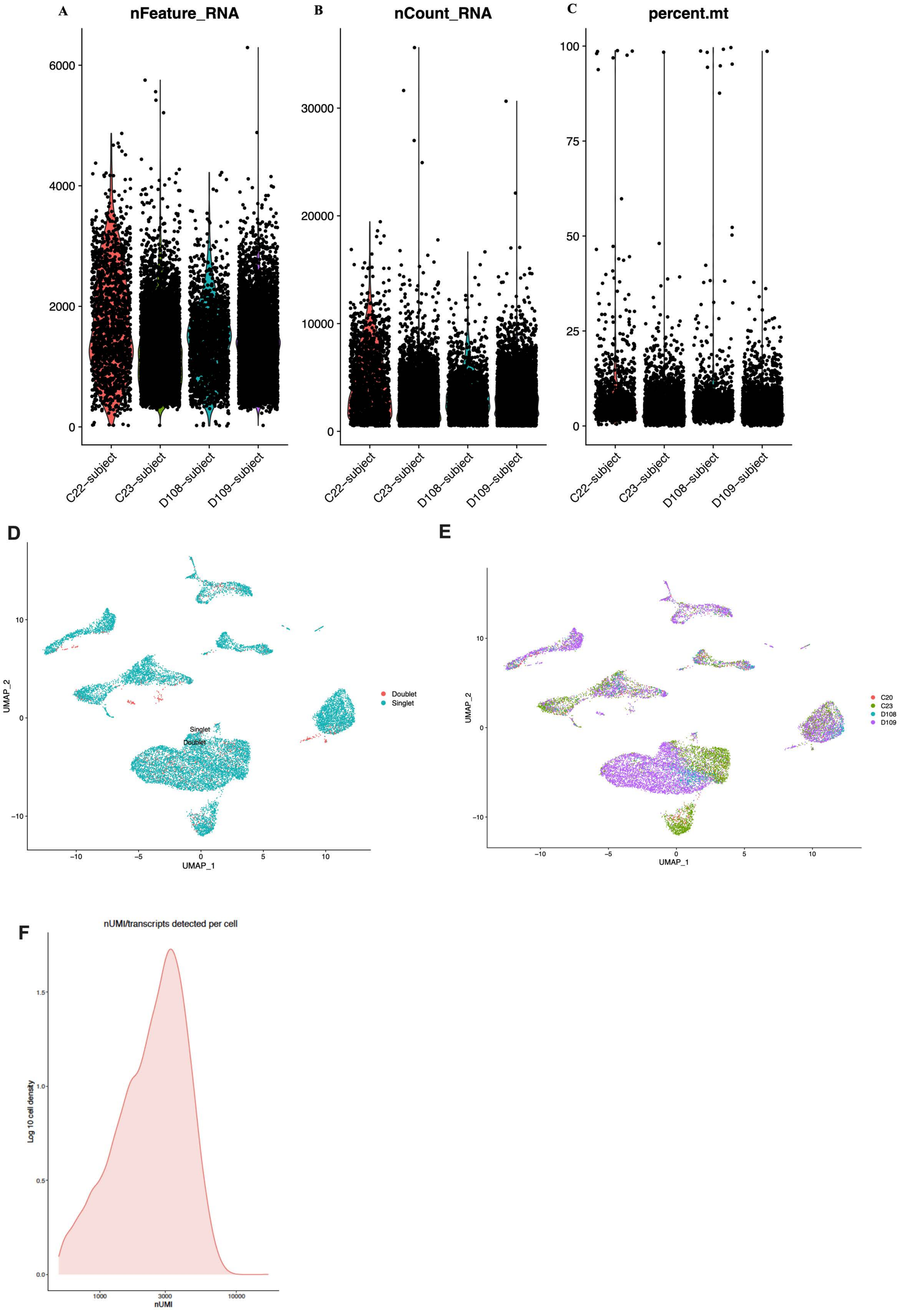
A
nFeature_RNA
B
nCount_RNA
C
percent.mt
C22–subject
C23–subject
D108–subject
D109–subject
D
UMAP_1
UMAP_2
Doublet
Singlet
E
C20
C23
D108
D109
F
nUMI/transcripts detected per cell
Log 10 cell density
nUMI

**Figure S6**

Quality control for scRNA-seq samples (n=4) including number genes identified (**A**), counts (**B**), percentage mitochondrial genes (**C**), identification of doublets (**D**) and donor contribution to each cell cluster (**E**).

Figure S7

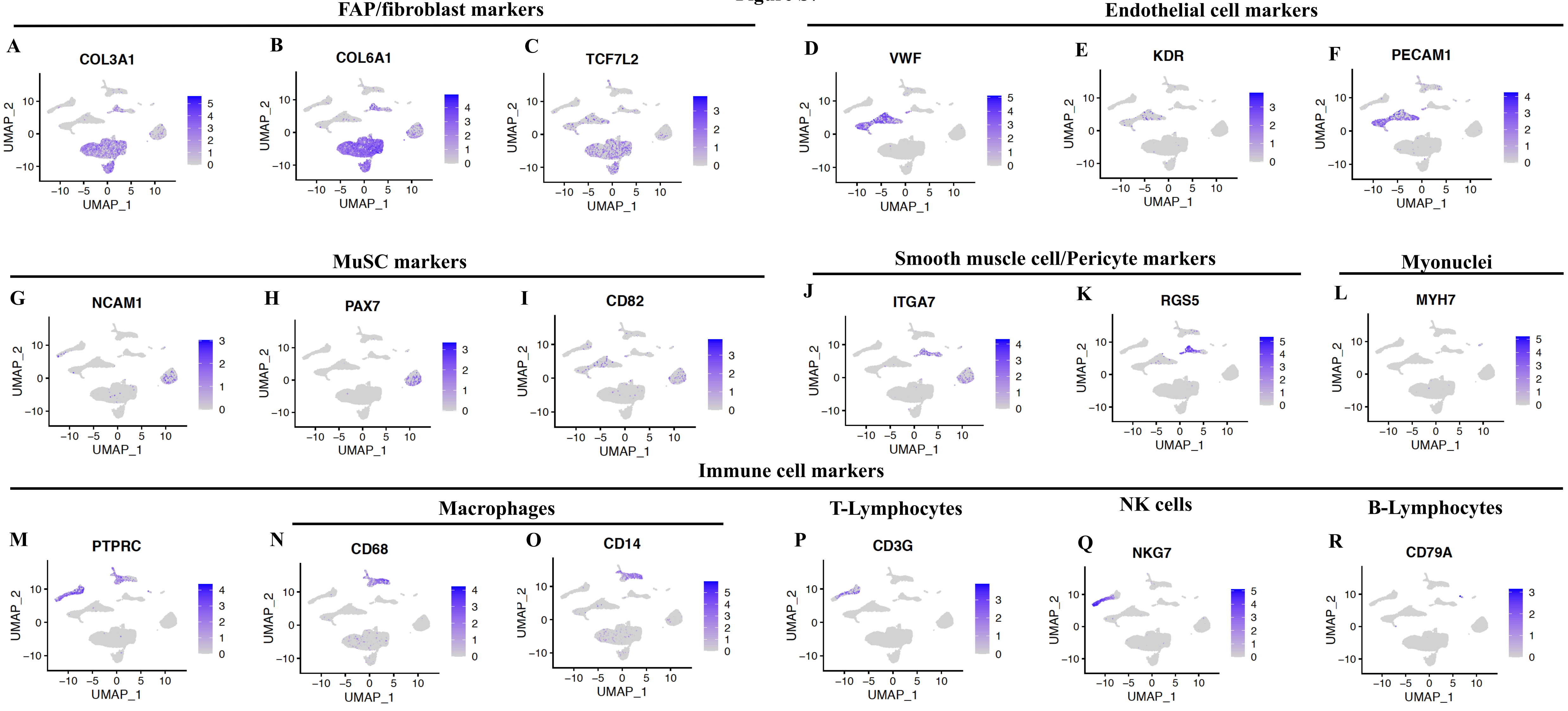
FAP/fibroblast markers
Endothelial cell markers
A
COL3A1
B
COL6A1
C
TCF7L2
D
VWF
E
KDR
F
PECAM1
MuSC markers
Smooth muscle cell/Pericyte markers
Myonuclei
G
NCAM1
H
PAX7
I
CD82
J
ITGA7
K
RGS5
L
MYH7
Immune cell markers
Macrophages
T-Lymphocytes
NK cells
B-Lymphocytes
M
PTPRC
N
CD68
O
CD14
P
CD3G
Q
NKG7
R
CD79A
UMAP_1
UMAP_2

**Figure S7**
Feature plots obtained from single cell RNA-seq (n=4) on human skeletal muscle displaying expression of genes/markers common for FAPs/fibroblasts: *COL3A1* (**A**), *COL6A1* (**B**), *TCF7L2*; Endothelial cells: (**C**) *VWF* (**D**), *KDR* (**E**), *PECAM1* (**F**); Muscle stem cell: *NCAM1* (**G**), *PAX7* (**H**), *CD82* (**I**); Smooth muscle cells/Pericytes: *ITGA7* (**J**), *RGS5* (**K**); Myonuclei: MYH7 (**L**); Immune cells: *PTPRC*/CD45 (**M**), *CD68* (**N**), *CD14* (**O**), *CD3G* (**P**), *NKG7* (**Q**), *CD79A* (**R**)

#### Figure S8

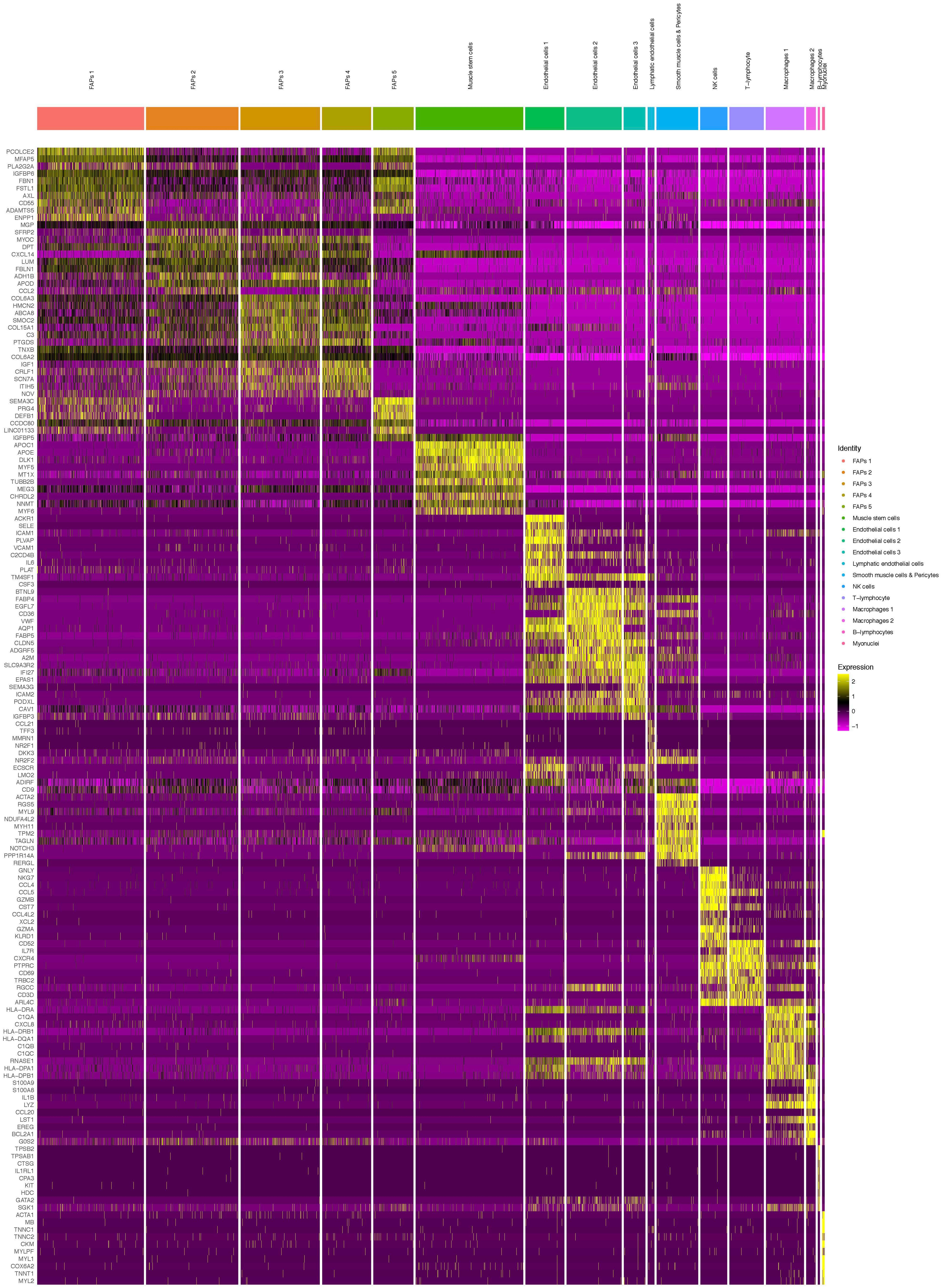
FAPs 1
FAPs 2
FAPs 3
FAPs 4
FAPs 5
Muscle stem cells
Endothelial cells 1
Endothelial cells 2
Endothelial cells 3
Lymphatic endothelial cells
Smooth muscle cells & Pericytes
NK cells
T−lymphocyte
Macrophages 1
Macrophages 2
B−lymphocytes
Myonuclei
PCOLCE2
MFAP5
PLA2G2A
IGFBP6
FBN1
FSTL1
AXL
CD55
ADAMTS5
ENPP1
MGP
SFRP2
MYOC
DPT
CXCL14
LUM
FBLN1
ADH1B
APOD
CCL2
COL6A3
HMCN2
ABCA8
SMOC2
COL15A1
C3
PTGDS
TNXB
COL6A2
IGF1
CRLF1
SCN7A
ITIH5
NOV
SEMA3C
PRG4
DEFB1
CCDC80
LINC01133
IGFBP5
APOC1
APOE
DLK1
MYF5
MT1X
TUBB2B
MEG3
CHRDL2
NNMT
MYF6
ACKR1
SELE
ICAM1
PLVAP
VCAM1
C2CD4B
IL6
PLAT
TM4SF1
CSF3
BTNL9
FABP4
EGFL7
CD36
VWF
AQP1
FABP5
CLDN5
ADGRF5
A2M
SLC9A3R2
IFI27
EPAS1
SEMA3G
ICAM2
PODXL
CAV1
IGFBP3
CCL21
TFF3
MMRN1
NR2F1
DKK3
NR2F2
ECSCR
LMO2
ADIRF
CD9
ACTA2
RGS5
MYL9
NDUFA4L2
MYH11
TPM2
TAGLN
NOTCH3
PPP1R14A
RERGL
GNLY
NKG7
CCL4
CCL5
GZMB
CST7
CCL4L2
XCL2
GZMA
KLRD1
CD52
IL7R
CXCR4
PTPRC
CD69
TRBC2
RGCC
CD3D
ARL4C
HLA−DRA
C1QA
CXCL8
HLA−DRB1
HLA−DQA1
C1QB
C1QC
RNASE1
HLA−DPA1
HLA−DPB1
S100A9
S100A8
IL1B
LYZ
CCL20
LST1
EREG
BCL2A1
G0S2
TPSB2
TPSAB1
CTSG
IL1RL1
CPA3
KIT
HDC
GATA2
SGK1
ACTA1
MB
TNNC1
TNNC2
CKM
MYLPF
MYL1
COX6A2
TNNT1
MYL2
Identity
FAPs 1
FAPs 2
FAPs 3
FAPs 4
FAPs 5
Muscle stem cells
Endothelial cells 1
Endothelial cells 2
Endothelial cells 3
Lymphatic endothelial cells
Smooth muscle cells & Pericytes
NK cells
T−lymphocyte
Macrophages 1
Macrophages 2
B−lymphocytes
Myonuclei
Expression
2
1
0
−1

**Figure S8**

Heat-map of top-ten differentially expressed genes in 17 populations of cells obtained from single cell RNA-seq (n=4) on human skeletal muscle.

#### Figure S9

#### GO: Biological processes from scRNA-seq

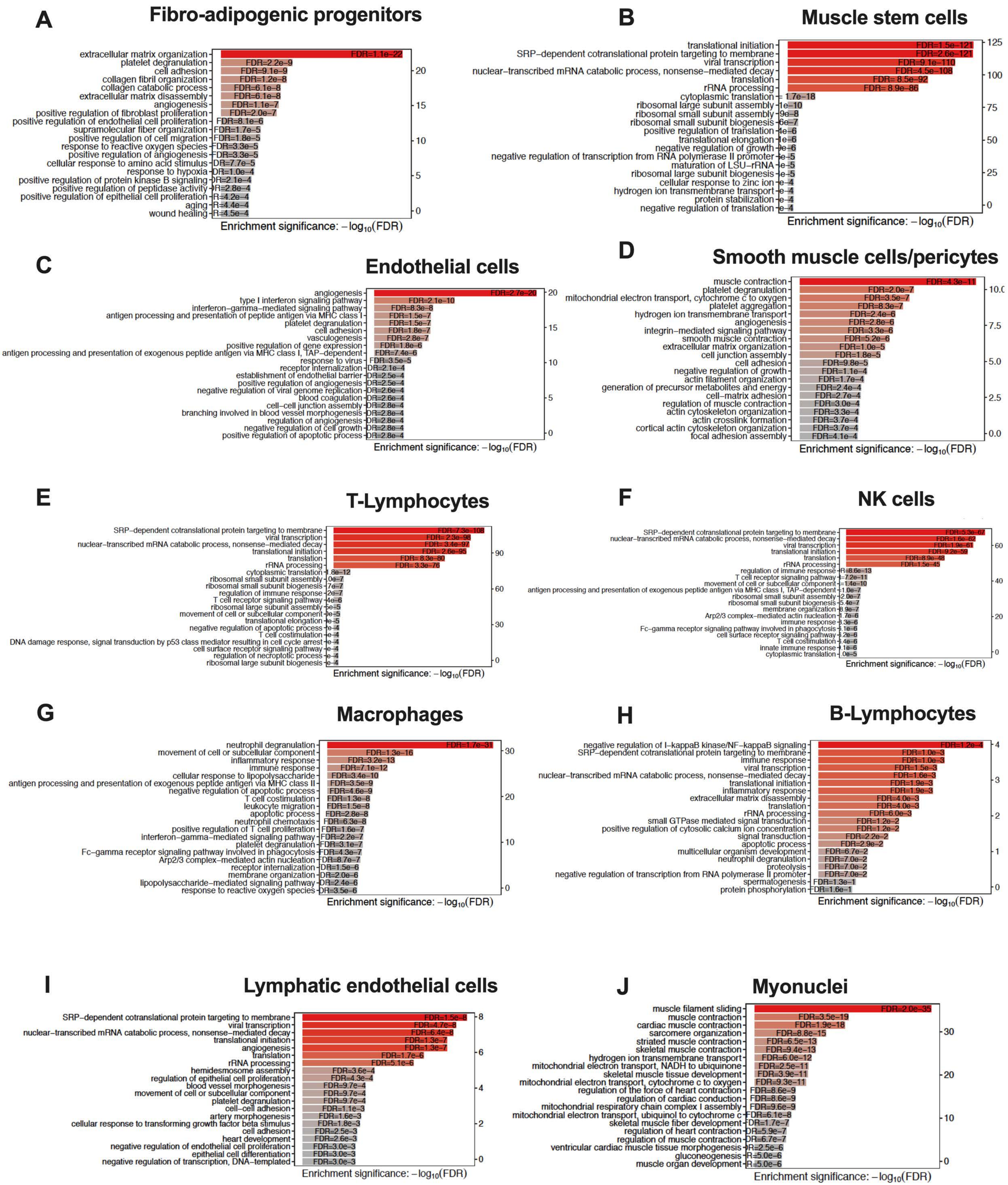

**Figure S9**

GO: Biological processes enriched in the ten major cell populations; Fibro-adipogenic progenitors (**A**), muscle stem cells (**B**), Endothelial cells (**C**), smooth muscle cells/pericytes (**D**), T-Lymphocytes (**E**), NK-cells (**F**), macrophages (**G**), B-Lymphocytes (**H**), Lymphatic Endothelial cells (**I**) and Myonuclei (**J**). Cells were obtained from single cell RNA-seq (n=4) on human skeletal muscle.

### Figure S10

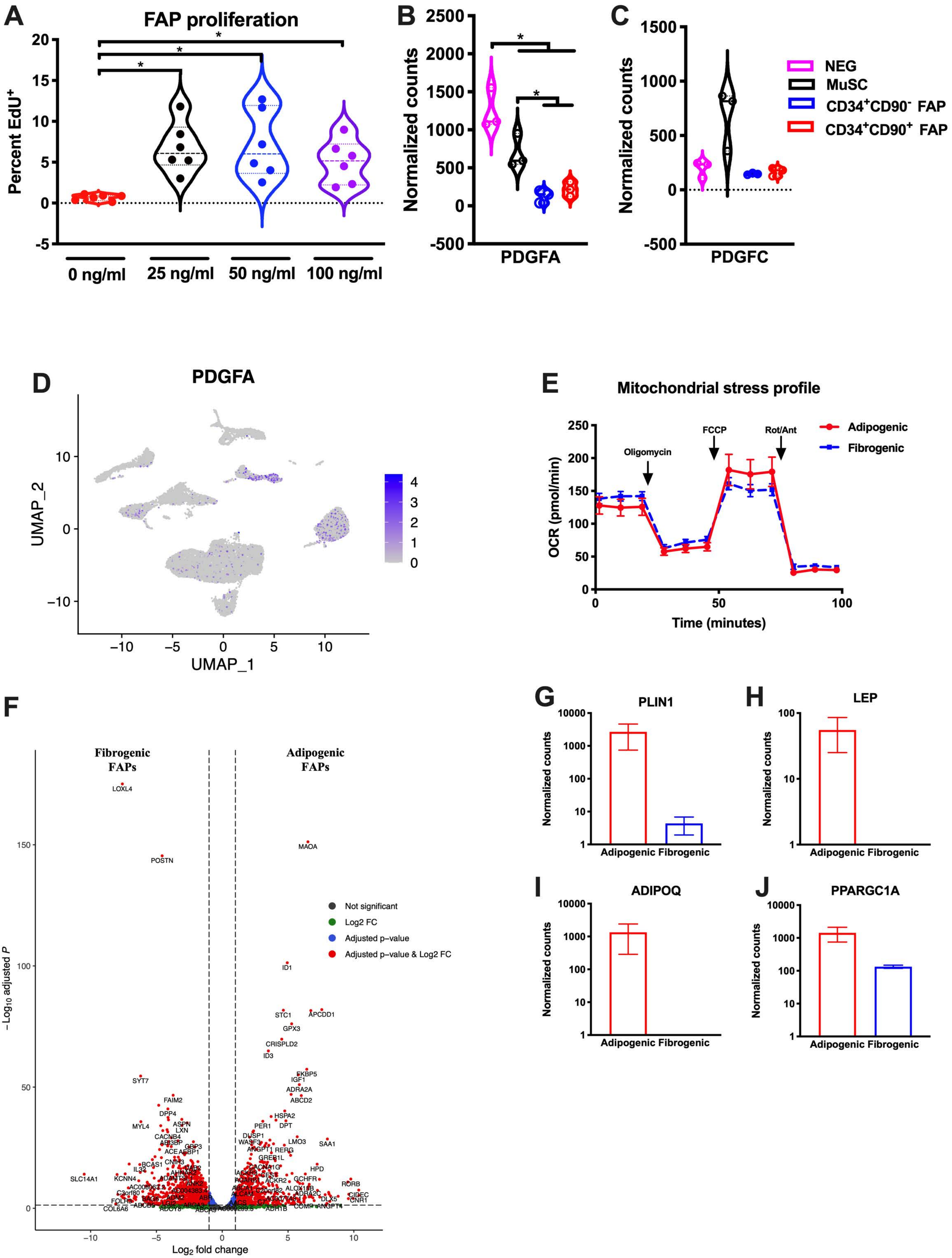

A
FAP proliferation
Percent EdU+
0 ng/ml
25 ng/ml
50 ng/ml
100 ng/ml
B
Normalized counts
PDGFA
C
PDGFC
NEG
MuSC
CD34+CD90- FAP
CD34+CD90+ FAP
D
PDGFA
UMAP_2
UMAP_1
E
Mitochondrial stress profile
Oligomycin
FCCP
Rot/Ant
Adipogenic
Fibrogenic
OCR (pmol/min)
Time (minutes)
F
Fibrogenic FAPs
Adipogenic FAPs
Not significant
Log2 FC
Adjusted p–value
Adjusted p–value & Log2 FC
−Log10 adjusted P
Log2 fold change
G
PLIN1
H
LEP
I
ADIPOQ
J
PPARGC1A
Adipogenic Fibrogenic

**Figure S10**

**A,** FACS isolated FAPs ($CD34^+CD90^+CD56^-CD31^-CD45^-$) stimulated with control (PBS) or 25, 50 100 ng/ml PDGF-AA for proliferation (EdU) in 24 hours (n=6 from 3 biological replicates); Normalized gene expression of *PDGFA* (**B**) and *PDGFC* (**C**) obtained from FACS sorted FAPs ($CD34^+CD90^-CD56^-CD31^-CD45^-$ and $CD34^+CD90^+CD56^-CD31^-CD45^-$), MuSCs ($CD56^+CD82^+CD34^-CD45^-CD31^-$) and mature skeletal muscle fragments/nuclei ($CD90^-CD56^-CD82^-CD34^-CD45^-CD31^-$); **D**, Feature plot of *PDGFA* gene expression in single-cell RNA-seq data; **E**, Mitochondrial profile (Oxygen consumption rate, pmol/min) of FAPs stimulated for 24h with adipogenic or fibrogenic media with sequential injections of Oligomycin, Carbonyl cyanide-*4*-(trifluoromethoxy) enylhydrazone (FCCP) and Rotenone/Antimycin (n=7, technical replicates); **F**, Volcano plot of differentially expressed genes in FAPs stimulated towards fibrogenesis (with PDGF-AA) or adipogenesis for six days (n=3); Normalized gene expression of key adipogenesis genes from RNA-seq data from FAPs stimulated for six days towards fibrogenesis and adipogenesis (n=3) including Perilipin-1 (*PLIN1*, **G**), Leptin (*LEP*, **H**), Adiponectin (*ADIPOQ*, **I**) and PGC1α (*PPARGC1A*, **J**). Significant difference denoted by $*p<0.05$.

Figure S11

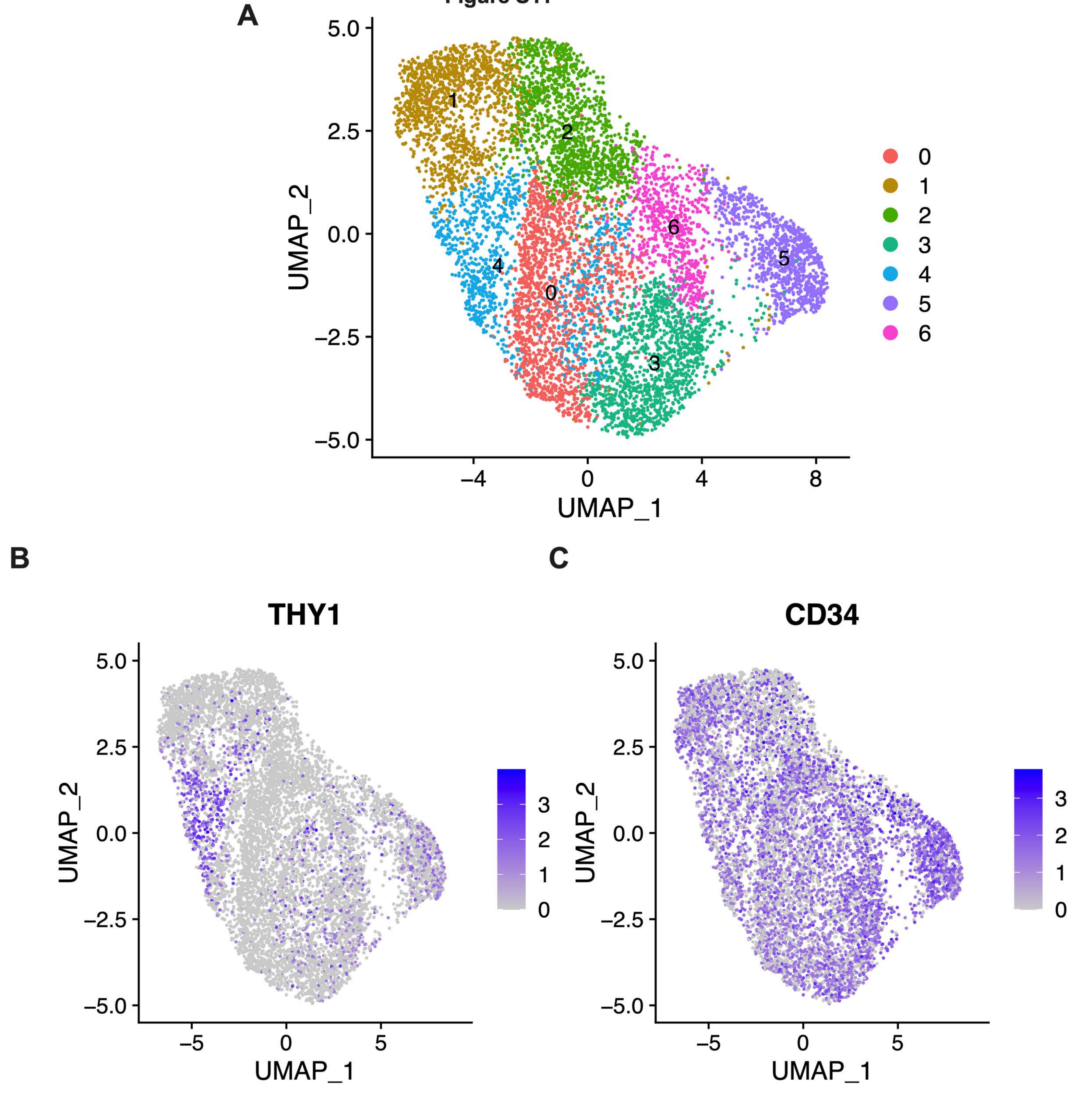

**Figure S11**
**A,** UMAP plot of FAPs revealing seven subpopulations (0-6) within the FAP population obtained from single-cell RNA-seq on whole human skeletal muscle (n=4); **B**, Feature plot of *THY1*/CD90 expression in the FAP subpopulations; **C**, Feature plot of *CD34* expression in the FAP subpopulations.

Figure S12

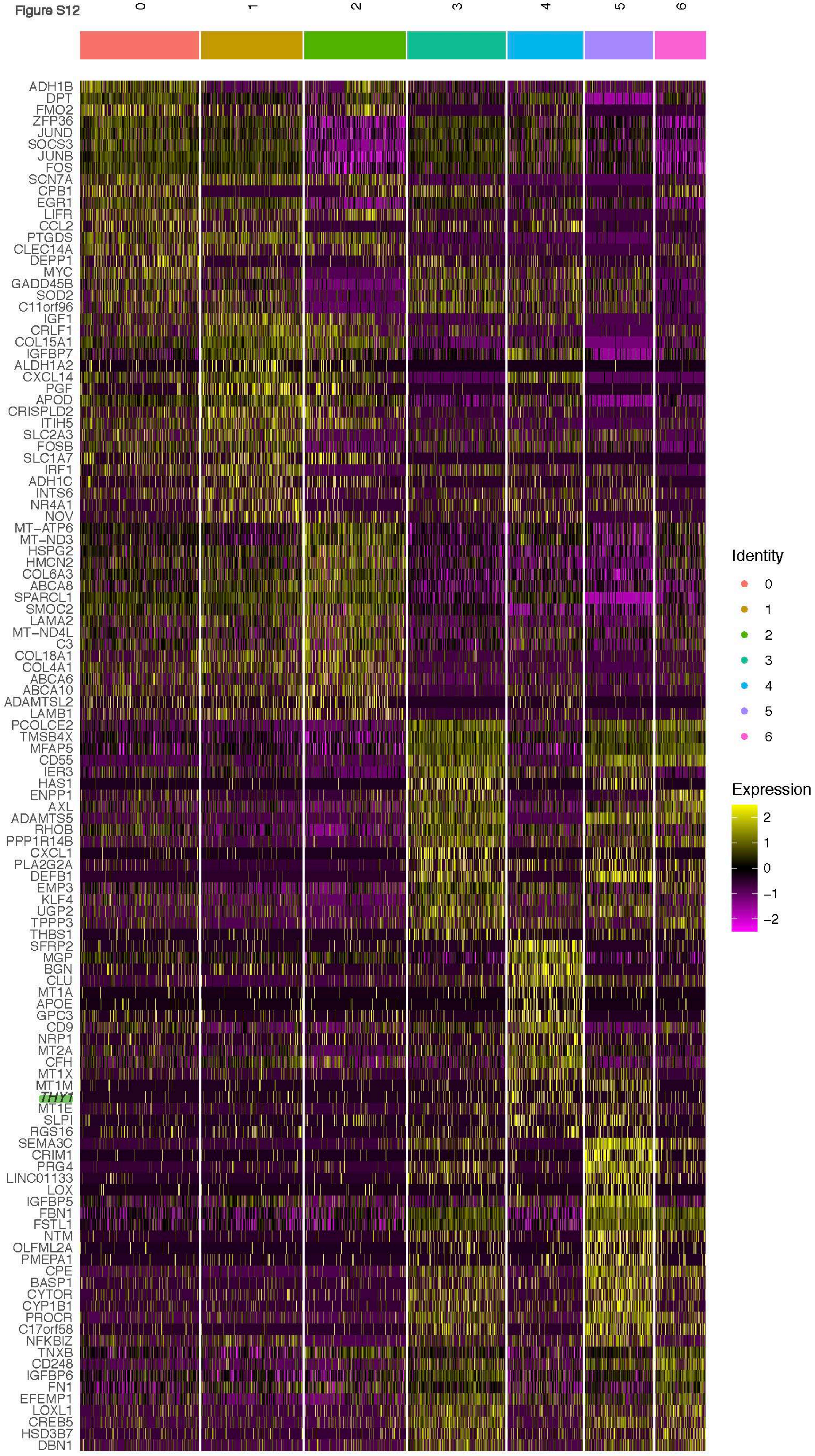
0
1
2
3
4
5
6
ADH1B
DPT
FMO2
ZFP36
JUND
SOCS3
JUNB
FOS
SCN7A
CPB1
EGR1
LIFR
CCL2
PTGDS
CLEC14A
DEPP1
MYC
GADD45B
SOD2
C11orf96
IGF1
CRLF1
COL15A1
IGFBP7
ALDH1A2
CXCL14
PGF
APOD
CRISPLD2
ITIH5
SLC2A3
FOSB
SLC1A7
IRF1
ADH1C
INTS6
NR4A1
NOV
MT−ATP6
MT−ND3
HSPG2
HMCN2
COL6A3
ABCA8
SPARCL1
SMOC2
LAMA2
MT−ND4L
C3
COL18A1
COL4A1
ABCA6
ABCA10
ADAMTSL2
LAMB1
PCOLCE2
TMSB4X
MFAP5
CD55
IER3
HAS1
ENPP1
AXL
ADAMTS5
RHOB
PPP1R14B
CXCL1
PLA2G2A
DEFB1
EMP3
KLF4
UGP2
TPPP3
THBS1
SFRP2
MGP
BGN
CLU
MT1A
APOE
GPC3
CD9
NRP1
MT2A
CFH
MT1X
MT1M
THY1
MT1E
SLPI
RGS16
SEMA3C
CRIM1
PRG4
LINC01133
LOX
IGFBP5
FBN1
FSTL1
NTM
OLFML2A
PMEPA1
CPE
BASP1
CYTOR
CYP1B1
PROCR
C17orf58
NFKBIZ
TNXB
CD248
IGFBP6
FN1
EFEMP1
LOXL1
CREB5
HSD3B7
DBN1
Identity
0
1
2
3
4
5
6
Expression
2
1
0
−1
−2

**Figure S12**

Heat-map of top-twenty differentially expressed genes in the seven identified subpopulations obtained from single cell RNA-seq on whole human skeletal muscle (n=4), *THY1*/CD90 is highlighted.

### Figure S13

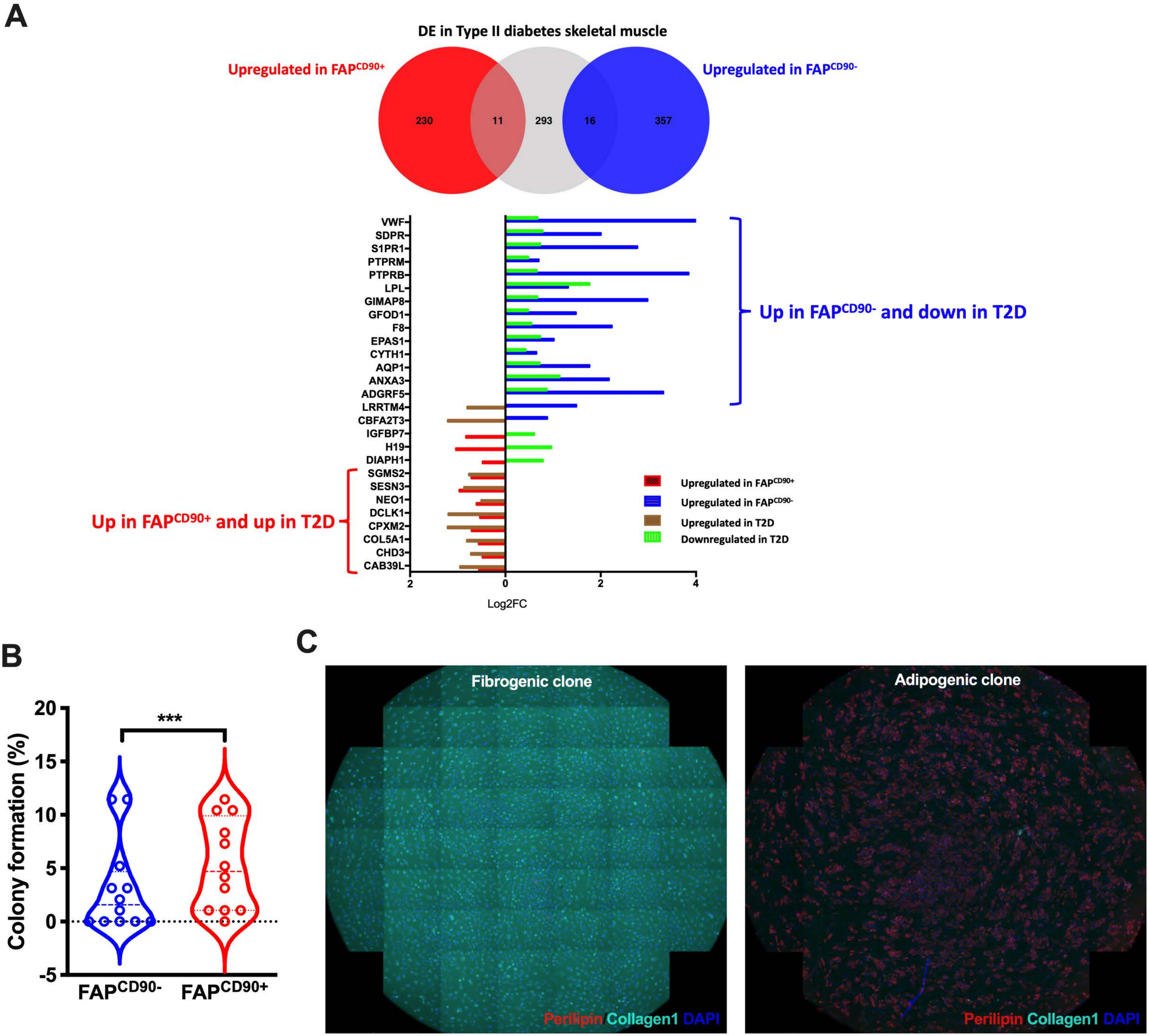

A
DE in Type II diabetes skeletal muscle
Upregulated in FAP^CD90+
Upregulated in FAP^CD90-
230
11
293
16
357
VWF
SDPR
S1PR1
PTPRM
PTPRB
LPL
GIMAP8
GFOD1
F8
EPAS1
CYTH1
AQP1
ANXA3
ADGRF5
LRRTM4
CBFA2T3
IGFBP7
H19
DIAPH1
SGMS2
SESN3
NEO1
DCLK1
CPXM2
COL5A1
CHD3
CAB39L
Up in FAP^CD90- and down in T2D
Up in FAP^CD90+ and up in T2D
Upregulated in FAP^CD90+
Upregulated in FAP^CD90-
Upregulated in T2D
Downregulated in T2D
2
0
2
4
Log2FC
B
Colony formation (%)
20
15
10
5
0
-5
***
FAP^CD90-
FAP^CD90+
C
Fibrogenic clone
Adipogenic clone
Perilipin Collagen1 DAPI
Perilipin Collagen1 DAPI

**Figure S13**

**A,** Correlation of differentially expressed genes in insulin dependent type 2 diabetes (itT2D) patients with upregulated genes in $FAP^{CD90-}$ and $FAP^{CD90+}$, respectively; **B**, Number of colonies generated from single freshly FACS isolated $FAP^{CD90-}$ and $FAP^{CD90+}$ (n=12); **C**, Stitched images from an entire well obtained from single-cell sorted FAPs displaying either fibrogenic (Collagen-1, cyan) or adipogenic (Perilipin-1, red) differentiation. Significant difference denoted by ***p<0.001.

Figure S14

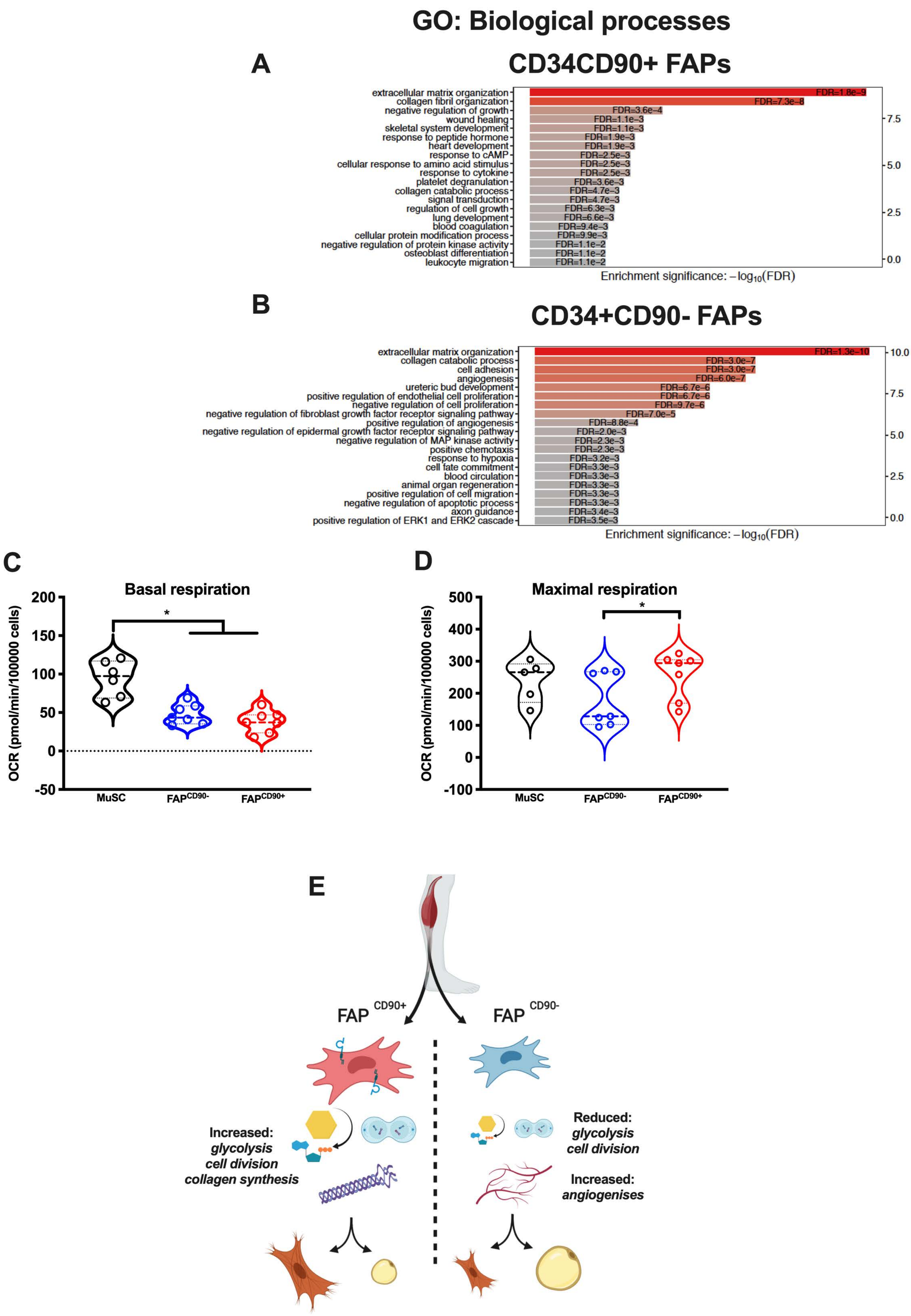

GO: Biological processes
A
CD34CD90+ FAPs
extracellular matrix organization
collagen fibril organization
negative regulation of growth
wound healing
skeletal system development
response to peptide hormone
heart development
response to cAMP
cellular response to amino acid stimulus
response to cytokine
platelet degranulation
collagen catabolic process
signal transduction
regulation of cell growth
lung development
blood coagulation
cellular protein modification process
negative regulation of protein kinase activity
osteoblast differentiation
leukocyte migration
FDR=1.8e-9
FDR=7.3e-8
FDR=3.6e-4
FDR=1.1e-3
FDR=1.1e-3
FDR=1.9e-3
FDR=1.9e-3
FDR=2.5e-3
FDR=2.5e-3
FDR=2.5e-3
FDR=3.6e-3
FDR=4.7e-3
FDR=4.7e-3
FDR=6.3e-3
FDR=6.6e-3
FDR=9.4e-3
FDR=9.9e-3
FDR=1.1e-2
FDR=1.1e-2
FDR=1.1e-2
Enrichment significance: $-\log_{10}(FDR)$
B
CD34+CD90- FAPs
extracellular matrix organization
collagen catabolic process
cell adhesion
angiogenesis
ureteric bud development
positive regulation of endothelial cell proliferation
negative regulation of cell proliferation
negative regulation of fibroblast growth factor receptor signaling pathway
positive regulation of angiogenesis
negative regulation of epidermal growth factor receptor signaling pathway
negative regulation of MAP kinase activity
positive chemotaxis
response to hypoxia
cell fate commitment
blood circulation
animal organ regeneration
positive regulation of cell migration
negative regulation of apoptotic process
axon guidance
positive regulation of ERK1 and ERK2 cascade
FDR=1.3e-10
FDR=3.0e-7
FDR=3.0e-7
FDR=6.0e-7
FDR=6.7e-6
FDR=6.7e-6
FDR=9.7e-6
FDR=7.0e-5
FDR=8.8e-4
FDR=2.0e-3
FDR=2.3e-3
FDR=2.3e-3
FDR=3.2e-3
FDR=3.3e-3
FDR=3.3e-3
FDR=3.3e-3
FDR=3.3e-3
FDR=3.3e-3
FDR=3.4e-3
FDR=3.5e-3
Enrichment significance: $-\log_{10}(FDR)$
C
Basal respiration
OCR (pmol/min/100000 cells)
MuSC
FAP^CD90-
FAP^CD90+
D
Maximal respiration
OCR (pmol/min/100000 cells)
MuSC
FAP^CD90-
FAP^CD90+
E
FAP CD90+
FAP CD90-
Increased:
glycolysis
cell division
collagen synthesis
Reduced:
glycolysis
cell division
Increased:
angiogenises

**Figure S14**

GO: Biological processes enriched in FAP subpopulations obtained from single-cell RNA-seq on whole human skeletal muscle (n=4). Subpopulation for enrichment analysis on biological processes were divided in (**A**) $CD34^{+}CD90^{+}$ FAPs and (**B**) $CD34^{+}CD90^{-}$ FAPs; **C**, Basal normalized Oxygen consumption rate (OCR, pmol/min) of freshly sorted Muscle stem cells (MuSCs), $FAP^{CD90-}$ and $FAP^{CD90+}$ with sequential injections of Oligomycin, Carbonyl cyanide-*4*-(trifluoromethoxy) phenylhydrazone (FCCP) and Rotenone/Antimycin (n=7); **D**, Maximal normalized respiration (pmol/min) of MuSCs, $FAP^{CD90-}$ and $FAP^{CD90+}$ (n=7); **E**, Schematic presentation of the $FAP^{CD90+}$ and $FAP^{CD90-}$ phenotypes. Significant difference denoted by $^{*}p<0.05$

**Figure S15**

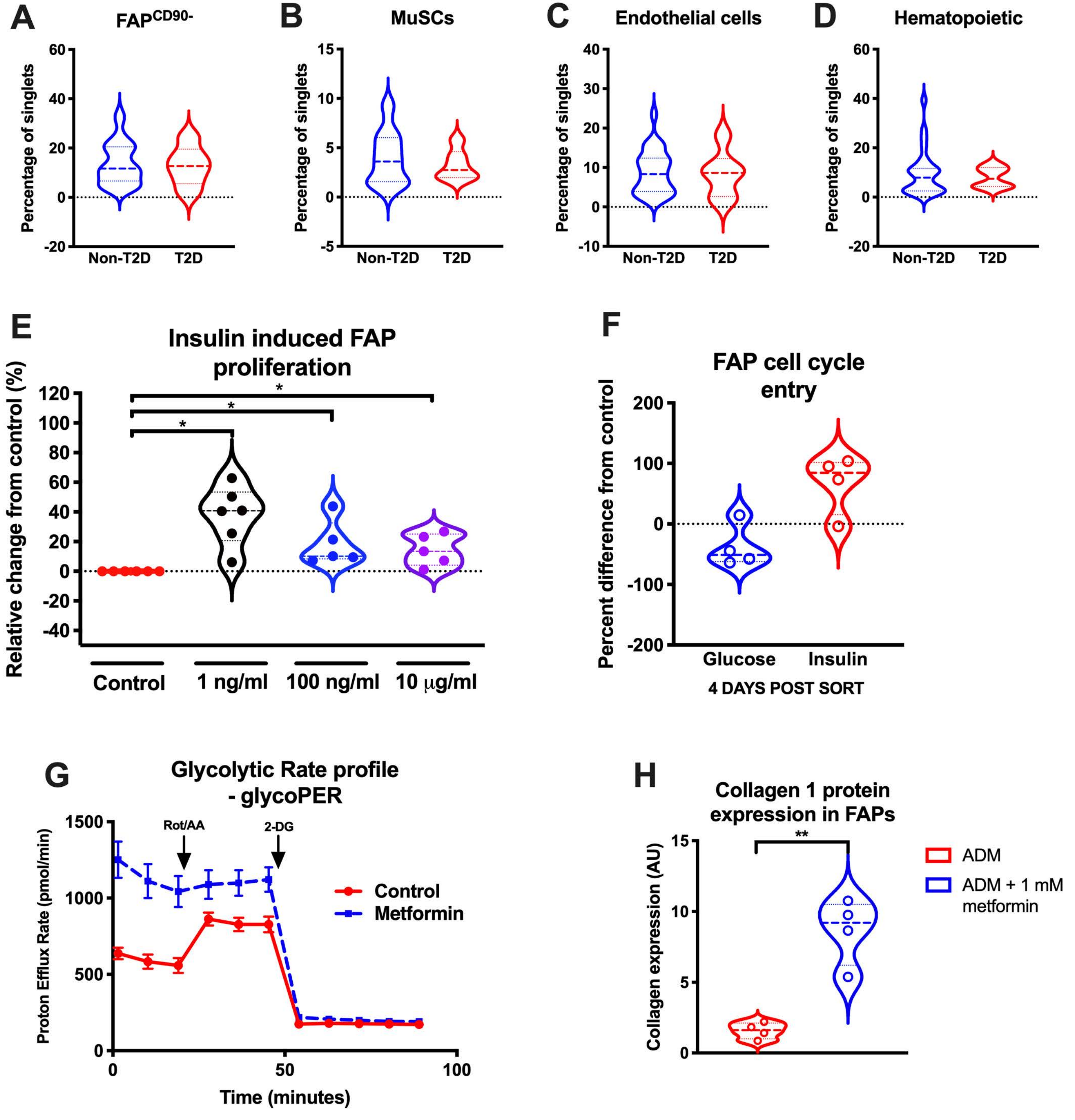
A
FAP$^{CD90-}$
B
MuSCs
C
Endothelial cells
D
Hematopoietic
Percentage of singlets
Non-T2D
T2D
E
Insulin induced FAP proliferation
Relative change from control (%)
Control
1 ng/ml
100 ng/ml
10 μg/ml
F
FAP cell cycle entry
Percent difference from control
Glucose
Insulin
4 DAYS POST SORT
G
Glycolytic Rate profile - glycoPER
Proton Efflux Rate (pmol/min)
Rot/AA
2-DG
Control
Metformin
Time (minutes)
H
Collagen 1 protein expression in FAPs
Collagen expression (AU)
ADM
ADM + 1 mM metformin

**Figure S15**

**A,** $FAP^{CD90-}$ content (percentage of single cells, %) in patients without type 2 diabetes (Non-T2D, n=27) and with type 2 diabetes (T2D, n=6); **B**, Muscle stem cell (MuSC) content (percentage of single cells, %) in Non-T2D and T2D patients; **C**, Endothelial cell content (percentage of single cells, %) in Non-T2D and T2D patients; **D** Hematopoietic cell content (percentage of single cells, %) in Non-T2D and T2D patients; **E**, FACS isolated FAPs ($CD34^{+}CD90^{+}CD56^{-}CD31^{-}CD45^{-}$) stimulated (expressed as percentage from control, %) with control (PBS) or 1 ng/ml, 100 ng/ml and 10 ug/ml recombinant human insulin for proliferation (EdU) for 24 hours (n=6 from 3 biological replicates); **F**, Cell cycle entry (EdU incorporation) 96 hours post FACS isolation of FAPs ($CD34^{+}CD90^{+}CD56^{-}CD31^{-}CD45^{-}$) stimulated with Glucose (low (1g/L) versus high (4.5g/L)) or insulin (PBS versus 1ng/ml); **G**, Glycolytic profile (glycolytic proton efflux rate; glycoPER, pmol/min per $6x10^4$ cells) on $FAP^{CD90+}$ exposed to control (PBS) or 1mM Metformin for 24h prior to bioenergetic analysis. Analysis was performed with a basal period followed by sequential injections of Rotenone/Antimycin and 2-deoxy-glucose (n=5-6, technical replicates); **H**, Collagen-1 protein expression (positive area per cell) in FAPs ($CD34^{+}CD90^{+}CD56^{-}CD31^{-}CD45^{-}$) stimulated towards adipogenesis for six days with control (PBS) or Metformin (1mM).
